# Skeletal muscle aging drives sex- and compartment-specific protein insolubility that is modifiable by proteostasis-directed treatments

**DOI:** 10.64898/2026.10.29.762839

**Authors:** P.H.C. Mesquita, T. Goyal, X. Ren, J. Papinska, M.E. Taylor, T.J. Lundgren, Z.R. Hettinger, B.F. Miller

**Author notes:** Address correspondence to Benjamin F. Miller, Aging & Metabolism Research Program, Oklahoma Medical Research Foundation, MS 21, 825 NE 13th St, Oklahoma City, OK 73104, USA. Co-Senior authors (Z.R. Hettinger, B.F. Miller).

## Abstract

Protein aggregation and altered protein solubility indicate proteostatic decline, yet their roles in skeletal muscle aging remain poorly understood. We combined light and electron microscopy, quantitative proteomics, and stable-isotope labeling of skeletal muscle to investigate age-associated changes in protein insolubility, and their responses to exercise and rapamycin. Aged muscle accumulated distinct aggresome and amyloid-positive aggregates despite unchanged bulk insoluble protein content. Instead, aging altered the abundance and soluble–insoluble shifts of lower-abundance proteins across cellular components, particularly mitochondrial, nuclear and extracellular matrix proteins, with greater changes in females. These changes were not broadly explained by intrinsic protein aggregation propensity. Exercise and rapamycin partially reversed age-associated changes in abundance and partitioning without increasing protein synthesis. Together, these findings highlight sex- and cellular component-dependent changes in protein insolubility as important, modifiable aspects of skeletal muscle proteostasis, and indicate that proteostatic maintenance may be a more important target than anabolism for muscle aging.

## INTRODUCTION

Muscle mass and function decline with aging, which accelerates the loss of independence and increases morbidity and mortality. Many therapeutic approaches have focused on activating mTOR to increase protein synthesis and muscle mass because of the concept that aged muscle has an “anabolic resistance” where there is a blunted increase in protein synthesis in response to anabolic stimuli. However, these therapies have largely failed to improve muscle mass, strength, or physical function in older adults ^1–3^. We and others have previously shown that protein synthesis rates are not slower, but rather are greater in old compared to adult muscle ^4–6^. Moreover, long-term inhibition of mTOR activity preserves muscle function with age ^7,8^. In addition, the magnitude of age-related declines in strength and physical performance exceed that of the loss of muscle mass ^9^, indicating that factors besides muscle mass also contribute to the loss of muscle quality. Collectively, these studies indicate that therapeutic approaches should move beyond targeting anabolic resistance.

Proteostasis encompasses the coordinated synthesis, folding, assembly, trafficking, and degradation of proteins ^10–12^. Proteostatic mechanisms decline with advancing age, promoting protein misfolding and the accumulation of protein aggregates (PA) across multiple cell and tissue types ^13–15^. Proteins live on the edge of solubility ^16^ and although insolubility can be a means of organizing protein structures, increased protein insolubility and aggregation stresses protein quality control mechanisms, creating a feed-forward cycle of proteostatic stress. Insoluble proteins and PA are defining features of several neurodegenerative diseases and have been extensively studied in neuronal tissues ^17,18^. In contrast, proteostatic mechanisms and the molecular features of age-associated PA and altered protein solubility in skeletal muscle remain poorly understood.

Much of the existing knowledge of PA in skeletal muscle comes from studies of muscular dystrophies ^19^, protein aggregate myopathies ^19^, or studies from model organisms such as *C. elegans* ^20^ and Drosophila ^21^. Some studies show the age-related accumulation of PA in the skeletal muscle of mice ^22^ and humans ^23^ and the remodeling of the insoluble proteome in male mice ^24^. However, the broader subcellular organization and molecular composition of the aged skeletal muscle insoluble proteome, including sex-specific remodeling, remain unknown.

Highly abundant proteins are the major contributors by mass to the accumulation of PA because their overproduction can push proteins beyond a critical solubility limit ^25^. In skeletal muscle, myofibrillar proteins are by far the most abundant proteins, which makes them likely contributors to PA. Consistent with this idea, we have previously shown that in aged rats undergoing reloading after disuse atrophy, 16 of the top 20 most abundant insoluble proteins were myofibrillar ^5^. Whether aging selectively alters the composition of the skeletal muscle insoluble proteome beyond these dominant structural proteins remains unclear.

In the present study, we combined light and electron microscopy, biochemical fractionation, proteomics, and stable isotope labeling to define how aging affects protein aggregation and the insoluble proteome of skeletal muscle. We further tested whether these changes differed between females and males and whether they were modified by voluntary exercise or pharmacological mTOR inhibition. We show that although aging does not increase bulk insoluble protein content in skeletal muscle, there is an accumulation of distinct aggregate structures with sex- and subcellular-specific changes in the insoluble proteome that are partially reversed by exercise and mTOR inhibition. These findings indicate that the maintenance of subcellular proteostasis could be a more important target than anabolism for maintaining muscle quality with age.

## RESULTS

### Aging promotes the accumulation of distinct protein aggregates in skeletal muscle

To test our hypothesis that aging increases protein aggregates in muscle, we used high resolution light microscopy and conformation-dependent fluorescent dyes, Proteostat™ and AmyTracker™, coupled with Transmission Electron Microscopy (TEM). Proteostat™ detects aggregate proteins associated with aggresomes and inclusion bodies^26^, whereas AmyTracker™, a fluorescent Congo Red derivative, preferentially identifies more stable and insoluble amyloid aggregates ^27^. Aggresomes (Proteostat™-positive) were detected in both adult and aged tibialis anterior (TA) muscle but were infrequent in adults and more prominent in aged muscle, where they appeared as large structures with luminal morphology (**Fig. 1a**). TEM further showed large structures in aged muscle with features consistent with aggresome-like morphology ^26,28^ (**Fig. 1b; Extended Data Fig. 1a**). TEM also showed the well-described sarcoplasmic reticulum (SR)-derived tubular aggregates, characterized by ordered arrays of membrane tubules as previously described, albeit at sizes nowhere close to the size of previously unreported and large aggresomes ^29,30^ (**Ext Data Fig. 1a,b**). Spherical AmyTracker™-positive aggregates were observed discretely compartmentalized to the sarcolemma in adult muscle, whereas aged muscle showed disrupted sarcolemmal architecture with apparent release of aggregates into the extracellular space (**Fig. 1c**). TEM further showed large and compositionally homogeneous ultrastructure of subsarcolemmal failed autophagolysosomes in aged muscle (**Fig. 1d**). Compared with adults, aged muscle showed a 1.91-fold increase in amyloid aggregate number and a 1.84-fold increase in aggregate-occupied volume (**Fig. 1e,f**), without detectable differences in aggregate morphology (**Supplementary Fig. 1**). These data suggest that amyloid aggregates can form under adult conditions, but are in significantly higher frequency in aged compared to adult muscles. Together, these findings show the age-associated accumulation of aggresome-like and amyloid-positive structures beyond the well-characterized tubular aggregates, highlighting distinct manifestations of proteostatic failure in aging skeletal muscle. To our knowledge, this is the first reporting of aggresomes and amyloid+ aggregates in aging murine muscle cells under normal physiological aging conditions.

**Fig. 1.**
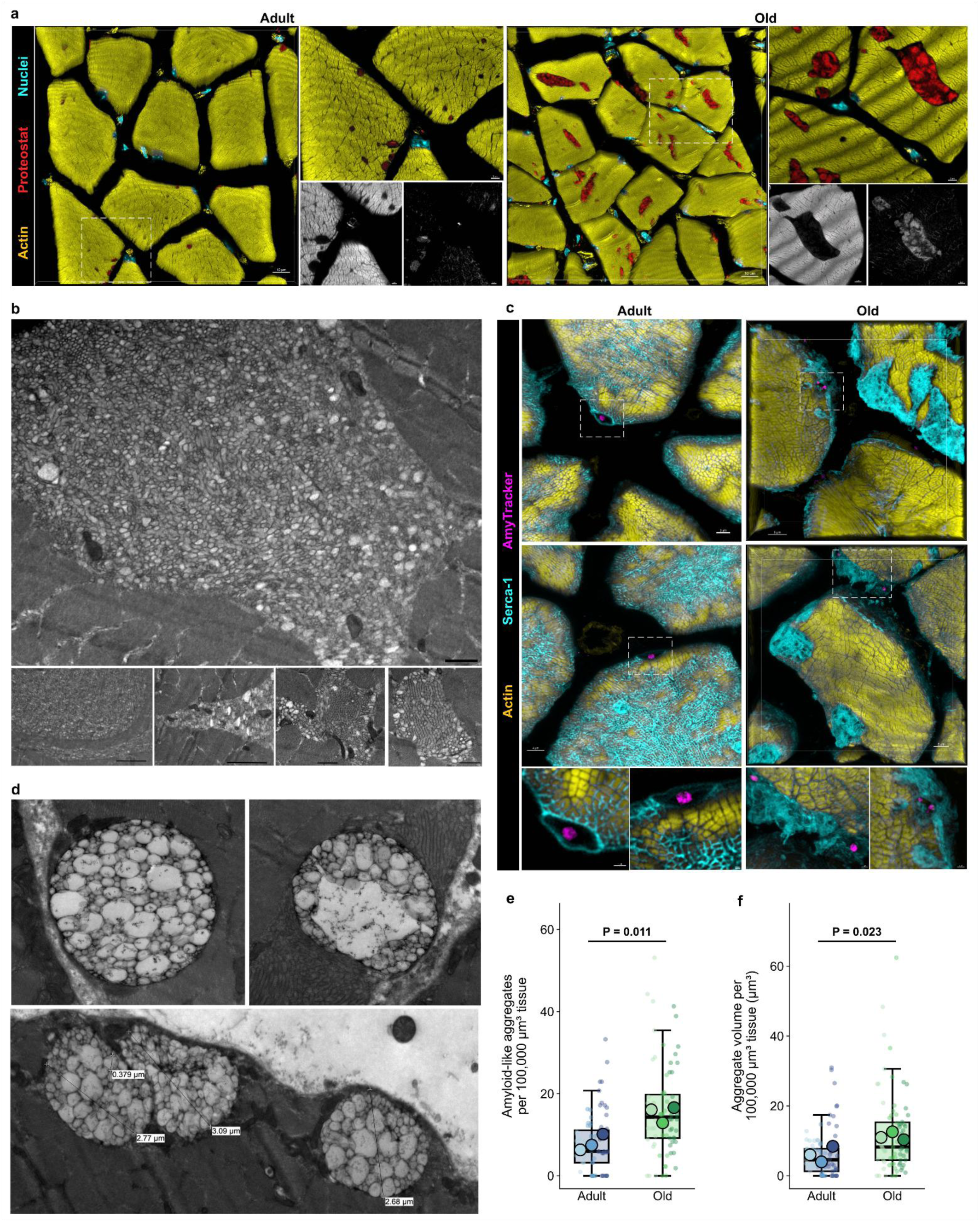
Aging promotes the accumulation of distinct protein aggregates in skeletal muscle. **a**, Subcellular fluorescence imaging of Proteostat™ (red), Phalloidin/Actin (yellow), and dsDNA/nuclei (cyan) in adult (7 month) and old (24 month) tibialis anterior (TA) muscles. Scale bar is 10, 3, and 2 microns for 3D, 2D, and 1D images, respectively. 1D grayscale insets highlight Actin (left) and Proteostat™ (right) signals. **b,** Transmission electron microscopy (TEM) micrograph of old (24 month) TA muscle of aggresomes viewed cross-sectionally (scale bar is 1 micron) and longitudinally (bottom row of images). Insets (white dotted boxes) and anatomical landmarks (yellow asterisks) of top left image of cross-sectionally oriented muscle cell are denoted for adjacent higher resolution views in the adjacent panel images. Micrographs of longitudinally oriented aggresome too large for the camera’s single field of view are positioned adjacently to highlight longitudinal aggresome morphology. **c,** Subcellular fluorescence imaging of AmyTracker™ (magenta), Phalloidin/Actin (yellow), and Serca-1 (cyan) in adult (7 month) and old (24 month) TA muscles. Insets (white dotted boxes) of 3D images highlight 2D magnified views shown below 3D overviews. **d,** TEM micrographs of old (24 month) TA muscle of amyloid aggregates viewed cross-sectionally (top row) and transverse (bottom image). Aggregate diameters are shown. Superplots of amyloid aggregate abundance by count **e,** and volume **f,** between adult (7 month) and old (24 month) TA muscles. Large circles indicate biological replicates. Small translucent circles correspond to technical replicates per biological replicate. Horizontal line corresponds to average per biological sample and error bars correspond to 95% confidence interval. Welch’s t-test. \**p*<0.05. N=3 mice/age.

### Aging induces sex-divergent proteostatic remodeling without bulk protein insolubilization

Following the identification of aggregate structures in aged skeletal muscle, we tested the hypothesis that this aggregated phenotype would be accompanied by increased insoluble protein abundance, particularly of contractile proteins. We fractionated gastrocnemius muscle into detergent-soluble and detergent-insoluble pools and analyzed both fractions by proteomics. Despite the prominent aggregate-like phenotype of aged muscle detected with imaging, Coomassie staining showed no age-related changes in total soluble, insoluble, or insoluble-to-soluble protein ratio (**Fig. 2a**). Similarly, stable-isotope labeling showed no age-related differences in bulk protein synthesis rates of the myofibrillar, mitochondrial, or cytosolic fractions (**Extended Data Fig. 2a-c**). We leveraged our tracers to test if newly synthesized proteins were less likely to be found in the insoluble protein fraction in aged muscle. Although unchanged with age, the lower fraction new in the insoluble fraction indicates a slower turnover of the insoluble fraction compared to the soluble fraction (**Extended Data Fig. 2d**). Importantly, our stable-isotope approaches used both sexes together prior to determining sex differences.

**Fig. 2.**
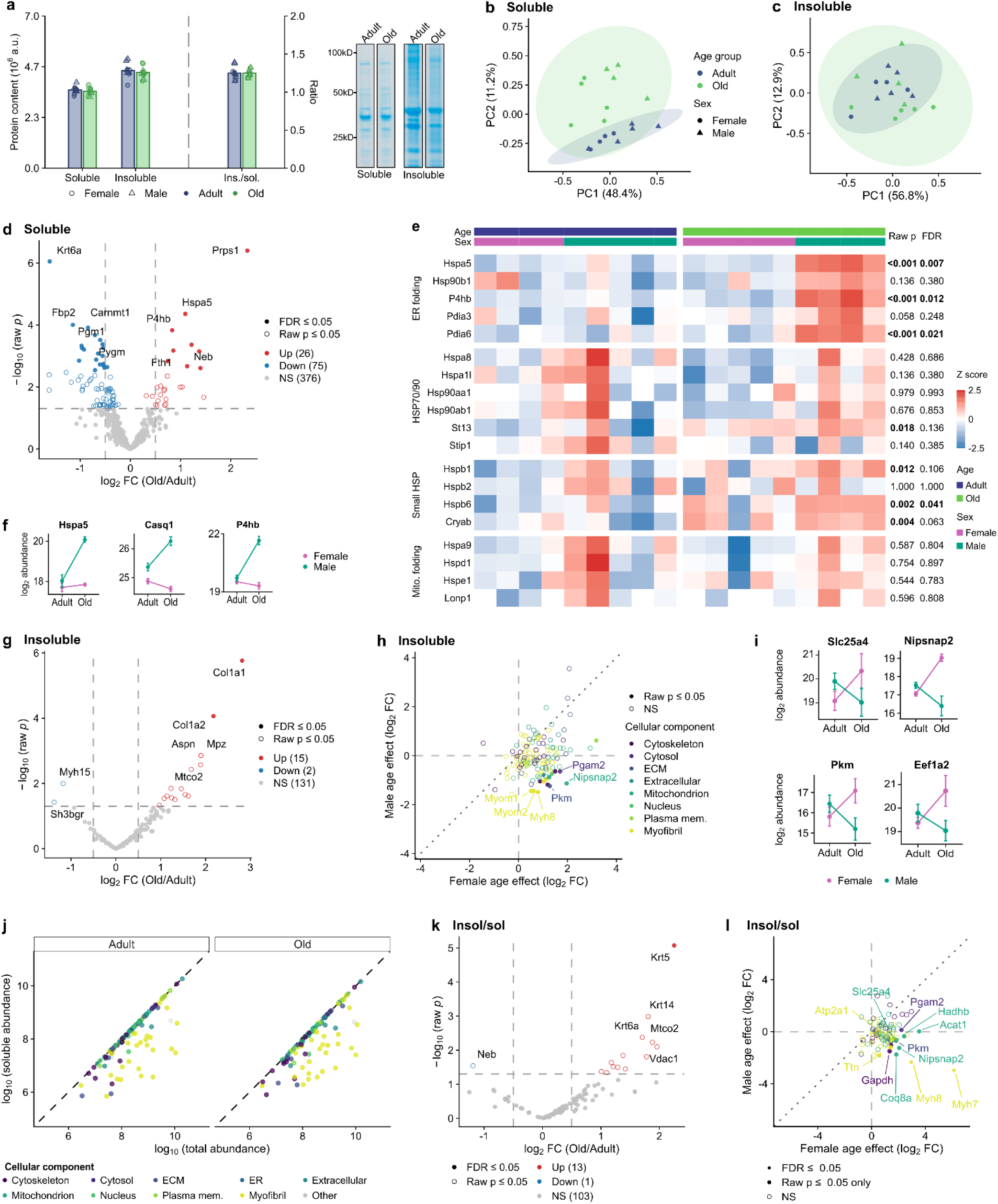
Aging induces sex-divergent proteostatic remodeling without bulk protein insolubilization. **a,** Total soluble and insoluble protein content and insoluble-to-soluble protein ratio in gastrocnemius muscle from adult and old mice, assessed by Coomassie staining. Representative gels are shown. **b,c,** Principal-component analysis (PCA) of the soluble **(b)** and insoluble **(c)** proteomes, colored by age and distinguished by sex. Ellipses represent 95% normal-theory distributions. **d,** Volcano plot of age-associated changes in soluble protein abundance. **e,** Heatmap of row-normalized protein abundances (Z-scores) for proteins involved in endoplasmic reticulum (ER) folding, HSP70/HSP90 systems, small heat-shock proteins and mitochondrial protein folding. Raw and false discovery rate (FDR)-adjusted P values are indicated. **f,** Abundance of representative proteostasis-related proteins showing sex-dependent aging responses. **g,** Volcano plot of age-associated changes in insoluble protein abundance. **h,** Female-versus male-specific aging effects on insoluble protein abundance. **i,** Abundance of representative insoluble proteins showing sex-dependent aging responses. **j,** Relationship between soluble and total protein abundance calculated from matched fractions. The dashed line indicates 100% solubility (Total abundance = soluble abundance). **k,** Volcano plot of age-associated changes in the insoluble-to-soluble protein ratio. **l,** Female-versus male-specific aging effects on protein insolubility. For all analyses, n = 4 adult female, 5 adult male, 5 old female and 4 old male mice. Values in **a**, **f** and **i** are means ± s.e.m.; symbols represent individual mice where shown. Bulk protein content (**a**) was analyzed by two-way ANOVA (age × sex, Type III sums of squares). Protein abundances and insoluble-to-soluble ratios (**b–l**) were analyzed using age × sex linear models with empirical Bayes-moderated statistics (limma). Missing protein abundances were not imputed. Benjamini–Hochberg correction was applied for multiple testing. Volcano plots show FDR-significant proteins (filled), nominally significant proteins (open) and nonsignificant proteins (grey); dashed lines indicate raw P = 0.05 and log2 fold change = ±0.5. Exact P values are shown where indicated.

We next used proteomics to test if aging altered the soluble and insoluble proteomes despite unchanged bulk protein content. Principal-component analysis showed age-associated separation of the soluble proteome with indication of sex differences (**Fig. 2b**), whereas the insoluble proteomes of adult and old mice largely overlapped (**Fig. 2c**). Differential-abundance analysis showed age-associated changes in the soluble proteome (**Fig. 2d**), with enrichment of protein-folding processes among those increased with age (**Extended data Fig. 3a,b**). Proteostasis-related proteins showed changes across ER-folding machinery, HSP70/HSP90 systems, small heat-shock proteins, and mitochondrial chaperones (**Fig. 2e**), consistent with a potential compensatory proteostatic response to the aggregated structures observed by microscopy. Several ER-associated proteins, including Hspa5, P4hb, and Casq1, increased predominantly in males (**Fig. 2f** and **Supplementary Fig. 2c**), consistent with recent evidence of sex-dependent changes in skeletal muscle proteostatic mechanisms with aging ^31^.

In contrast to the extensive remodeling of the soluble proteome with aging, comparatively few proteins changed abundance within the insoluble proteome when sexes were combined, with prominent increases in the ECM proteins Col1a1 and Col1a2 (**Fig. 2g, Extended Data Fig. 3c**). However, sex-stratified analyses revealed substantial differences obscured by the combined comparison (**Fig. 2h**). Several mitochondrial and nuclear proteins increased in insoluble abundance in females but showed little change or decreased in males (**Fig. 2i** and **Extended Data Fig. 3d**). Thus, the limited overall effect of age on the insoluble proteome partly reflects opposing responses between females and males rather than uniformly stable protein abundance across both sexes.

We next tested if the relative partitioning of individual proteins between the soluble and insoluble pools changes with aging. The overall solubility landscape was mostly unaltered by aging. Sarcomeric and myofibrillar proteins had the lowest relative solubility at both ages (**Fig. 2j**), consistent with their incorporation into stable, detergent-resistant supramolecular assemblies under physiological conditions. Only a subset of proteins showed age-associated changes in insoluble-to-soluble ratios, predominantly toward greater relative insolubility in old muscle (**Fig. 2k**). Sex-stratified analyses again showed marked differences. Several mitochondrial proteins, including Coq8, Nispsnap2, and Slc25a4, shifted toward greater insolubility with aging in females, whereas males showed smaller or opposite changes (**Fig. 2l**, **Supplementary Fig. 3**). These differences indicate sex-dependent phase-shifts with aging, rather than a uniform redistribution of the skeletal muscle proteome toward the insoluble state.

### Aging alters the skeletal muscle insoluble proteome in a compartment- and sex-dependent manner

To test whether age-associated changes in the insoluble proteome were masked by highly abundant proteins or overlooked by conventional differential abundance analysis, we examined the insoluble proteome according to cellular compartment, protein identities, and protein detection prevalence. Abundance-weighted analysis showed that the insoluble proteome was overwhelmingly dominated by sarcomeric/myofibrillar and cytoskeletal proteins across ages and sexes (**Fig. 3a**). In contrast, count-weighted analysis showed greater proteomic diversity, with proteins assigned to mitochondrial, extracellular, ECM, nuclear, cytosolic, and other cellular compartments (**Fig. 3b**), which became apparent when excluding the dominant structural components (**Fig. 3c**).

**Fig. 3.**
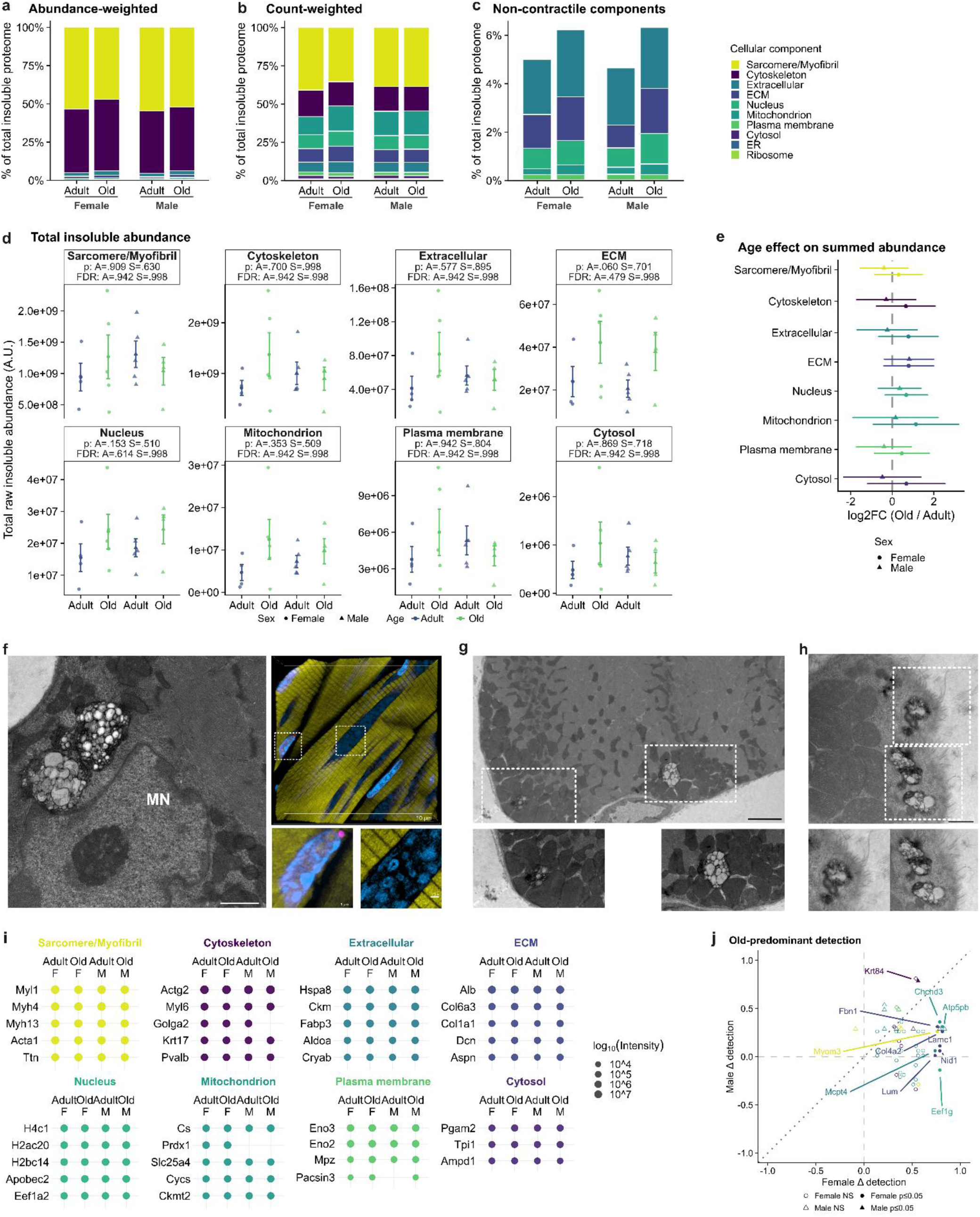
Aging alters the skeletal muscle insoluble proteome in a compartment- and sex-dependent manner. **a,b,** Relative contributions of cellular components to the insoluble proteome of adult and old female and male mice, calculated from summed protein abundances (**a**) or detected protein counts (**b**). **c,** Expanded abundance-weighted representation of less abundant cellular components, excluding sarcomeric/myofibrillar and cytoskeletal components from the display without renormalization. **d,** Summed raw insoluble protein abundances across cellular components by age (A) and sex (S). **e,** Female- and male-specific aging effects on summed protein abundance within each cellular component, expressed as log2 fold changes (Old/Adult). **f,** Qualitative transmission electron microscopy (TEM) image showing failed autophagolysosome structures adjacent to a myonucleus (MN). Fluorescence imaging of AmyTracker™ (magenta), Phalloidin/Actin (yellow), and Hoechst (cyan) in an aged (24 month) gastrocnemius muscle cell. Insets (white dotted boxes) highlight magnified views. **g,h,** Qualitative TEM images showing failed autophagolysosome structures among subsarcolemmal mitochondria and within the ECM, respectively. Dashed boxes indicate magnified regions. Scale bars are 1 micron (**f,g**) and 500 nm (**h**). **i,** Top 5 most abundant insoluble proteins within cellular components, stratified by age and sex. Dot size represents log10 protein intensity. **j,** Female-versus male-specific differences in age-associated protein detection prevalence, expressed as the difference in detection frequency between old and adult mice. For proteomic analyses (**a–e,i,j**), n = 4 adult female, 5 adult male, 5 old female and 4 old male mice. Composition analyses included 194 proteins retained by the reproducibility filter. Values in **a–c** are descriptive group means. In **d**, individual symbols represent mice, and summary values are means ± s.e.m. Summed raw abundances were analyzed using two-way ANOVA (age × sex, Type III sums of squares) on log2-transformed intensities. Points in **e** indicate estimated effects with 95% confidence intervals from the same factorial model. Relative composition was additionally analyzed using arcsine-square-root-transformed component fractions. Detection prevalence was compared using two-sided Fisher’s exact tests within each sex; filled symbols indicate raw P ≤ 0.05. Benjamini–Hochberg correction was applied across cellular components or proteins, as appropriate. Exact P and FDR values are shown where indicated.

Aging did not significantly alter summed insoluble protein abundance within any cellular component (**Fig. 3d**). Nevertheless, several lower abundance compartments, including extracellular, ECM, nuclear, and mitochondrial proteins, showed increases approaching 100% (**Fig. 3e**). When expressed relative to the total insoluble proteome, aging decreased sarcomeric/myofibrillar representation while increasing ECM, nuclear and mitochondrial components (**Extended data Fig. 4a,b**). Females had a larger reduction in sarcomeric/myofibrillar representation and greater relative increases in cytoskeletal, extracellular, and mitochondrial components, whereas males showed comparatively larger increases in ECM and nuclear representation. We then used our imaging methods to determine if compositional shifts were reflected in the spatial distribution of aggregate structures in aged skeletal muscle. We found aggregate structures in close proximity to myonuclei (**Fig. 3f**), and failed autophagolysosome structures among subsarcolemmal mitochondria (**Fig. 3g**) and within the extracellular matrix of myofibres (**Fig. 3h**). These observations provide spatial context for the nuclear, mitochondrial and ECM components identified by proteomics. Together, these results indicate that aging changes lower-abundance, non-contractile components of the insoluble proteome rather than causing broad increases in compartment-wide total insoluble abundance.

We next determined if the observed age-associated compositional changes of the insoluble proteome reflected a replacement of the most abundant proteins within each cellular compartment. Despite the age-related compositional remodeling, the identities of the most abundant insoluble proteins within each compartment were largely conserved across ages and sexes (**Fig. 3i**). Analysis of relative abundances nevertheless showed distinct age trajectories among several of these dominant proteins, including sex-specific patterns within ECM and mitochondrial compartments (**Extended Data Fig. 4c**). Thus, aging did not substantially replace the high-abundance core of individual compartments, further suggesting that much of the remodeling occurred among less-abundant proteins.

Last, we asked whether aging led to the representation of specific proteins within the insoluble pool that were absent or infrequently detected in adult muscle. We hypothesized that aging would shift normally soluble proteins in adult muscle into the insoluble pool. We found substantial sex-specificity, with 42 proteins becoming predominantly or exclusively detected in the insoluble proteome of old female muscle and only 11 in old male muscle (**Fig. 3j, Extended data Fig. 4d-f**). The female old-predominant set contained ECM proteins, including Lum, Fbn1, and Col4a2, as well as numerous mitochondrial proteins, including Atp5pb and Chchd3. In males, the old-predominant set was smaller, with Krt84 having a significant within-sex increase in detection. Thus, detection-prevalence analysis showed that a greater variety of proteins went “over the edge” of solubility in aged female mice compared to aged male mice.

Collectively, these findings show that aging primarily alters the representation of lower-abundance mitochondrial, ECM and nuclear-associated proteins in the insoluble proteome, while largely preserving its dominant proteins, with more pronounced changes in females than males.

### Intrinsic aggregation propensity shows sex- and compartment-dependent associations with protein insolubility

We hypothesized that age-related changes in protein insolubility were related to intrinsic sequence-derived aggregation propensity. We evaluated three complementary metrics: AGGRESCAN (general aggregation propensity)^32^, TANGO (β-aggregation propensity)^33^ and WALTZ (amyloid-forming propensity)^34^. Using proteins detected in the adult soluble proteome as our starting population, we compared predicted propensity across cellular components and identified the highest-scoring proteins. AGGRESCAN and WALTZ differed significantly among components (**Fig. 4a**). In particular, the mitochondrial compartment had greater general aggregation propensity than several other compartments, including the nuclear, extracellular, cytoskeletal, and sarcomeric/myofibrillar (**Fig. 4a**). Consistent with this compartment-level pattern, mitochondrial proteins were frequently represented among the highest-scoring proteins for aggregation (**Fig. 4b**).

**Fig. 4.**
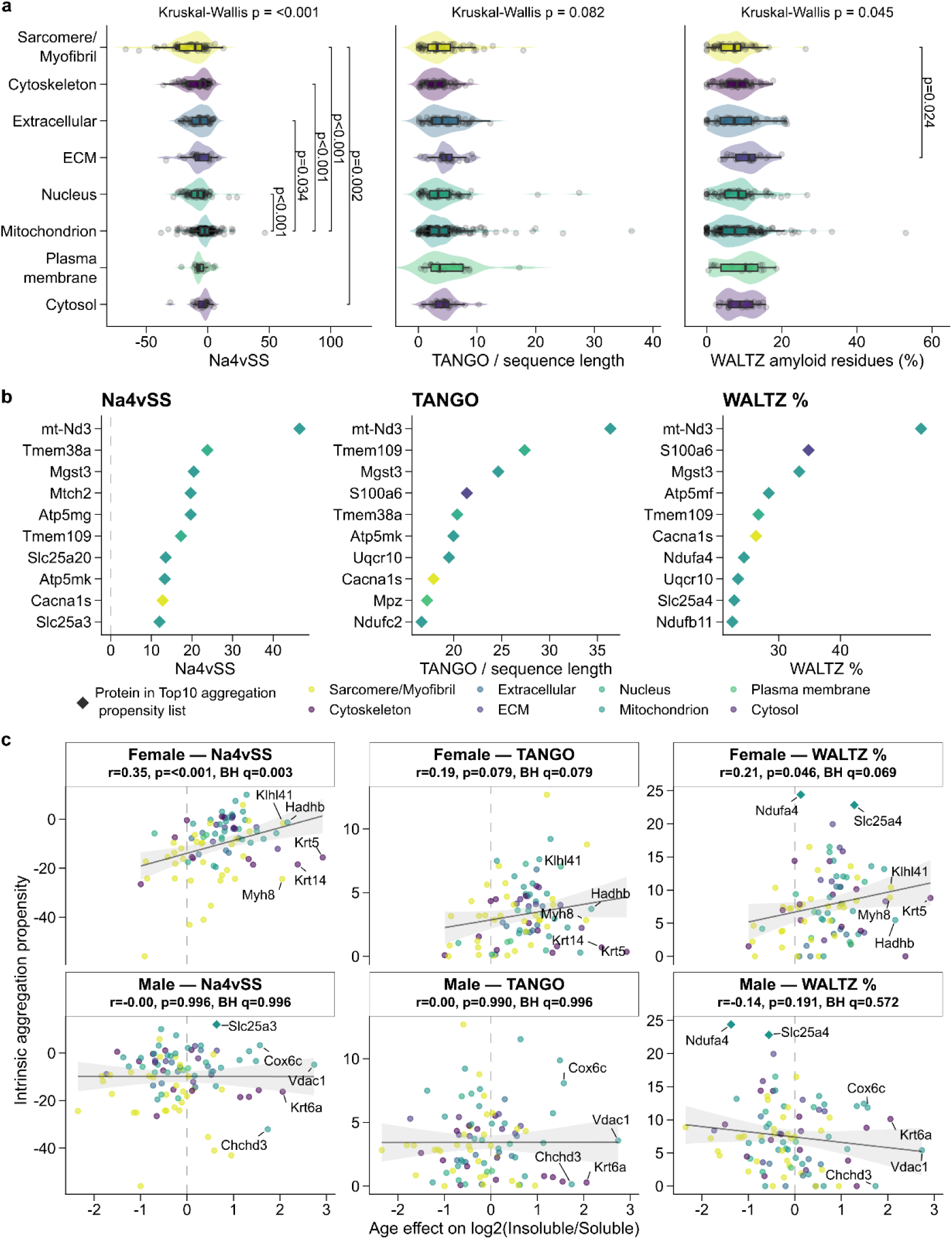
Intrinsic aggregation propensity shows sex- and compartment-dependent associations with protein insolubility. **a,** Distributions of intrinsic aggregation-propensity scores across cellular components, including AGGRESCAN Na4vSS, TANGO score normalized to protein sequence length and WALTZ-predicted amyloid-forming residues (%). Each point represents an individual protein. Violins indicate score distributions and boxplots indicate medians and interquartile ranges. **b,** Ten proteins with the highest predicted aggregation propensity according to each metric, ranked from proteins detected in the adult soluble fraction and colored by cellular-component annotation. **c,** Relationships between age-associated changes in the log2 insoluble-to-soluble protein ratio and intrinsic aggregation propensity, analyzed separately in females and males. Each point represents an individual protein, colored by cellular-component annotation, with proteins detected in the top 10 shown as diamonds and selected proteins labeled. Solid lines indicate linear regressions, with shaded 95% confidence intervals. Aggregation-propensity metrics were calculated from amino acid sequences and represent protein-level properties rather than independent biological replicates. Proteomic measurements were obtained from n = 4 adult female, 5 adult male, 5 old female and 4 old male mice. Differences in propensity scores among cellular components (**a**) were assessed using Kruskal–Wallis tests followed by Dunn’s multiple-comparison tests with Holm adjustment. Associations in **c** were evaluated using two-sided Pearson correlation tests, with Benjamini–Hochberg correction across the three aggregation metrics within each sex. Correlation coefficients (r), raw P values and adjusted q values are shown in each panel. The vertical dashed line indicates no age-associated change in protein insolubility (log2 fold change = 0). The number of proteins in each correlation depends on the availability of valid paired soluble and insoluble abundances.

We next tested whether proteins with greater intrinsic propensity showed larger age-associated shifts toward insolubility. In females, general and amyloid aggregation propensities were positively associated with the age-associated increase in the insoluble-to-soluble ratio, whereas β-aggregation propensity had no significant association (**Fig. 4c**). In contrast, none of these relationships were significant in males (**Fig. 4c**). However, high intrinsic propensity did not consistently predict individual protein fate, as only a subset of the highest-scoring proteins showed substantial age-associated shifts toward insolubility. Thus, although mitochondrial proteins have a relatively aggregation-prone sequence and are increasingly represented in the aged insoluble proteome, intrinsic propensity, at least as defined by current bioinformative resources, did not predict the fate of individual mitochondrial proteins.

We next assessed the relationship between aggregation propensity and age-related solubility shifts within each compartment (**Extended Data Fig. 5b-d**). These relationships were generally non-significant and inconsistent across compartments, except among female sarcomeric/myofibrillar proteins, where greater propensity correlated positively with age-associated shifts toward insolubility across all three metrics. Consistent with this context-dependent relationship, the aged insoluble proteome was not globally enriched for proteins with greater predicted intrinsic aggregation propensity. Mean AGGRESCAN, TANGO, and WALTZ scores did not differ significantly with age, sex, or their interaction (**Extended Data Fig. 5a**). Thus, aging does not appear to simply increase the representation of the most intrinsically aggregation-prone proteins in the insoluble proteome.

We further asked whether intrinsic aggregation propensity could explain why specific proteins become predominantly or exclusively detected in the aged insoluble fraction (**Fig. 3j**, **Extended Data Fig. 4e,f**). The old-predominant insoluble proteins had neither significantly greater aggregation propensity nor enrichment among the highest-scoring proteins relative to proteins detected in both age groups in either sex (**Extended Data Fig. 5e-g**). Nevertheless, several proteins, including Atp5pb and Lum, showed high predicted aggregation propensity (**Supplementary Fig. 4a-c**).

Collectively, our results show that higher intrinsic aggregation propensity is associated with greater age-related shifts toward insolubility in females, particularly among sarcomeric/myofibrillar proteins, whereas these associations are largely absent in males and other cellular components. However, neither the aged insoluble proteome overall nor proteins preferentially detected in old muscle were enriched for high-propensity sequences, indicating that intrinsic aggregation propensity alone does not explain which proteins become more prevalent in the insoluble fraction with aging.

### Exercise and rapamycin alter distinct features of the aged insoluble proteome

We next asked whether exercise or mTOR inhibition would modify the aged insoluble proteome, indicating a change in proteostatic maintenance. Old mice underwent voluntary wheel running or pharmacological mTOR inhibition with rapamycin for up to 60 days. Neither intervention altered the bulk total soluble or insoluble protein content or their ratio (**Fig. 5a**), and principal-component analyses showed substantial overlap among groups in both fractions (**Fig. 5b,c**). Stable-isotope labeling further showed that exercise did not significantly alter synthesis or turnover of the myofibrillar, mitochondrial, or cytosolic fractions, nor the fraction of newly synthesized proteins within the soluble or insoluble proteome (**Extended Data Fig. 6a-d**). In contrast, rapamycin reduced myofibrillar protein synthesis and soluble fraction new after 60 days of labeling (**Extended Data Fig. 6a,d**). Thus, intervention-associated proteomic alterations occurred without changes in bulk protein content and minor changes in protein synthesis.

**Fig. 5.**
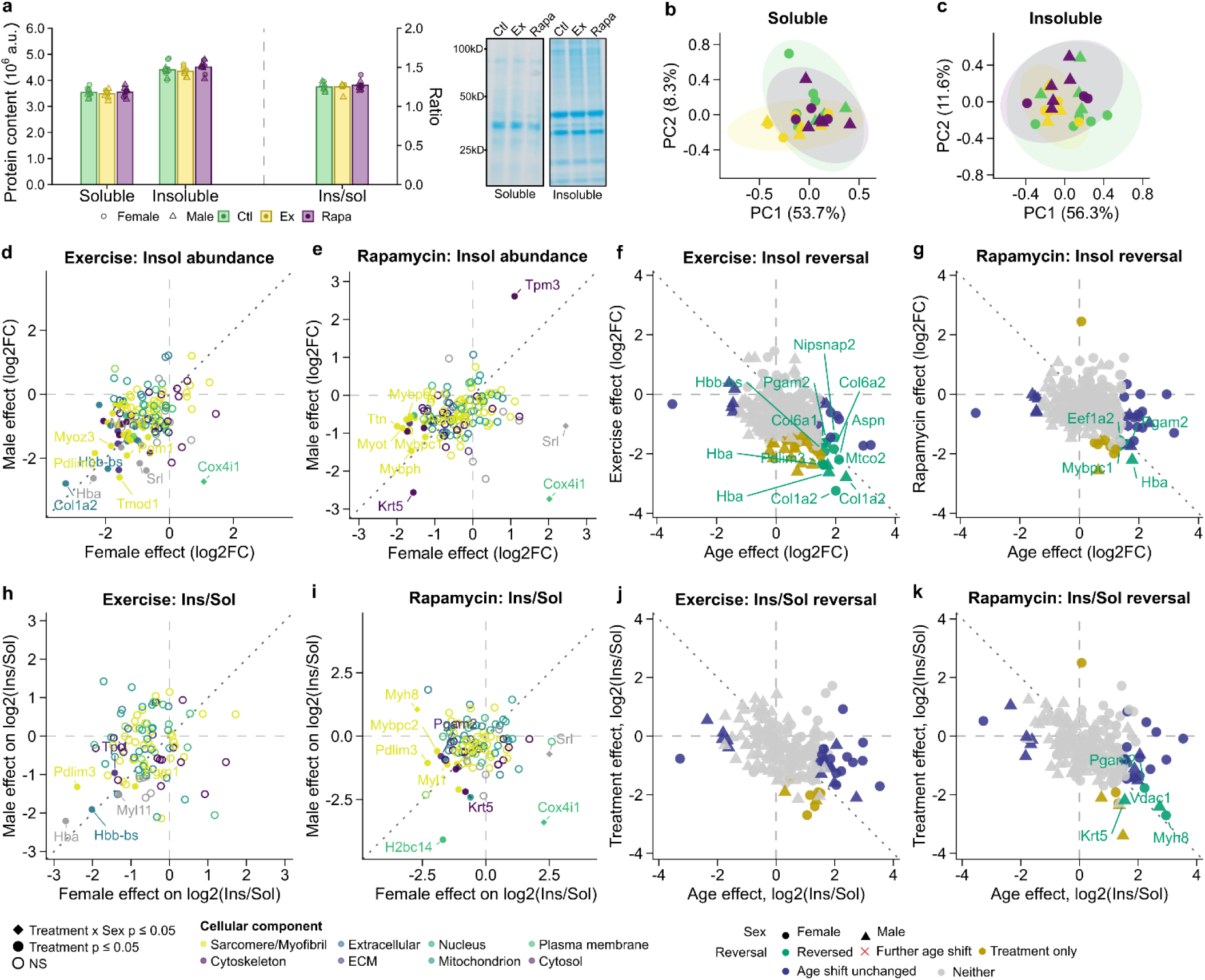
Exercise and rapamycin alter distinct features of the aged insoluble proteome. **a,** Total soluble and insoluble protein content and insoluble-to-soluble protein ratio in old control (Ctl), exercise (Ex) and rapamycin-treated (Rapa) mice, assessed by Coomassie staining, with representative gels. **b,c,** Principal-component analysis (PCA) of soluble (**b**) and insoluble (**c**) proteomes, distinguished by treatment and sex. **d,e,** Female-versus male-specific effects of exercise (**d**) and rapamycin (**e**) on insoluble protein abundance, expressed as log2 fold changes relative to old controls. **f,g,** Relationships between age-associated changes in insoluble protein abundance and subsequent effects of exercise (**f**) or rapamycin (**g**), showing proteins classified by directional reversal of aging effects. **h,i,** Female-versus male-specific effects of exercise (**h**) and rapamycin (**i**) on the log2 insoluble-to-soluble protein ratio. **j,k,** Relationships between age-associated changes in protein insolubility and subsequent effects of exercise (**j**) or rapamycin (**k**), using the same reversal classifications. For all analyses, n = 5 control female, 4 control male, 4 exercise female, 4 exercise male, 3 rapamycin female and 5 rapamycin male mice. Values in **a** are means ± s.e.m. and symbols represent individual mice. Bulk protein content was analyzed by two-way ANOVA (treatment × sex, Type III sums of squares). Protein-level changes were analyzed using treatment × sex linear models with empirical Bayes-moderated statistics (limma), including sex-specific treatment contrasts and treatment-by-sex interactions. Benjamini–Hochberg correction was applied for multiple testing. In **d,e,h,i**, colors indicate cellular-component annotations, filled points indicate nominally significant treatment effects (P ≤ 0.05), and diamonds indicate nominally significant treatment-by-sex interactions. In **f,g,j,k,** colors indicate reversal classifications, including reversal, further age-associated shift, unchanged age-associated shift, treatment-only change or neither; circles and triangles represent females and males, respectively. Dotted diagonals indicate equal treatment effects between sexes or complete directional reversal of the aging effect, as appropriate.

Despite the limited changes observed at the bulk level, both interventions altered solubility of individual proteins. In the soluble proteome, exercise primarily affected metabolic and proteostasis-related proteins, including decreased abundance of several HSP70/HSP90 proteins (**Extended Data Fig. 7a,b)**, while rapamycin produced fewer changes involving mitochondrial and structural proteins (**Extended Data Fig. 7e**). In the insoluble proteome, exercise produced broader abundance changes than rapamycin (**Fig. 5d,e**), predominantly reducing structural, glycolytic and ECM proteins, including the age-elevated Col1a2, as well as SR-associated proteins such as sarcalumenin and Casq1, previously identified as components of tubular aggregates ^29,30^ (**Fig. 5d; Extended Data Fig. 7c**). Rapamycin affected fewer insoluble proteins, primarily reducing selected contractile and structural proteins (**Fig. 5e; Extended Data Fig. 7f**). Although response magnitudes varied, most treatment-responsive proteins changed in the same direction in females and males, with relatively few showing significant treatment-by-sex interactions.

To determine whether these treatment-associated changes in the insoluble proteome specifically corrected those induced by aging, we directly compared the age effect (Old Control − Adult Control) with the treatment effect (Exercise or Rapamycin − Old Control) in females and males (**Fig. 5f,g**). Exercise reversed more age-associated increases in insoluble protein abundance in females than males (**Fig. 5f**), while rapamycin reversed changes in fewer proteins, consistent with its more limited effect on the insoluble proteome (**Fig. 5g**). Many treatment-responsive proteins did not correspond to proteins significantly increased with aging, indicating that both interventions produced effects beyond reversal of the aged state.

Consistent with the aging comparison (**Fig. 2j**), the protein solubility landscape of skeletal muscle remained largely unchanged across treatments (**Extended data Fig. 7h**). Nevertheless, both interventions altered the partitioning of selected proteins between the soluble and insoluble pools. Notably, despite reducing the insoluble abundance of numerous proteins, exercise decreased insoluble-to-soluble ratios for relatively few proteins (**Fig. 5h**, **Extended Data Fig. 7d**) and showed limited reversal of age-associated partitioning changes (**Fig. 5j**). Thus, most exercise-associated reductions in insoluble abundance were not accompanied by preferential redistribution toward the soluble pool. In contrast, rapamycin altered the insoluble abundance of fewer proteins but produced a broader reduction in insoluble-to-soluble ratios (**Fig. 5i, Extended Data Fig. 7g**), reversing age-associated partitioning changes in a subset of proteins (**Fig. 5k**).

Collectively, our findings demonstrate that exercise and rapamycin differentially alter insoluble protein abundance and partitioning without changing bulk insoluble protein content, suggesting distinct mechanisms through which these interventions may influence proteostasis in aged muscle.

### Exercise and rapamycin alter lower-abundance components of the aged insoluble proteome

The insoluble proteome of aged skeletal muscle remained dominated by highly abundant structural proteins following exercise and rapamycin (**Fig. 6a–c**). However, both treatments reduced the summed insoluble abundance of several non-contractile compartments, including extracellular, ECM, nuclear, mitochondrial, and plasma-membrane proteins, with generally larger effects for exercise (**Fig. 6d,e**). Both treatments increased the relative representation of sarcomeric/myofibrillar proteins and reduced that of cytoskeletal proteins, partially opposing age-associated compositional changes (**Extended Data Fig. 8a,b**). Importantly, the increased sarcomeric/myofibrillar representation reflected reductions in other components rather than greater absolute sarcomeric abundance. These compartment-level changes observed with exercise and rapamycin were generally similar between females and males, again indicating less sex-specificity compared to the effects of aging. Despite these compositional shifts, neither intervention substantially changed the identity of the most abundant proteins within each cellular component (**Fig. 6f, Extended Data Fig. 8c**), indicating that the high-abundance core of the insoluble proteome remained largely unaltered.

**Fig. 6.**
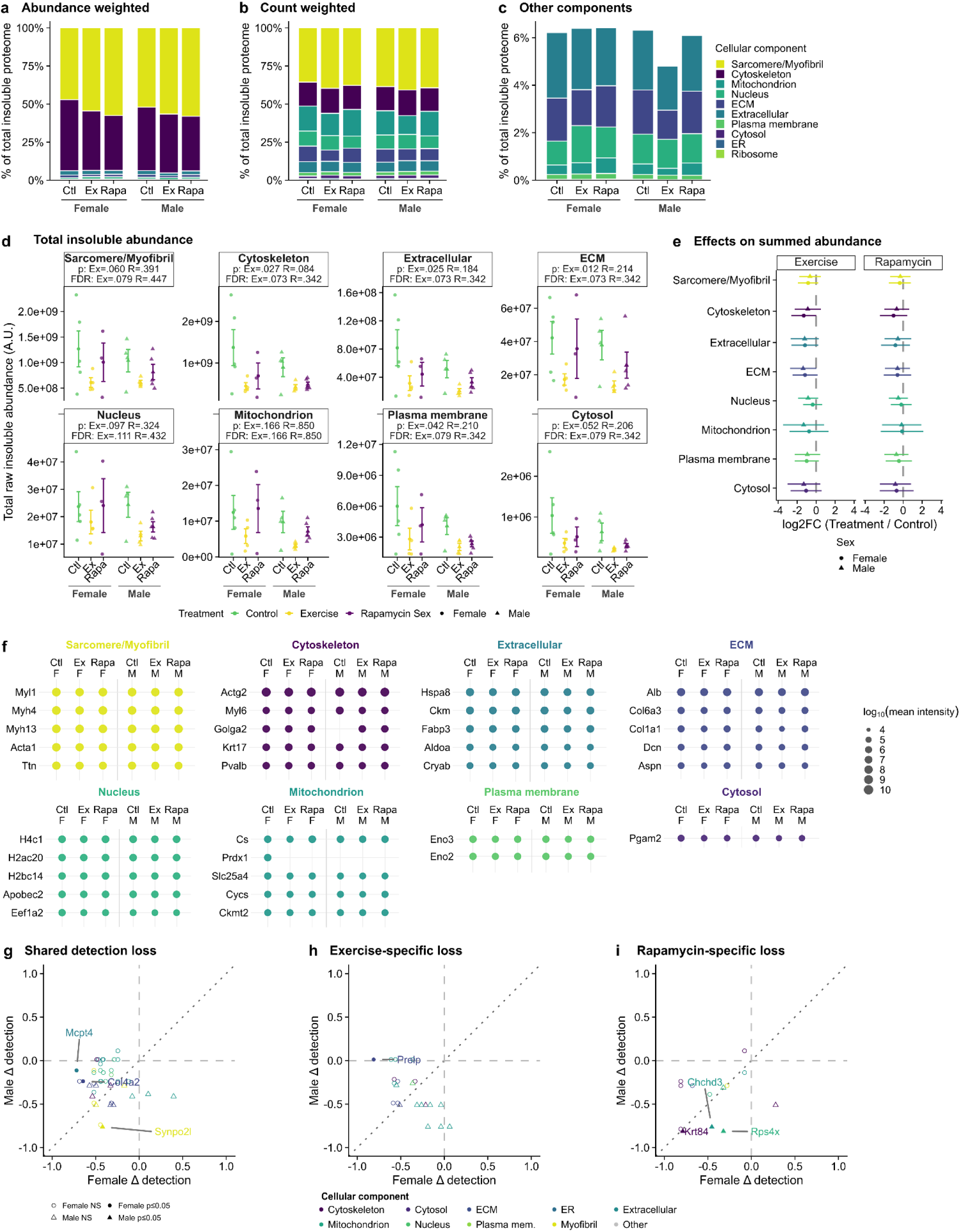
Exercise and rapamycin alter lower-abundance components of the aged insoluble proteome. **a,b,** Relative contributions of cellular components to the insoluble proteome of old control (Ctl), exercise (Ex) and rapamycin-treated (Rapa) mice, calculated from summed protein abundances (a) or detected protein counts (b), separately for females and males. **c,** Expanded abundance-weighted representation of less abundant cellular components. **d,** Summed raw insoluble protein abundances across cellular components by treatment and sex. **e,** Exercise- and rapamycin-associated changes in summed protein abundance within each cellular component, expressed as log2 fold changes relative to old controls. **f,** Top 5 most abundant insoluble across treatment and sex groups, organized by cellular-component annotation. Dot size represents log10 protein intensity. **g–i,** Female-versus male-specific changes in protein detection prevalence for proteins showing reduced detection after both interventions (**g**), exercise-specific loss of detection (**h**) or rapamycin-specific loss of detection (**i**). Selected proteins are labeled. For all analyses, n = 5 control female, 4 control male, 4 exercise female, 4 exercise male, 3 rapamycin female and 5 rapamycin male mice. Composition analyses included 197 proteins retained by the reproducibility filter. Values in **a–c** are descriptive group means. In **d**, symbols represent individual mice and summary values are means ± s.e.m. Summed raw abundances were analyzed using treatment × sex linear models, with prespecified exercise-versus-control and rapamycin-versus-control contrasts on log2-transformed intensities. Relative composition was analyzed using arcsine-square-root-transformed component fractions and Type III ANOVA. Points in **e** indicate estimated sex-averaged treatment effects with 95% confidence intervals. Detection prevalence was assessed using two-sided Fisher’s exact tests within each sex and treatment comparison. Filled symbols in **g–i** indicate raw P ≤ 0.05; open symbols indicate nonsignificant differences. Benjamini–Hochberg correction was applied across tested components or proteins, as appropriate. Exact P and adjusted values are shown where indicated.

Because aging caused several lower-abundance proteins to become represented in the insoluble proteome, we next asked whether exercise or rapamycin reversed this phenotype by reducing their detection in aged muscle. We identified numerous proteins that were removed or became less consistently detected in the insoluble proteome after both interventions, particularly in females (**Fig. 6g, Extended data Fig. 8d,e**). Notably, Col4a2, whose detection increased in aged females, showed reduced detection after both treatments, although the rapamycin effect did not reach statistical significance. Additional treatment-specific losses involved ECM and mitochondrial proteins after exercise (**Fig. 6h)** and selected cytoskeletal, nuclear and mitochondrial proteins after rapamycin (**Fig. 6i, Extended Data Fig. 8d,e**).

Together, these findings suggest that exercise and rapamycin may prevent the entry to or promote the clearance of lower-abundance proteins from the insoluble proteome, including proteins that become more prevalent with aging, while largely preserving its high-abundance structural components.

### Intrinsic aggregation propensity does not broadly predict intervention-associated changes in the aged insoluble proteome

Finally, we tested if intrinsic aggregation propensity correlated with exercise- and rapamycin-induced shifts in protein solubility, using proteins detected in the old-control insoluble fraction as our starting population. Consistent with the aging analyses, mitochondrial proteins were frequently represented among those with the highest predicted aggregation propensity (**Fig. 7a**). In females, greater general aggregation propensity (AGGRESCAN) was associated with exercise-induced decreases in insoluble-to-soluble protein ratios (**Fig. 7b**), opposite to the positive association observed with aging. However, these associations did not consistently involve the highest-scoring proteins. Neither β-aggregation (TANGO) nor amyloid-forming propensity (WALTZ) was significantly associated with exercise responses, and no significant relationships were detected in males (**Fig. 7b**). Rapamycin-induced changes in insoluble-to-soluble partitioning were not significantly correlated with any aggregation propensity metric in females or males (**Fig. 7c**).

**Fig. 7.**
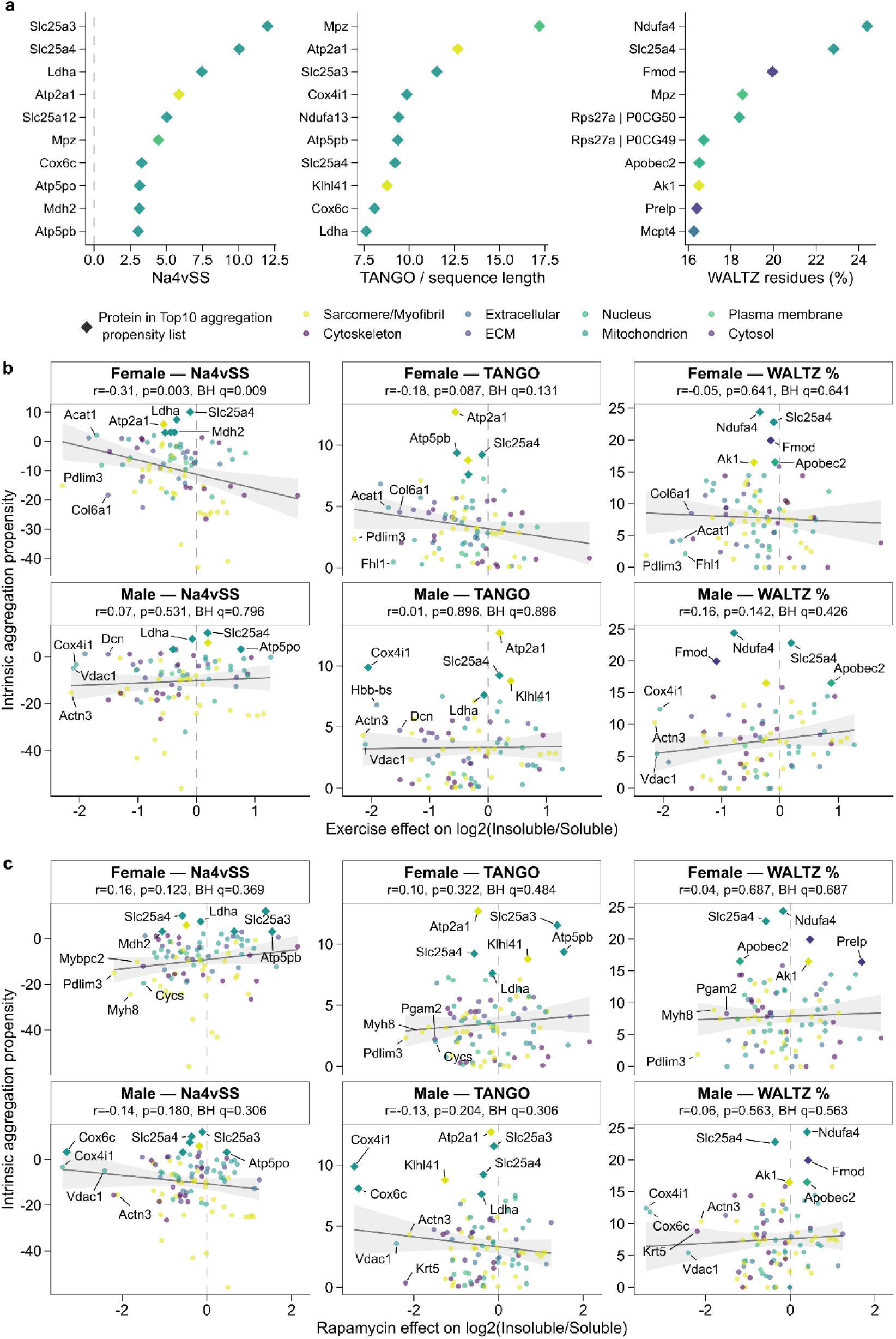
Intrinsic aggregation propensity does not broadly predict intervention-associated changes in the aged insoluble proteome. **a,** Ten proteins with the highest predicted intrinsic aggregation propensity according to AGGRESCAN Na4vSS, TANGO score normalized to sequence length and WALTZ-predicted amyloid-forming residues (%), ranked from proteins detected in the old-control insoluble fraction and colored by cellular-component annotation. **b,c,** Relationships between intrinsic aggregation propensity and changes in the log2 insoluble-to-soluble protein ratio following exercise (**b**) or rapamycin treatment (**c**), analyzed separately in females and males. Each point represents an individual protein, colored by cellular-component annotation, with selected proteins labeled. Solid lines indicate linear regressions, with shaded 95% confidence intervals. The reference protein pool was defined by detection in at least two old-control insoluble samples, irrespective of sex. Sex-specific treatment analyses required at least two valid measurements per group in both soluble and insoluble fractions. The proteomic cohort comprised n = 5 control female, 4 control male, 4 exercise female, 4 exercise male, 3 rapamycin female and 5 rapamycin male mice. Aggregation-propensity scores were calculated from protein amino acid sequences. Associations were evaluated using two-sided Pearson correlation tests, with Benjamini–Hochberg correction across the three aggregation metrics within each treatment and sex. Correlation coefficients (r), raw P values and adjusted q values are shown in each panel. Dashed vertical lines indicate no treatment-associated change in protein insolubility (log2 fold change = 0). Numbers of proteins included in each correlation depend on valid paired soluble and insoluble measurements.

Compartment-specific analyses further indicated that the relationship between intrinsic aggregation propensity and intervention responsiveness was heterogeneous. Exercise-induced decreases in insoluble-to-soluble ratio correlated with greater AGGRESCAN propensity in female sarcomeric/myofibrillar and cytoskeletal proteins, whereas rapamycin-associated partitioning changes correlated positively with TANGO in male nuclear proteins and negatively with WALTZ in female cytoskeletal proteins (**Extended Data Fig. 9a-f**). Despite these relationships, neither intervention altered the overall distributions of general, β-aggregation, or amyloid-forming propensity within the insoluble proteome (**Extended Data Fig. 10a**), indicating that treatment did not globally shift the insoluble proteome toward proteins with lower intrinsic aggregation propensity.

We further tested whether proteins with reduced insoluble detection after treatment had greater intrinsic aggregation propensity. No consistent differences were observed, except for higher general aggregation propensity among proteins showing exercise-specific loss of detection in males, predominantly mitochondrial proteins (**Extended Data Fig. 10b**). At the individual protein level, several mitochondrial proteins with intervention-induced loss of detection, including Slc25a3 and Cox6c, showed high predicted aggregation propensity despite the absence of a consistent group-level association (**Supplementary Fig. 9a-g**).

Collectively, our findings indicate that intrinsic aggregation propensity is associated with selected sex- and compartment-specific treatment responses, but is not a global determinant of intervention-induced changes in the aged insoluble proteome.

## DISCUSSION

In this study, we show that aged skeletal muscle accumulates distinct aggregate structures and undergoes changes in the abundance and composition of its insoluble proteome, despite unchanged bulk insoluble content. These shifts primarily involved lower-abundance proteins and varied across cellular compartments. We observed striking sex differences with females showing more pronounced age-associated changes in the insoluble proteome than males. Exercise and rapamycin partially reversed age-associated changes in the insoluble proteome, demonstrating that these changes are modifiable and highlighting the potential to develop new therapeutic approaches. Together, our findings provide an alternative approach to anabolic-centric treatments for maintaining muscle quality with age (**Fig. 8**).

**Fig. 8.**
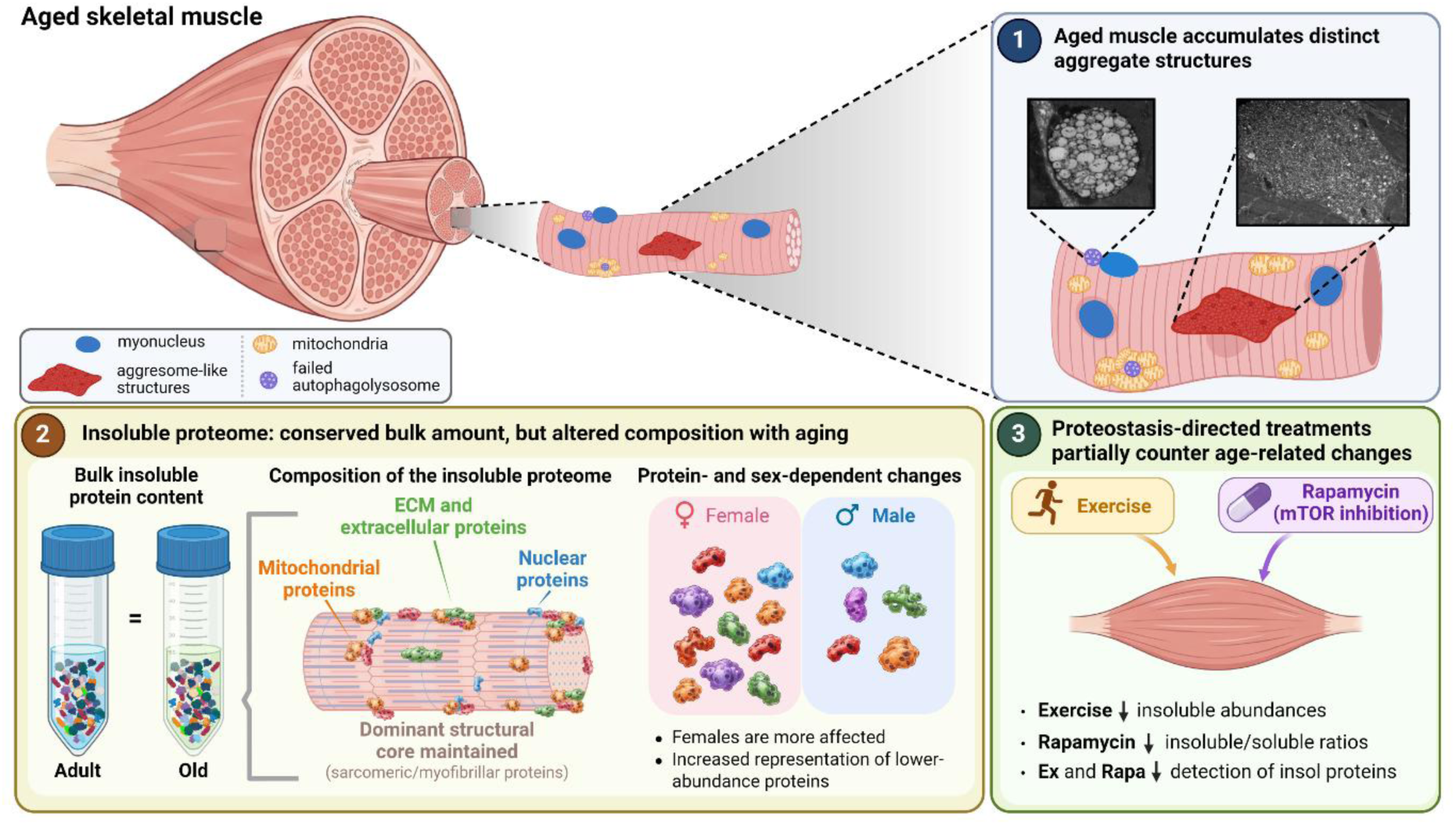
Summary of age-associated changes in skeletal muscle protein aggregation and insolubility and their modulation by exercise and rapamycin. **(1)** Aging is associated with the accumulation of distinct aggregate-like structures, including aggresome-like and failed autophagolysosome structures. Representative transmission electron micrograph illustrates aggresome-like and failed autophagolysosomes in aged skeletal muscle. **(2)** Despite unchanged bulk insoluble protein content, aging alters the abundance and composition of the insoluble proteome, particularly among lower-abundance mitochondrial, nuclear, and extracellular matrix proteins, while the dominant sarcomeric/myofibrillar component remains largely preserved. These changes are more pronounced in females than males. **(3)** Exercise and rapamycin partially counteract age-associated changes in the insoluble proteome through distinct responses: exercise produces broader reductions in individual insoluble protein abundances, whereas rapamycin produces more widespread reductions in insoluble-to-soluble protein ratios. Both interventions also reduce the detection of subsets of age-associated insoluble proteins. Created in BioRender. Mesquita, P. (2026) https://BioRender.com/xtzfjs4

Protein aggregation and loss of protein solubility are related but distinct manifestations of proteostatic decline. Beyond the well-described SR-derived tubular aggregates, our imaging showed an age-related accumulation of distinct aggresome-like and amyloid-positive structures, often beneath the sarcolemma. Some of the large membrane-rich structures observed in our study resemble the autophagosome–lysosomal structures ^35^ and multi-vesicular bodies ^36^ previously reported in aged muscle. In contrast, protein insolubility reflects both biological factors and the biochemical conditions used for extraction and does not, by itself, indicate pathological aggregation. Structural and contractile proteins accounted for approximately 95% of the insoluble proteome despite their relatively low predicted aggregation propensity, consistent with their incorporation into stable assemblies such as myofilaments, Z-discs and intermediate-filament networks under physiological conditions. This dominance of structural and contractile proteins indicates that bulk insoluble protein abundance is unlikely to adequately capture proteostatic decline and can actually obscure the age-associated solubility shifts of lower-abundance proteins or cellular compartments.

Proteome solubility is increasingly recognized as context dependent. Stephan et al. ^24^ recently demonstrated substantial tissue-specific variation in age-associated protein insolubility. Based on our previous work ^5^, we expected contractile proteins to account for much of the age-related changes in the insoluble proteome. Instead, its structural core remained comparatively stable, whereas lower-abundance mitochondrial, nuclear and ECM proteins showed substantial age-related changes in insoluble abundance and partitioning. These shifts in lower-abundance cellular compartments suggest that protein insolubility and proteostatic vulnerability in aged muscle are determined not only by intrinsic protein properties but also by the cellular context in which proteins are assembled and maintained. Our super-resolution microscopy further showed aggregate structures localized to distinct subcellular regions, consistent with evidence for accumulation of aggregates through compartment-specific quality-control pathways ^37^.

The age-related increase in insolubility of mitochondrial and ECM proteins is particularly interesting given their fundamentally distinct cellular environments and protein quality control systems. Skeletal muscle mitochondria form a highly dynamic network with coordinated protein import, folding, respiratory-complex assembly and turnover, while ECM proteins often form long-lived, highly crosslinked assemblies. Impaired mitochondrial protein import, for example, can promote cytosolic aggregation of mitochondrial proteins, which in turn can promote secondary protein aggregation ^38^. Consistent with our findings, Stephan et al. ^24^ reported greater insolubility of mitochondrial and ECM proteins across tissues, including skeletal muscle. The fact that proteins from such biologically different compartments both increase insolubility with age argues against a single mechanism driving the aged insoluble proteome and instead supports multiple, compartment-dependent routes to proteostatic failure. Consistent with this interpretation, intrinsic aggregation propensity did not globally predict age-related changes in the insoluble proteome, but showed sex- and compartment-dependent associations. Thus, sequence-derived aggregation propensity alone is unlikely to explain the complex changes in protein insolubility observed with aging.

One of our most striking findings was the greater extent of age-associated changes in the insoluble proteome of females than males, spanning protein abundance, partitioning, detection and associations with aggregation propensity. This finding contrasts with the male-biased accumulation of SR-derived tubular aggregates reported in aged mice ^22,29,30,39^, suggesting these phenomena reflect separate aspects of muscle aging. This distinction may be clinically relevant, as older women experience greater mobility limitations than men ^40,41^, although the functional relevance of our proteomic findings remains unknown. Importantly, our microscopy experiments were not designed or sufficiently powered to determine whether sex differences in proteome solubility are accompanied by corresponding differences in aggregate type, burden, or morphology.

Our intervention studies demonstrate that the age-related changes in the insoluble proteome is a modifiable phenotype of aged muscle. Neither voluntary exercise nor chronic mTOR inhibition changed bulk insoluble proteome content, yet both affected the abundance and partitioning of individual proteins in distinct ways. Exercise produced broader reductions in insoluble abundance across structural, metabolic, ECM and SR-related proteins, whereas rapamycin affected fewer insoluble proteins but produced more widespread decreases in insoluble-to-soluble protein ratios. Both interventions also reduced the representation or detection of several lower-abundance mitochondrial, ECM and other non-contractile proteins that were among the components most affected by aging, while largely preserving the dominant structural core. These intervention-driven responses were similar between females and males, suggesting that the strong sex dependence in the age-related proteostatic disruption does not necessarily translate into sex-dependent responsivesses to interventions. Notably, exercise altered the aged insoluble proteome without increasing bulk protein synthesis or turnover, while rapamycin corrected select age-related changes despite reducing myofibrillar protein synthesis. Thus, increasing protein synthesis was not required for the observed changes in the aged proteome. These findings do not support the prevailing anabolic framework of muscle aging, in which preservation of muscle is often approached primarily through increasing protein synthesis. Our data instead support a model in which improving proteostatic mechanisms can change the representation of proteins in soluble and insoluble fractions. In this context, increasing synthesis in an aged cell with constrained folding and degradation capacity could theoretically increase proteostatic load and worsen the burden on quality control systems. Whether the reduced representation of insoluble proteins reflects decreased recruitment or enhanced clearance remains unresolved. Nevertheless, our findings support further investigation of proteostatic mechanisms as potential therapeutic targets in aged muscle.

In summary, we demonstrate that aging is associated with the accumulation of distinct aggregate-like structures and sex- and compartment-dependent changes in the skeletal muscle insoluble proteome that are not captured by bulk protein measurements. Exercise and rapamycin partially counteract these alterations, demonstrating that protein insolubility can be modified without increasing bulk protein synthesis. Together, these findings highlight protein quality and fate, rather than protein quantity alone, as important targets for preserving skeletal muscle health with aging.

## METHODS

### Animals and experimental design

Animal experiments were approved by the OMRF Institutional Animal Care and Use Committee. Adult (4–6-month-old) and old (22–24-month-old) male and female C57BL/6 and C57BL/6J mice were housed with ad libitum food and water under a 14-h light/10-h dark cycle. Animals were group-housed until random assignment to adult control, old control, old exercise or old rapamycin groups. Old mice underwent voluntary wheel running or received encapsulated rapamycin chow (42 ppm; Emtora Biosciences and Purina Lab Diet) for up to 60 days, concurrent with D₂O labeling. Mice were euthanized via CO₂ followed by cervical dislocation and cardiac exsanguination. Gastrocnemius muscle was collected, flash-frozen and stored at −80 °C.

### Immunohistochemistry (IHC)

Sample preparation for super-resolution light microscopy including Proteostat™ (Enzo Life Sciences), AmyTracker 680™ (Ebba Biotech AB) as well as anti-Serca1 antibody (Thermo) labeling were performed as follows. Briefly, mice were euthanized and the tibialis anterior muscle was excised and immediately submerged in fixative solution (i.e., 3% glyoxal, 1% acetic acid in pure water, pH 4.5; Electron Microscopy Sciences) for overnight incubation at 4°C. The following day, samples were washed with 1X PBS and serially submerged in increasing concentrations of sucrose diluted in 1X PBS for 6 hr increments at 4°C (i.e., 10, 30, and 45% sucrose; Sigma). Cryoprotected samples were then cryopreserved using liquid nitrogen chilled isopentane in stainless steel cryomolds (Pelco). Ten-micron cryostat-generated tissue sections were retrieved from −80°C storage, thawed, and prepared for IHC by hydrophobic barrier using a pap-pen (VWR). Samples were then hydrated with 1X PBS, blocked (i.e., 5% IgG-free bovine serum albumin in 1X PBS; Jackson), and incubated with primary antibody (anti-Serca1 antibody in blocking buffer) overnight at 4°C. The following day, samples were washed and diluted in species isotype matched secondary VH-domain secondary antibodies (i.e., 2 μg/ml) for 1 hr at room temperature (RT). For all dyes (i.e., cytoskeletal actin, nuclei, Proteostat™, and AmyTracker 680™), sections were incubated for one hour incubation at RT followed by 1X PBS washing and cover slipping using Prolong Glass (Invitrogen) and 1.5H coverslip mounting (Corning).

### Subcellular light microscopy and quantitative analysis

Tissue sections were imaged using a Zeiss LSM 980 multiphoton microscope with airyscan 2.0 deconvolution and a 63X 1.4 NA objective (Zeiss). A minimum of 20 volumetrically sampled muscle cells were subcellularly sampled across three biological replicates per age group where statistical considerations were performed on biological replicates to mitigate pseudoreplication. Briefly, images were captured using 10 μm z-stacked acquisition settings sampled at Nyquist. Raw files were airyscan processed using standard settings (Zeiss) and converted to the imaris file format using the Imaris File Converter (Bitplane). Machine learning-based image analysis was performed on AmyTracker 680 signals and then applied evenly across the entire dataset (i.e., 6 biological replicates; 20-30 technical replicates per sample; 0-150 3D segmented objects per image; Bitplane). The average morphological summary per biological replicate was then identified and compared per age group.

### Transmission electron microscopy (TEM)

The Reynolds Lead Citrate Protocol was used for TEM sample preparation. Briefly, samples were fixed in 2% paraformaldehyde, 2.5% glutaraldehyde, 0.1M sodium cacodylate (pH 7.4) at 4°C with agitation and post-fixed with 1% osmium tetroxide. Following post-fixation, samples were dehydrated in serial concentrations of ethanol, finished with propylene oxide, and embedded in epon/araldiate resin. Ultrathin 100 nm sections were collected, post-stained with lead citrate, and visualized using a Hitachi H-7600 transmission electron microscope (Hitachi).

### Biochemical fractionation

Detergent-soluble and detergent-insoluble proteins were isolated as described previously ^5,42^. Approximately 20 mg powdered gastrocnemius was homogenized in 1% Triton X-100 buffer with protease/phosphatase inhibitors. Aliquots containing 600 µg protein were centrifuged at 17,000g for 10 min at 4 °C. Supernatants were collected, and pellets were solubilized in 2% SDS buffer by sonication. Bulk fraction content was quantified by GelCode Blue staining of SDS–PAGE gels and lane-intensity integration.

### Proteomics analyses

Aliquots (5 µL) of the soluble and insoluble fractions described above were used for proteomic analysis. BSA (1 µg) was added to each sample as an internal standard, and peptides were prepared by in-gel digest as previously described ^43^. For analysis by liquid chromatography mass spectrometry, peptides were separated and measured with a ThermoScientific Orbitrap Exploris 480 operating in the data-independent acquisition (DIA) mode. MS1 spectra were collected at a resolution of 15,000 with a scan range of 300-1100 m/z. MS2 spectra were collected at a resolution of 30,000 using 20 m/z isolation windows from 350-950 m/z with no overlap.

Proteins were identified and quantified from the LCMS data using DIANN (version 2.5.1 Academia). Protein sequences were downloaded from UniProt ^44^, and default settings without normalization were used for identification using a predicted spectral library. Soluble and insoluble protein-group abundances were analyzed separately. Values were log₂-transformed without additional downstream normalization or imputation. Proteins required detection in at least 50% of samples in both age groups or each old-treatment group, pooling sexes for eligibility. Principal component analysis used centered and scaled variables and the NIPALS algorithm. Heatmaps displayed protein-wise Z scores.

Differential abundance was analyzed using limma age-by-sex or treatment-by-sex cell-means models and empirical Bayes moderation with an intensity-dependent variance trend. Contrasts estimated age, sex, intervention and interaction effects, including within-sex effects. Sex-averaged effects weighted females and males equally. Moderated omnibus F-tests assessed intervention effects and their sex interactions. Benjamini– Hochberg adjustment was applied across proteins within each contrast. Adjusted P ≤ 0.05 defined FDR-significant changes, whereas raw p-value refers to nominal P ≤ 0.05. No fold-change threshold was imposed.

Gene Ontology Biological Process enrichment used clusterProfiler and org.Mm.eg.db, with increased and decreased protein sets analyzed separately against the corresponding filtered, Entrez-mapped background. Nominal-P and FDR-selected sets were evaluated separately, with Benjamini–Hochberg adjustment of GO terms (P cutoff 0.05; q cutoff 0.25). Curated protein-folding and chaperone modules used the corresponding proteome-wide statistics.

### Protein partitioning and insoluble proteome composition

Soluble and insoluble measurements were matched by protein group and animal and corrected for fraction volume and aliquot dilution. Partitioning was log₂(corrected insoluble/corrected soluble abundance), requiring positive measurements in both fractions and at least two valid animal pairs per age or treatment group. Ratios were analyzed using the limma models described above. Soluble-versus-total abundance plots were used only for descriptive purposes.

Proteins were assigned to mutually exclusive GO Cellular Component bins using descendant terms and a predefined priority order. Unassigned proteins were excluded. Composition analyses retained proteins detected in at least two samples in either age group or any treatment group. Per-animal composition was calculated from summed component intensities or detected protein counts relative to all retained, assigned proteins. Missing values contributed zero. Arcsine-square-root-transformed proportions were tested using pooled and factorial models, Type III tests and planned contrasts, with Benjamini–Hochberg adjustment across components. Effects were expressed as percentage-point or relative changes, with bootstrap age-effect intervals and model-based treatment-effect intervals. Summed component intensities were analyzed separately on the log₂ scale, with a half-minimum-positive pseudocount for treatment analyses. Dominant-protein rankings and abundance profiles were descriptive.

### Detection prevalence

Proteins with successful, positive quantification by DIANN with standard quality filters were considered ‘detected’. Two-sided Fisher’s exact tests compared detection frequencies between ages or each intervention and control, pooled and within sex, with Benjamini–Hochberg adjustment within comparison. Predominant or reduced detection contrasted at least two detections against at most one by group. Shared and intervention-specific loss patterns were count-defined.

### Age-shift reversal

Age and intervention effects were matched by protein group and sex. Significant opposite-direction effects defined reversal, and significant same-direction effects defined further age-associated change. Classifications used nominal and adjusted P ≤ 0.05 separately. The extent of reversal was calculated by dividing the intervention-associated change by the age-associated change and reversing the sign, such that positive values indicate changes opposing the effects of aging.

### Aggregation propensity

Precomputed AGGRESCAN Na4vSS, TANGO-derived and WALTZ amyloid-percentage scores were matched to cleaned protein accessions. The TANGO-derived field was divided by sequence length. Mean percentile ranks provided a consensus ranking. Age analyses used adult-control soluble proteins and intervention analyses used old-control insoluble proteins, requiring at least two baseline detections across sexes. Sex-specific effects required at least two observations per comparison group. Abundance effects were log₂ ratios of arithmetic means. Partitioning effects were differences in mean log₂ ratios.

Pearson correlations related propensity to abundance and partitioning changes, with Benjamini– Hochberg adjustment across metrics within comparison, sex and endpoint. Component distributions used Kruskal–Wallis and Holm-adjusted Dunn tests. Component-specific correlations and component-adjusted regressions assessed compartment effects. Nested F-tests assessed propensity-by-component interactions. Exploratory soluble-abundance correlations reported nominal P values. Global accession-score distributions and animal-level abundance-weighted scores used planned treatment-by-sex comparisons: HC3-robust linear models for Na4vSS and aligned rank transform models for TANGO and WALTZ, with Holm adjustment across interventions.

To assess whether proteins showing age- or treatment-associated changes in insoluble detection had greater intrinsic aggregation propensity, AGGRESCAN, TANGO, and WALTZ scores were converted to percentiles relative to the old-control insoluble proteome, separately by sex. Propensity distributions were compared with those of proteins consistently detected across the relevant groups using one-sided Wilcoxon rank-sum tests. Enrichment among proteins above the 75th propensity percentile was assessed using one-sided Fisher’s exact tests. Analyses were repeated for proteins showing nominally significant detection changes (P < 0.05). Benjamini–Hochberg correction was applied within each sex and analysis subset, separately for each test.

### Stable-isotope labeling and protein turnover

Mice received an intraperitoneal bolus of 99% D₂O equivalent to 5% of the body-water pool, followed by 8% D₂O drinking water for 0–60 days ^45^. Body water was recovered from 50 µl plasma overnight at 80 °C, diluted 1:300 and measured using a Los Gatos Research isotope analyzer against D₂O standards. Myofibrillar, mitochondrial and cytosolic fractions were isolated by differential centrifugation, hydrolyzed and analyzed for protein-bound alanine enrichment by gas chromatography-mass spectrometry ^45–48^. To determine the protein turnover of the different subcellular fractions, we calculated the rise to plateau of fraction new over the time-course according to our previously published studies ^5,45^. This analysis calculates the synthesis rate (*k*) over the 60-day period using a one-phase association. The plateau fraction new value (*p*) obtained indicates the fraction of a protein pool that is actively turning over, with 1.0 equal to 100% of the protein pool.

Peptide M+0 and M+1 isotopologue intensities were extracted from soluble and insoluble fractions using Skyline after DIA-NN quality filtering. M+0 depletion was calculated using observed and theoretical unlabeled and fully labeled isotope envelopes. Fraction new was then calculated using peptide composition, the effective number of labeling hydrogens, and animal-specific body-water enrichment. Values outside of 0-1 by at most 0.10 were clipped to the nearest bound; more extreme or invalid values were excluded. Peptides were retained only if they had valid measurements in all analyzed samples within a given fraction. Animal-level means were analyzed separately at 30, 45, and 60 days using mixed-effects models with age × fraction or treatment × fraction fixed effects and a random intercept for animal. Planned intervention contrasts were adjusted using the Holm method.

### Statistical analyses

Statistical analyses were performed using R v4.5.0 and GraphPad Prism v11.0.2. Individual animals were considered independent biological replicates for animal-level outcomes, whereas individual protein groups were the analytical units for protein-level aggregation-propensity analyses and correlations. Microscopy-based age comparisons were performed using two-sided, unpaired Student’s t-tests assuming equal variances, with individual mice considered independent biological replicates (n = 3 per age group). Measurements from multiple microscopy images were aggregated at the mouse level before statistical testing. Unless otherwise specified, statistical significance was defined as P ≤ 0.05, with Benjamini–Hochberg or Holm adjustments for multiple comparisons applied as described in the respective sections. Data are presented as means ± s.e.m. or estimated effects with 95% confidence intervals, unless otherwise indicated. Sample sizes, statistical tests and significance thresholds are specified in the corresponding figure legends.

## Acknowledgments

We thank Frederick F. Peelor III for sample preparation and analysis of GC-MS data, and Ben Fowler, Julie Crane, and Justin Willige for the sample preparation and processing of transmission electron microscopy samples and troubleshooting of light microscope image acquisition and analysis of the images generated for this manuscript. This work was supported by grants from the NIH (R01AG074551 to B.F.M.), the U.S. Department of Veterans Affairs (IK6BX007150 to B.F.M.), and the Presbyterian Health Foundation Collaborative Science Grant (to. Z.R.H). The funders had no role in study design, data collection and analysis, decision to publish or preparation of the manuscript.

## Author contributions

P.H.C.M., B.F.M., and Z.R.H. conceived and designed the study. P.H.C.M. performed the animal experiments, including exercise and rapamycin interventions, tissue collection and detergent-solubility experiments, and conducted proteomic, bioinformatic and statistical analyses. Z.R.H. led the microscopy experiments, with contributions from T.G., X.R. and J.P. T.J.L. performed proteomic sample preparation and LC–MS analyses. M.E.T. performed the aggregation-propensity, fraction-new analyses, and analyzed data. B.F.M. supervised the study and secured funding. P.H.C.M., Z.R.H., and B.F.M. performed manuscript writing, reviewing and editing. All authors reviewed and approved the final manuscript.

## Competing interests

All authors declare no competing interests.

## Use generative AI

OpenAI’s ChatGPT was used to assist with R code development and troubleshooting, language editing, text-style refinement, and figure conceptualization. All scientific interpretations, statistical analyses and AI-assisted outputs were critically reviewed and validated by the authors, who take full responsibility for the accuracy and integrity of the manuscript.

## Data availability

The data supporting the findings of this study are provided in the manuscript, Extended Data and Supplementary Information. The mass spectrometry proteomics datasets generated in this study will be deposited in the ProteomeXchange Consortium via the PRIDE partner repository and made publicly available before publication. Underlying source data and additional information necessary to reproduce the analyses are available from the corresponding author upon reasonable request during peer review.

## Notes

### Competing Interest Statement

The authors have declared no competing interest.

