## Extended Data Figures for "Skeletal muscle aging drives sex- and compartment-specific protein insolubility that is modifiable by proteostasis-directed treatments"

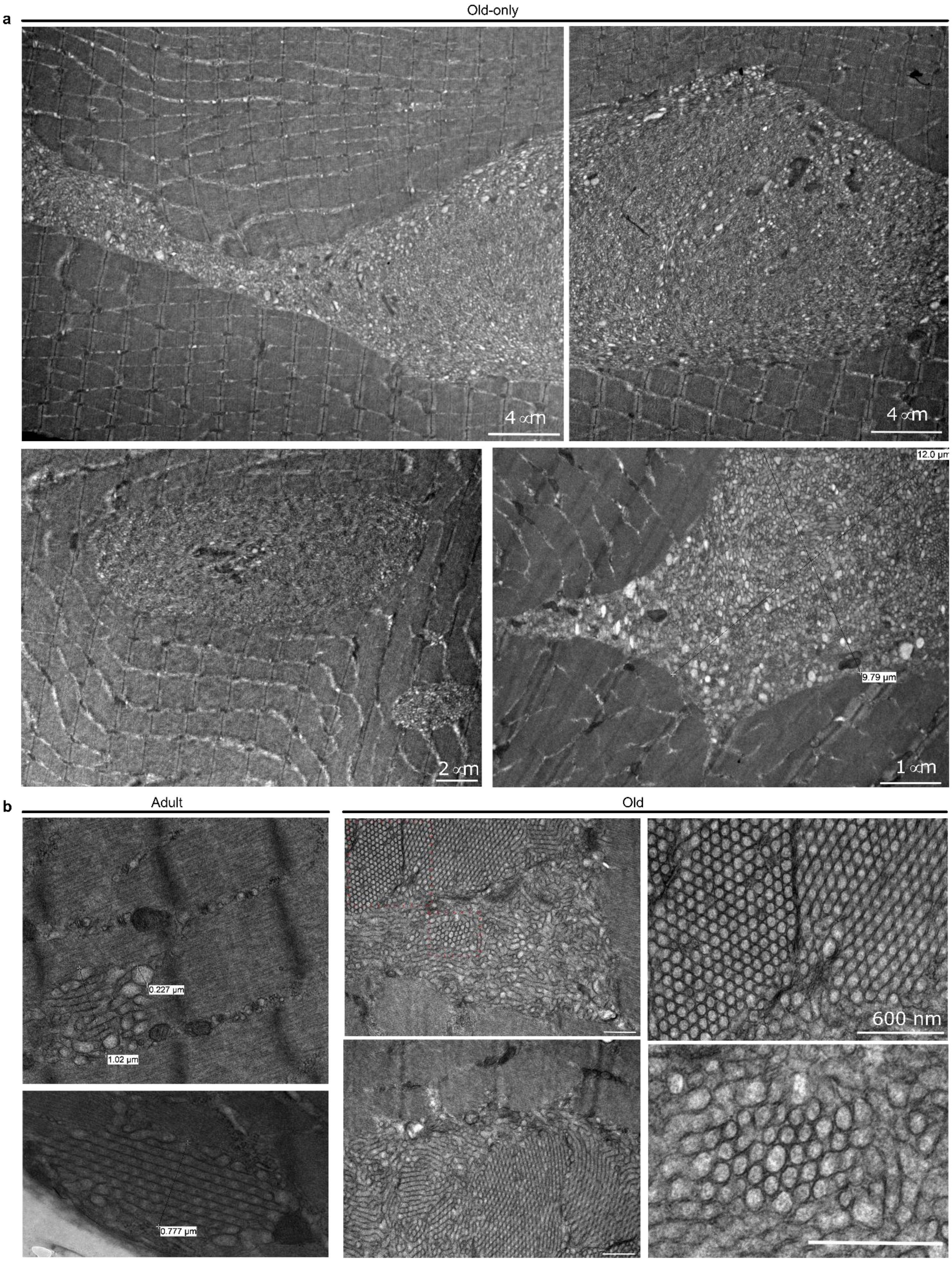


**Extended Data Fig. 1 | Ultrastructural differences between age-associated aggresome-like structures and tubular aggregates. a**, Representative transmission electron microscopy (TEM) images showing large, irregularly shaped aggresome-like structures within aged skeletal muscle fibers. The upper images show adjoining regions of an extensive inclusion characterized by heterogeneous accumulation of vesicular and membranous structures and disruption of the surrounding myofibrillar organization. The lower images show additional examples of these structures at different magnifications, highlighting their variable morphology and extensive occupation of the intracellular space. **b**,

Representative TEM images of tubular aggregates observed in adult and old skeletal muscle, showing densely packed, regularly organized tubular membranes in longitudinal and transverse orientations. In contrast to the heterogeneous architecture of aggresome-like structures, tubular aggregates show a more uniform and ordered ultrastructural organization. Scale bars are indicated in each micrograph.

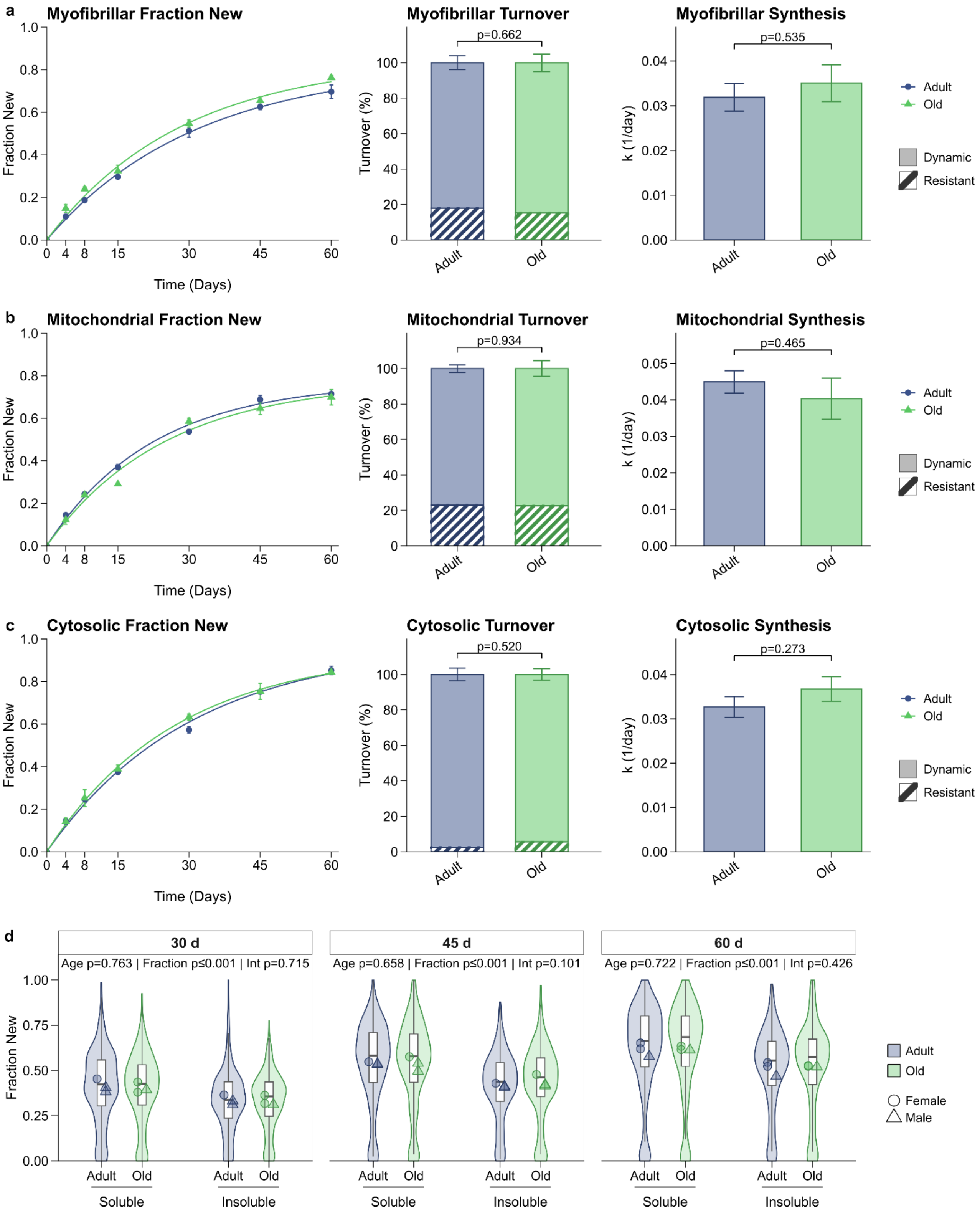

a
Myofibrillar Fraction New
Fraction New
Time (Days)
Myofibrillar Turnover
p=0.662
Turnover (%)
Adult
Old
Myofibrillar Synthesis
p=0.535
k (1/day)
Adult
Old
Dynamic
Resistant
b
Mitochondrial Fraction New
Mitochondrial Turnover
p=0.934
Mitochondrial Synthesis
p=0.465
c
Cytosolic Fraction New
Cytosolic Turnover
p=0.520
Cytosolic Synthesis
p=0.273
d
30 d
Age p=0.763 | Fraction p≤0.001 | Int p=0.715
45 d
Age p=0.658 | Fraction p≤0.001 | Int p=0.101
60 d
Age p=0.722 | Fraction p≤0.001 | Int p=0.426
Soluble
Insoluble
Female
Male

**Extended Data Fig. 2 | Protein synthesis and turnover in adult and aged skeletal muscle. a–c,** Deuterium oxide labeling time courses for myofibrillar (**a**), mitochondrial (**b**) and cytosolic (**c**) protein fractions from adult and old gastrocnemius muscle. Left, fraction of newly synthesized protein (fraction new) during up to 60 days of labeling, with separate one-phase association curves. Middle, dynamic (Plateau) and resistant (1 − Plateau) protein-pool proportions. Right, fitted synthesis rate constant, k ($day^{-1}$). **d,** Fraction new in detergent-soluble and detergent-insoluble fractions after 30, 45 and 60 days of labeling. Violins and boxplots show peptide-level values; overlaid circles and triangles represent individual female and male mouse means, respectively. In **a–c**, the fits included 19/16, 18/16 and 17/17 observations for adult/old groups in the myofibrillar, mitochondrial and cytosolic fractions, respectively. These are fit-observation counts. In **d**, n = 3 mice per age at each duration, with paired fractions from each mouse; retained peptide sets comprised 144 soluble and 110 insoluble peptide species. Time-course values are means ± s.e.m.; error bars for pool proportions and k show s.e. of the fitted plateau and k, respectively. Curves were fitted as $FN(t) = Plateau \times (1 - e^{-kt})$, with the initial value fixed at zero. Age comparisons of fitted parameters used two-sided, equal-variance unpaired Student's t-tests calculated from parameter estimates, their s.e. and fit-observation counts. In **d**, boxes show medians and interquartile ranges; whiskers extend to the most extreme values within 1.5 times the interquartile range. Animal-level means were analyzed using separate mixed models at each duration, with age × fraction fixed effects and a random mouse intercept. P values are from Type III F-tests. Displayed P values are unadjusted.

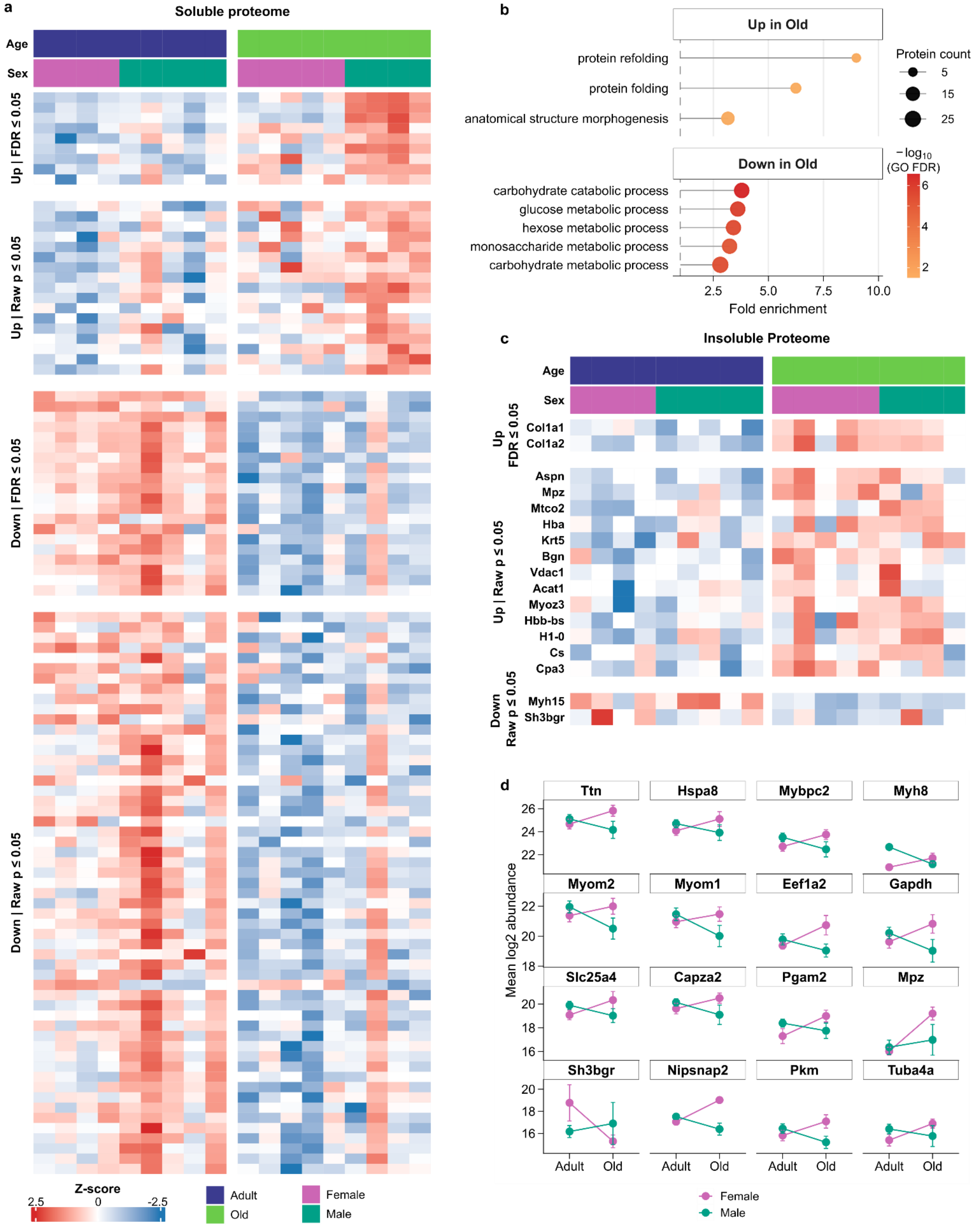

a
Soluble proteome
Age
Sex
Up | FDR ≤ 0.05
Up | Raw p ≤ 0.05
Down | FDR ≤ 0.05
Down | Raw p ≤ 0.05
Z-score
2.5
0
-2.5
Adult
Old
Female
Male
b
Up in Old
protein refolding
protein folding
anatomical structure morphogenesis
Down in Old
carbohydrate catabolic process
glucose metabolic process
hexose metabolic process
monosaccharide metabolic process
carbohydrate metabolic process
Protein count
5
15
25
−log10 (GO FDR)
6
4
2
2.5
5.0
7.5
10.0
Fold enrichment
c
Insoluble Proteome
Age
Sex
Up FDR ≤ 0.05
Col1a1
Col1a2
Up | Raw p ≤ 0.05
Aspn
Mpz
Mtco2
Hba
Krt5
Bgn
Vdac1
Acat1
Myoz3
Hbb-bs
H1-0
Cs
Cpa3
Down Raw p ≤ 0.05
Myh15
Sh3bgr
d
Ttn
Hspa8
Mybpc2
Myh8
Myom2
Myom1
Eef1a2
Gapdh
Slc25a4
Capza2
Pgam2
Mpz
Sh3bgr
Nipsnap2
Pkm
Tuba4a
Mean log2 abundance
26
24
22
20
18
16
Adult
Old
Female
Male

**Extended Data Fig. 3 | Age-associated changes in soluble and insoluble protein abundance. a,** Heatmap of soluble proteins showing nominally significant age effects ($P \leq 0.05$), grouped by direction of change and false discovery rate (FDR) significance. Columns represent individual mice, annotated by age and sex; values are row Z-scores of $\log_2$ abundance, with the color scale clipped at ±2.5. **b,** Gene Ontology (GO) biological-process enrichment among proteins increased or decreased with age, selected at nominal age-effect $P \leq 0.05$ (26 increased and 75 decreased proteins before Entrez mapping). Up to five terms with the lowest adjusted P values are shown per direction. Position indicates fold enrichment, circle size indicates protein count and color indicates $-\log_{10}$(GO FDR). **c,** Corresponding heatmap of age-associated changes in the insoluble proteome. **d,** Insoluble protein abundances for proteins selected at nominal age × sex interaction $P \leq 0.05$, without a fold-change cutoff. For both fractions, n = 4 adult female, 5 adult male, 5 old female and 4 old male mice. The analyses tested 477 soluble and 148 insoluble proteins; valid observations vary by protein. Values in **d** are means ± s.e.m. Protein-level effects were estimated using age × sex linear models with intensity-dependent empirical Bayes moderation (limma), averaging female and male age effects equally. Tests were two-sided, with Benjamini–Hochberg correction across proteins separately for each contrast. GO enrichment (**b**) used one-sided hypergeometric tests in clusterProfiler, with all filtered, Entrez-mapped soluble proteins as background and Benjamini–Hochberg correction across GO terms within each direction. Exact P and FDR values are shown where indicated.

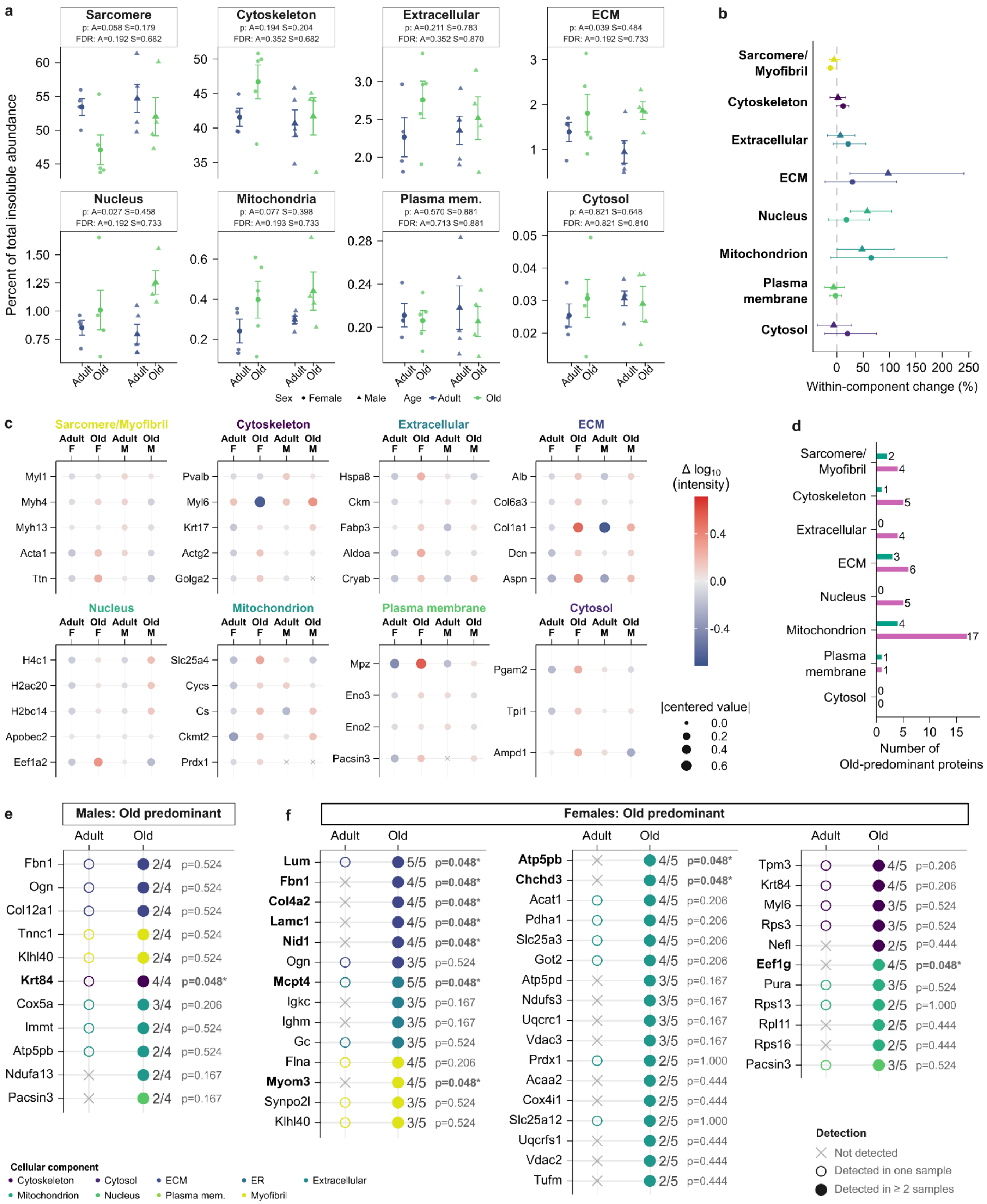

a
Sarcomere
p: A=0.058 S=0.179
FDR: A=0.192 S=0.682
Cytoskeleton
p: A=0.194 S=0.204
FDR: A=0.352 S=0.682
Extracellular
p: A=0.211 S=0.783
FDR: A=0.352 S=0.870
ECM
p: A=0.039 S=0.484
FDR: A=0.192 S=0.733
Nucleus
p: A=0.027 S=0.458
FDR: A=0.192 S=0.733
Mitochondria
p: A=0.077 S=0.398
FDR: A=0.193 S=0.733
Plasma mem.
p: A=0.570 S=0.881
FDR: A=0.713 S=0.881
Cytosol
p: A=0.821 S=0.648
FDR: A=0.821 S=0.810
Percent of total insoluble abundance
Adult Old Adult Old
Sex • Female ▲ Male Age Adult Old
b
Sarcomere/ Myofibril
Cytoskeleton
Extracellular
ECM
Nucleus
Mitochondrion
Plasma membrane
Cytosol
0 50 100 150 200 250
Within-component change (%)
c
Sarcomere/Myofibril
Adult F Old F Adult M Old M
Myl1
Myh4
Myh13
Acta1
Ttn
Cytoskeleton
Pvalb
Myl6
Krt17
Actg2
Golga2
Extracellular
Hspa8
Ckm
Fabp3
Aldoa
Cryab
ECM
Alb
Col6a3
Col1a1
Dcn
Aspn
Nucleus
H4c1
H2ac20
H2bc14
Apobec2
Eef1a2
Mitochondrion
Slc25a4
Cycs
Cs
Ckmt2
Prdx1
Plasma membrane
Mpz
Eno3
Eno2
Pacsin3
Cytosol
Pgam2
Tpi1
Ampd1
Δ log10 (intensity)
0.4
0.0
-0.4
|centered value|
0.0
0.2
0.4
0.6
d
Sarcomere/ Myofibril 2 4
Cytoskeleton 1 5
Extracellular 0 4
ECM 3 6
Nucleus 0 5
Mitochondrion 4 17
Plasma membrane 1 1
Cytosol 0 0
0 5 10 15
Number of Old-predominant proteins
e
Males: Old predominant
Adult Old
Fbn1 2/4 p=0.524
Ogn 2/4 p=0.524
Col12a1 2/4 p=0.524
Tnnc1 2/4 p=0.524
Klhl40 2/4 p=0.524
Krt84 4/4 p=0.048*
Cox5a 3/4 p=0.206
Immt 2/4 p=0.524
Atp5pb 2/4 p=0.524
Ndufa13 2/4 p=0.167
Pacsin3 2/4 p=0.167
f
Females: Old predominant
Adult Old
Lum 5/5 p=0.048*
Fbn1 4/5 p=0.048*
Col4a2 4/5 p=0.048*
Lamc1 4/5 p=0.048*
Nid1 4/5 p=0.048*
Ogn 3/5 p=0.524
Mcpt4 5/5 p=0.048*
Igkc 3/5 p=0.167
Ighm 3/5 p=0.167
Gc 3/5 p=0.524
Flna 4/5 p=0.206
Myom3 4/5 p=0.048*
Synpo2l 3/5 p=0.524
Klhl40 3/5 p=0.524
Atp5pb 4/5 p=0.048*
Chchd3 4/5 p=0.048*
Acat1 4/5 p=0.206
Pdha1 4/5 p=0.206
Slc25a3 4/5 p=0.206
Got2 4/5 p=0.206
Atp5pd 3/5 p=0.167
Ndufs3 3/5 p=0.167
Uqcrc1 3/5 p=0.167
Vdac3 3/5 p=0.167
Prdx1 2/5 p=1.000
Acaa2 2/5 p=0.444
Cox4i1 2/5 p=0.444
Slc25a12 2/5 p=1.000
Uqcrfs1 2/5 p=0.444
Vdac2 2/5 p=0.444
Tufm 2/5 p=0.444
Tpm3 4/5 p=0.206
Krt84 4/5 p=0.206
Myl6 3/5 p=0.524
Rps3 3/5 p=0.524
Nefl 2/5 p=0.444
Eef1g 4/5 p=0.048*
Pura 3/5 p=0.524
Rps13 2/5 p=1.000
Rpl11 2/5 p=0.444
Rps16 2/5 p=0.444
Pacsin3 3/5 p=0.524
Cellular component
Cytoskeleton Cytosol ECM ER Extracellular
Mitochondrion Nucleus Plasma mem. Myofibril
Detection
Not detected
Detected in one sample
Detected in ≥ 2 samples

**Extended Data Fig. 4 | Cellular composition and detection patterns of the aging insoluble proteome. a,** Contributions of cellular components to insoluble protein abundance in adult and old female and male mice. Percentages use all GO-assigned, filtered proteins as the denominator, including components omitted from the display. **b,** Sex-specific relative changes in component representation, calculated as 100 × (mean old fraction − mean adult fraction)/mean adult fraction. **c,** Five proteins with the highest pooled old-group mean abundance per component, where available. Color shows $\log_{10}$(mean intensity among positive detections) centered within each protein across the four age × sex groups; size shows the absolute centered value, and crosses indicate unavailable values. **d,** Numbers of old-predominant proteins by component and sex. **e,f,** Detection patterns in male (**e**) and female (**f**) mice. Old-predominant proteins have ≥2 old and ≤1 adult detections within sex. Symbols distinguish zero, one and ≥2 detections; fractions show samples with detection/total samples. For all analyses, n = 4 adult female, 5 adult male, 5 old female and 4 old male mice. The detection analysis identified 42 old-predominant proteins in females and 11 in males. Composition analyses retained proteins detected in at least two adult or two old samples, pooling sexes within age. In **a**, symbols represent individual mice and values are means ± s.e.m. Component fractions were analyzed using age × sex linear models after arcsine-square-root transformation, with Type III F-tests and Benjamini–Hochberg correction across components separately for each effect. Error bars in **b** show percentile bootstrap 95% confidence intervals from 5,000 within-age resamples per component and sex. Detection prevalence (**e,f**) was compared using two-sided Fisher's exact tests; displayed P values are unadjusted, *P ≤ 0.05.

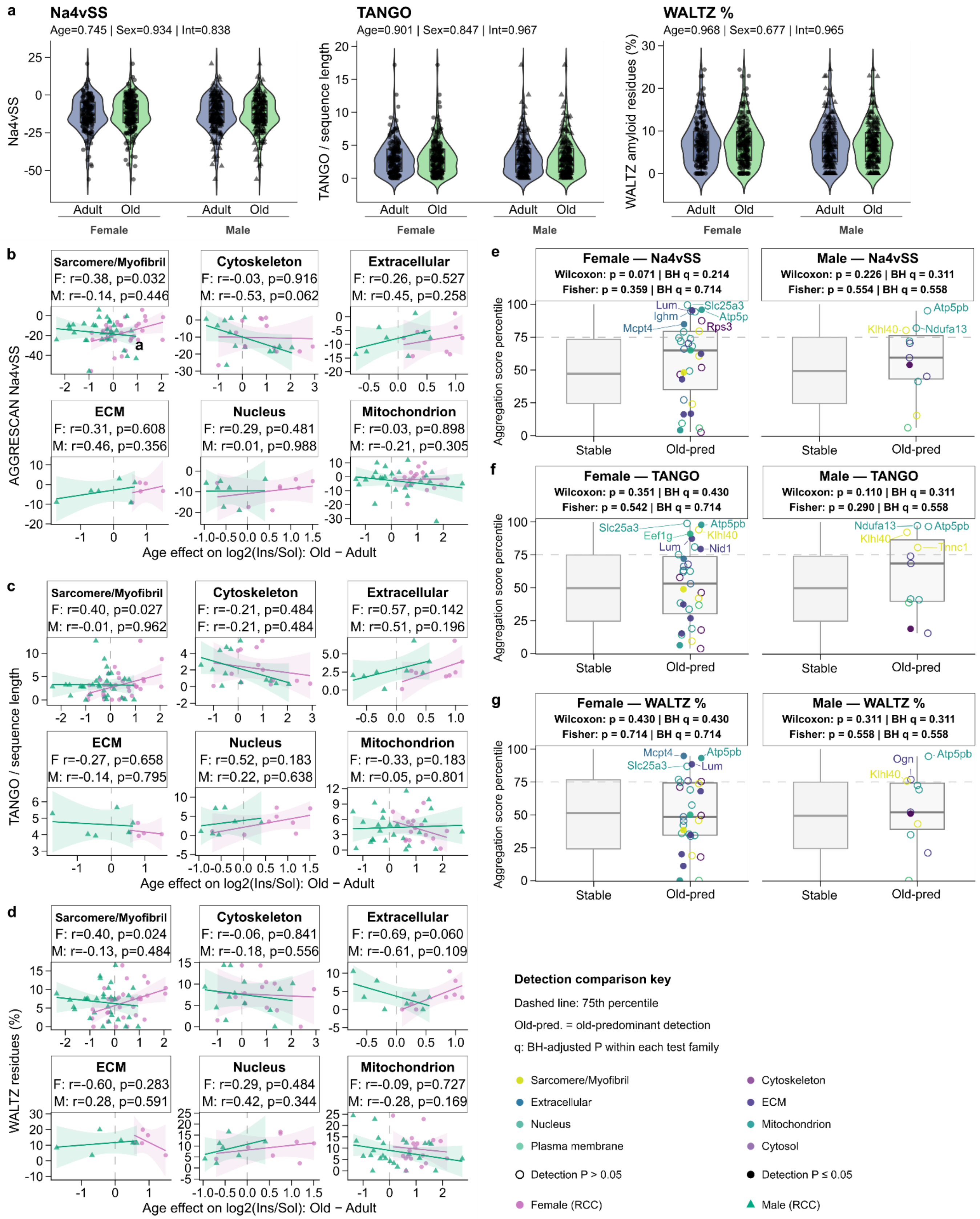
a
Na4vSS
Age=0.745 | Sex=0.934 | Int=0.838
TANGO
Age=0.901 | Sex=0.847 | Int=0.967
WALTZ %
Age=0.968 | Sex=0.677 | Int=0.965
Na4vSS
TANGO / sequence length
WALTZ amyloid residues (%)
Adult Old
Female
Male
b
Sarcomere/Myofibril
F: r=0.38, p=0.032
M: r=-0.14, p=0.446
Cytoskeleton
F: r=-0.03, p=0.916
M: r=-0.53, p=0.062
Extracellular
F: r=0.26, p=0.527
M: r=0.45, p=0.258
ECM
F: r=0.31, p=0.608
M: r=0.46, p=0.356
Nucleus
F: r=0.29, p=0.481
M: r=0.01, p=0.988
Mitochondrion
F: r=0.03, p=0.898
M: r=-0.21, p=0.305
AGGRESCAN Na4vSS
Age effect on log2(Ins/Sol): Old − Adult
c
Sarcomere/Myofibril
F: r=0.40, p=0.027
M: r=-0.01, p=0.962
Cytoskeleton
F: r=-0.21, p=0.484
F: r=-0.21, p=0.484
Extracellular
F: r=0.57, p=0.142
M: r=0.51, p=0.196
ECM
F r=-0.27, p=0.658
M: r=-0.14, p=0.795
Nucleus
F: r=0.52, p=0.183
M: r=0.22, p=0.638
Mitochondrion
F: r=-0.33, p=0.183
M: r=0.05, p=0.801
TANGO / sequence length
Age effect on log2(Ins/Sol): Old − Adult
d
Sarcomere/Myofibril
F: r=0.40, p=0.024
M: r=-0.13, p=0.484
Cytoskeleton
F: r=-0.06, p=0.841
M: r=-0.18, p=0.556
Extracellular
F: r=0.69, p=0.060
M: r=-0.61, p=0.109
ECM
F: r=-0.60, p=0.283
M: r=0.28, p=0.591
Nucleus
F: r=0.29, p=0.484
M: r=0.42, p=0.344
Mitochondrion
F: r=-0.09, p=0.727
M: r=-0.28, p=0.169
WALTZ residues (%)
Age effect on log2(Ins/Sol): Old − Adult
e
Female — Na4vSS
Wilcoxon: p = 0.071 | BH q = 0.214
Fisher: p = 0.359 | BH q = 0.714
Male — Na4vSS
Wilcoxon: p = 0.226 | BH q = 0.311
Fisher: p = 0.554 | BH q = 0.558
Aggregation score percentile
Lum Slc25a3 Ighm Atp5p Mcpt4 Rps3
Atp5pb Klhl40 Ndufa13
Stable Old-pred
f
Female — TANGO
Wilcoxon: p = 0.351 | BH q = 0.430
Fisher: p = 0.542 | BH q = 0.714
Male — TANGO
Wilcoxon: p = 0.110 | BH q = 0.311
Fisher: p = 0.290 | BH q = 0.558
Slc25a3 Atp5pb Eef1g Klhl40 Lum Nid1
Ndufa13 Atp5pb Klhl40 Tnnc1
g
Female — WALTZ %
Wilcoxon: p = 0.430 | BH q = 0.430
Fisher: p = 0.714 | BH q = 0.714
Male — WALTZ %
Wilcoxon: p = 0.311 | BH q = 0.311
Fisher: p = 0.558 | BH q = 0.558
Mcpt4 Atp5pb Slc25a3 Lum
Ogn Atp5pb Klhl40
Detection comparison key
Dashed line: 75th percentile
Old-pred. = old-predominant detection
q: BH-adjusted P within each test family
Sarcomere/Myofibril
Cytoskeleton
Extracellular
ECM
Nucleus
Mitochondrion
Plasma membrane
Cytosol
Detection P > 0.05
Detection P ≤ 0.05
Female (RCC)
Male (RCC)

**Extended Data Fig. 5 | Intrinsic aggregation propensity in relation to aging-associated protein partitioning and detection. a,** Unweighted distributions of AGGRESCAN Na4vSS, sequence-length-normalized TANGO scores and WALTZ-predicted amyloidogenic residues (%) among insoluble proteins detected in ≥2 samples within each age × sex group. **b–d,** Relationships between the age effect on the volume-corrected $\log_2$ insoluble-to-soluble protein ratio (Old − Adult) and Na4vSS (**b**), TANGO (**c**) or WALTZ (**d**) within cellular components, using the adult-soluble starting protein pool. Circles and triangles represent females and males, respectively; solid lines show linear regressions with 95% confidence intervals. **e–g,** Na4vSS (**e**), TANGO (**f**) and WALTZ (**g**) score percentiles for stable and old-predominant proteins. Old-predominant proteins have ≥2 old and ≤1 adult detections; stable proteins have ≥2 detections at both ages, within sex. Percentiles reference all scored proteins with ≥2 old insoluble detections within sex. Colors indicate components; filled symbols indicate nominal detection $P < 0.05$, and open symbols indicate other candidates. Proteomic measurements were obtained from n = 4 adult female, 5 adult male, 5 old female and 4 old male mice. Each propensity point represents a protein rather than an independent biological replicate. In **e–g**, old-predominant protein counts were 31 females and 11 males per metric; stable-protein counts were 161/172 for Na4vSS and 160/171 for TANGO and WALTZ, in females/males, respectively. Na4vSS distributions (**a**) were analyzed using age × sex linear models with Type III HC3-robust F-tests; TANGO and WALTZ used aligned-rank-transform factorial models. Displayed P values are unadjusted. Associations in **b–d** used two-sided Pearson correlation tests; components are shown when both sex-specific tests include ≥5 proteins. Headers show r and raw P values. In **e–g**, boxes show medians and interquartile ranges, with whiskers extending to the most extreme values within 1.5 times the interquartile range. One-sided Wilcoxon rank-sum and Fisher's exact tests assessed higher scores and enrichment at or above the 75th percentile, respectively. Benjamini–Hochberg correction was applied across the three metrics within sex, separately for each test family.

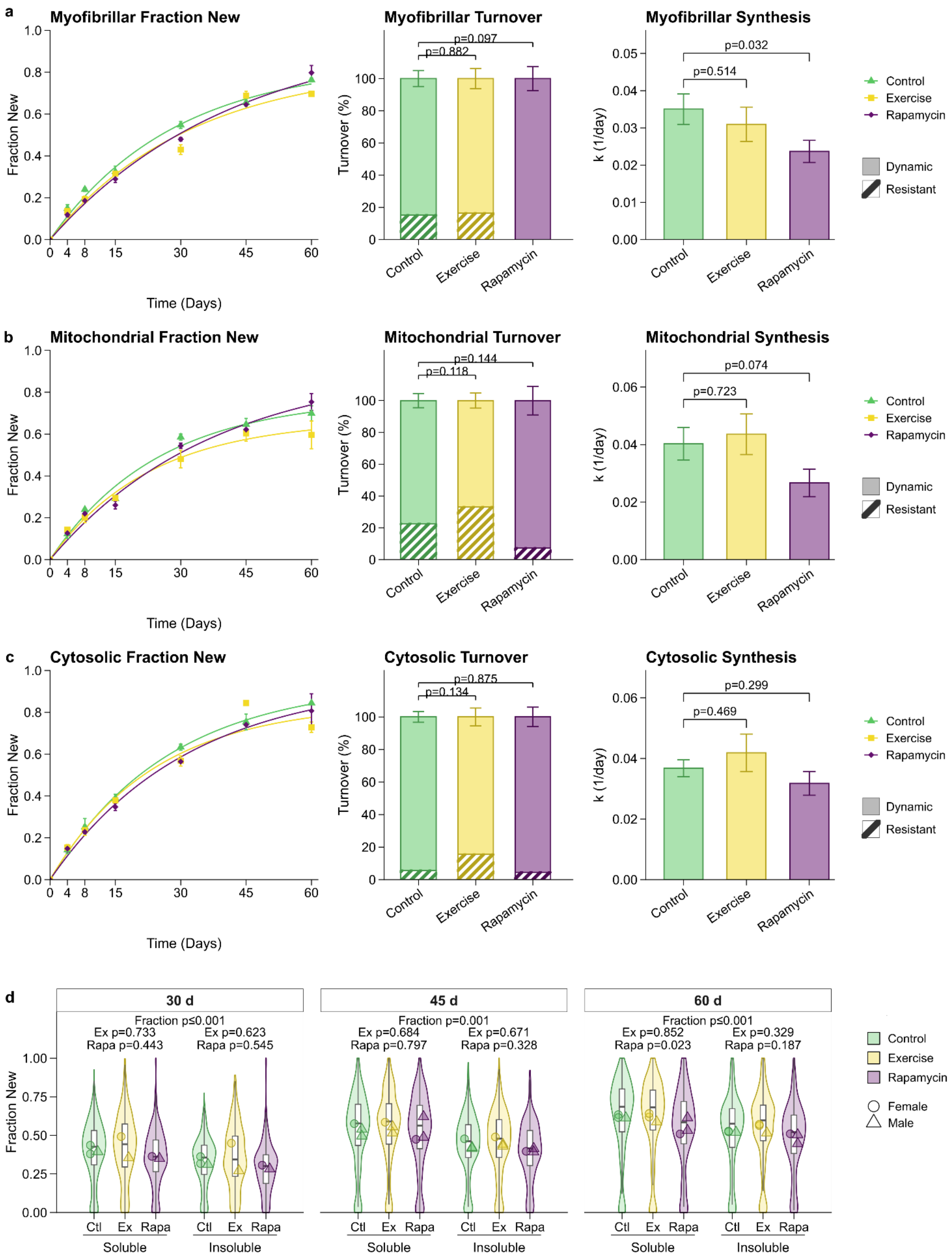


**Extended Data Fig. 6 | Protein synthesis and turnover following exercise or rapamycin treatment in aged skeletal muscle. a–c,** Deuterium oxide labeling time courses for myofibrillar (**a**), mitochondrial (**b**) and cytosolic (**c**) protein fractions from old control (Ctl), exercise (Ex) and rapamycin-treated (Rapa) mice. Left, fraction new with separate one-phase association curves. Middle, dynamic (Plateau) and resistant (1 − Plateau) pool proportions. Right, fitted synthesis rate constant, k ($day^{-1}$). **d,** Fraction new in detergent-soluble and detergent-insoluble fractions after 30, 45 and 60 days of labeling. Violins and boxplots show peptide-level values; overlaid circles and triangles represent individual female and male mouse means, respectively. In **a–c**, included fit observations for control/exercise/rapamycin were 16/17/16 (myofibrillar), 16/18/17 (mitochondrial) and 17/18/16 (cytosolic). In **d**, n = 3/3/3 control mice, 2/3/3 exercise mice and 2/3/3 rapamycin mice at 30/45/60 days, respectively, with paired fractions from each mouse. Retained sets comprised 144 soluble and 110 insoluble peptide species. Time-course values are means ± s.e.m.; error bars for pool proportions and k show s.e. of the fitted plateau and k, respectively. Curves were fitted as $FN(t) = \mathrm{Plateau} \times (1 - e^{-kt})$, with the initial value fixed at zero. Fitted parameters were compared using two prespecified treatment-versus-control, two-sided, equal-variance unpaired Student's t-tests calculated from parameter estimates, their s.e. and fit-observation counts. In **d**, boxes show medians and interquartile ranges; whiskers extend to the most extreme values within 1.5 times the interquartile range. Animal means were analyzed using separate mixed models at each duration, with treatment × fraction fixed effects and a random mouse intercept. The displayed fraction comparison averages across treatments. Displayed P values are unadjusted planned comparisons.

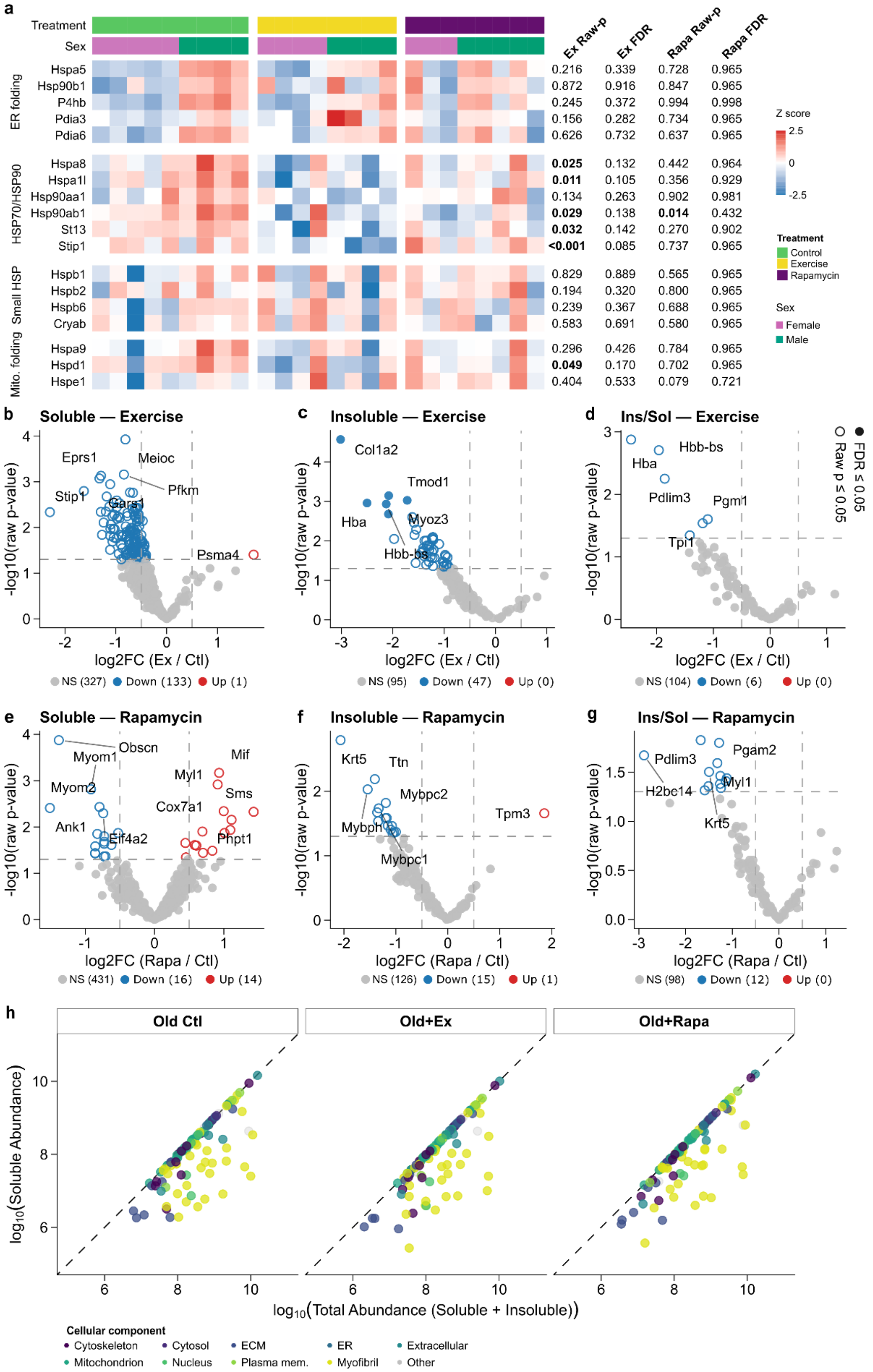
a
Treatment
Sex
Ex Raw-p
Ex FDR
Rapa Raw-p
Rapa FDR
ER folding
Hspa5 0.216 0.339 0.728 0.965
Hsp90b1 0.872 0.916 0.847 0.965
P4hb 0.245 0.372 0.994 0.998
Pdia3 0.156 0.282 0.734 0.965
Pdia6 0.626 0.732 0.637 0.965
HSP70/HSP90
Hspa8 0.025 0.132 0.442 0.964
Hspa1l 0.011 0.105 0.356 0.929
Hsp90aa1 0.134 0.263 0.902 0.981
Hsp90ab1 0.029 0.138 0.014 0.432
St13 0.032 0.142 0.270 0.902
Stip1 <0.001 0.085 0.737 0.965
Small HSP
Hspb1 0.829 0.889 0.565 0.965
Hspb2 0.194 0.320 0.800 0.965
Hspb6 0.239 0.367 0.688 0.965
Cryab 0.583 0.691 0.580 0.965
Mito. folding
Hspa9 0.296 0.426 0.784 0.965
Hspd1 0.049 0.170 0.702 0.965
Hspe1 0.404 0.533 0.079 0.721
Z score
2.5
0
-2.5
Treatment
Control
Exercise
Rapamycin
Sex
Female
Male
b
Soluble — Exercise
Eprs1
Meioc
Pfkm
Stip1
Gars1
Psma4
-log10(raw p-value)
log2FC (Ex / Ctl)
NS (327) Down (133) Up (1)
c
Insoluble — Exercise
Col1a2
Tmod1
Hba
Myoz3
Hbb-bs
log2FC (Ex / Ctl)
NS (95) Down (47) Up (0)
d
Ins/Sol — Exercise
Hbb-bs
Hba
Pdlim3
Pgm1
Tpi1
log2FC (Ex / Ctl)
NS (104) Down (6) Up (0)
FDR ≤ 0.05
Raw p ≤ 0.05
e
Soluble — Rapamycin
Obscn
Myom1
Myom2
Ank1
Eif4a2
Mif
Myl1
Sms
Cox7a1
Phpt1
log2FC (Rapa / Ctl)
NS (431) Down (16) Up (14)
f
Insoluble — Rapamycin
Krt5
Ttn
Mybpc2
Tpm3
Mybph
Mybpc1
log2FC (Rapa / Ctl)
NS (126) Down (15) Up (1)
g
Ins/Sol — Rapamycin
Pdlim3
Pgam2
H2bc14
Myl1
Krt5
log2FC (Rapa / Ctl)
NS (98) Down (12) Up (0)
h
Old Ctl
Old+Ex
Old+Rapa
log10(Soluble Abundance)
log10(Total Abundance (Soluble + Insoluble))
Cellular component
Cytoskeleton
Cytosol
ECM
ER
Extracellular
Mitochondrion
Nucleus
Plasma mem.
Myofibril
Other

**Extended Data Fig. 7 | Effects of exercise and rapamycin on protein abundance and solubility in aged skeletal muscle. a,** Heatmap of soluble proteins involved in endoplasmic reticulum folding, HSP70/HSP90 systems, small heat-shock protein responses and mitochondrial folding. Columns represent individual mice, annotated by treatment and sex; values are row Z-scores of $\log_2$ abundance, with the color scale clipped at ±2.5. Raw and Benjamini–Hochberg-adjusted P values are shown for treatment-versus-control contrasts. **b–g,** Effects of exercise (**b–d**) or rapamycin (**e–g**) on soluble abundance (**b,e**), insoluble abundance (**c,f**) and volume-corrected insoluble-to-soluble protein ratios (**d,g**). Each point represents a protein. **h,** Relationship between soluble and total protein abundance, calculated from paired, volume-corrected fractions and averaged within treatment. Points represent proteins, colored by cellular component; dashed diagonals indicate 100% solubility. For both fractions and paired-ratio analyses, n = 5 control female, 4 control male, 4 exercise female, 4 exercise male, 3 rapamycin female and 5 rapamycin male mice. The volcano analyses tested 461 soluble proteins, 142 insoluble proteins and 110 protein ratios; valid observations vary by protein. Protein-level effects were estimated using treatment × sex linear models with intensity-dependent empirical Bayes moderation (limma), averaging female and male treatment effects equally. Tests were two-sided, with Benjamini–Hochberg correction across proteins separately within each contrast and analysis. Volcano plots show FDR-significant proteins (filled colored circles), proteins meeting nominal $P \leq 0.05$ only (open colored circles) and nonsignificant proteins (grey). Red and blue indicate positive and negative effects. Dashed lines indicate raw $P = 0.05$ and $\log_2$ fold change = ±0.5; the fold-change lines are visual guides and do not define significance.

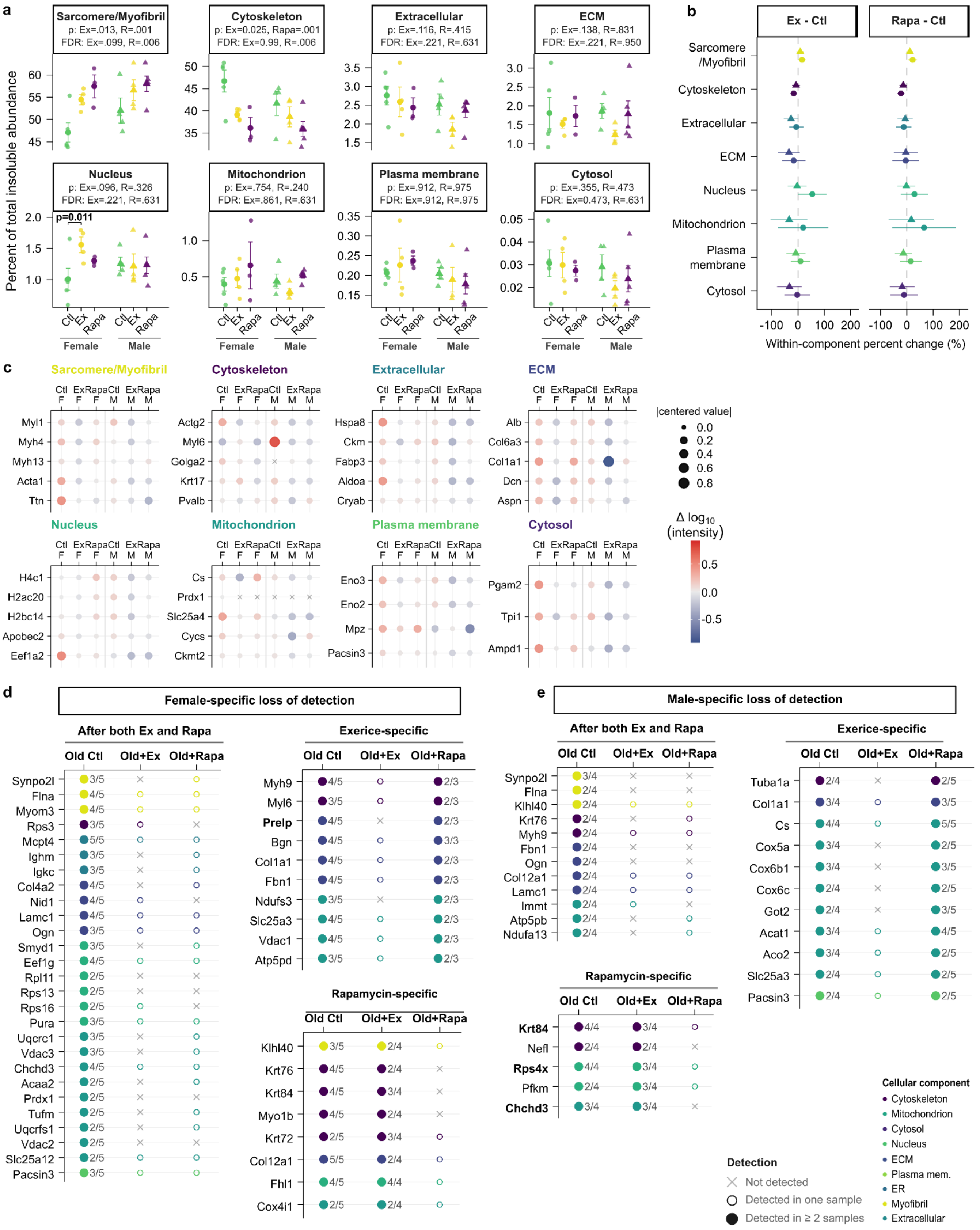
a
Percent of total insoluble abundance
Sarcomere/Myofibril
p: Ex=.013, R=.001
FDR: Ex=.099, R=.006
Cytoskeleton
p: Ex=0.025, Rapa=.001
FDR: Ex=0.99, R=.006
Extracellular
p: Ex=.116, R=.415
FDR: Ex=.221, R=.631
ECM
p: Ex=.138, R=.831
FDR: Ex=.221, R=.950
Nucleus
p: Ex=.096, R=.326
FDR: Ex=.221, R=.631
p=0.011
Mitochondrion
p: Ex=.754, R=.240
FDR: Ex=.861, R=.631
Plasma membrane
p: Ex=.912, R=.975
FDR: Ex=.912, R=.975
Cytosol
p: Ex=.355, R=.473
FDR: Ex=0.473, R=.631
Ctl Ex Rapa Ctl Ex Rapa
Female Male
b
Ex - Ctl
Rapa - Ctl
Sarcomere /Myofibril
Cytoskeleton
Extracellular
ECM
Nucleus
Mitochondrion
Plasma membrane
Cytosol
Within-component percent change (%)
c
Sarcomere/Myofibril
Myl1 Myh4 Myh13 Acta1 Ttn
Cytoskeleton
Actg2 Myl6 Golga2 Krt17 Pvalb
Extracellular
Hspa8 Ckm Fabp3 Aldoa Cryab
ECM
Alb Col6a3 Col1a1 Dcn Aspn
Nucleus
H4c1 H2ac20 H2bc14 Apobec2 Eef1a2
Mitochondrion
Cs Prdx1 Slc25a4 Cycs Ckmt2
Plasma membrane
Eno3 Eno2 Mpz Pacsin3
Cytosol
Pgam2 Tpi1 Ampd1
Ctl F Ex F Rapa F Ctl M Ex M Rapa M
|centered value|
0.0 0.2 0.4 0.6 0.8
Δ log10 (intensity)
0.5 0.0 -0.5
d
Female-specific loss of detection
After both Ex and Rapa
Old Ctl Old+Ex Old+Rapa
Synpo2l Flna Myom3 Rps3 Mcpt4 Ighm Igkc Col4a2 Nid1 Lamc1 Ogn Smyd1 Eef1g Rpl11 Rps13 Rps16 Pura Uqcrc1 Vdac3 Chchd3 Acaa2 Prdx1 Tufm Uqcrfs1 Vdac2 Slc25a12 Pacsin3
Exerice-specific
Myh9 Myl6 Prelp Bgn Col1a1 Fbn1 Ndufs3 Slc25a3 Vdac1 Atp5pd
Rapamycin-specific
Klhl40 Krt76 Krt84 Myo1b Krt72 Col12a1 Fhl1 Cox4i1
e
Male-specific loss of detection
After both Ex and Rapa
Synpo2l Flna Klhl40 Krt76 Myh9 Fbn1 Ogn Col12a1 Lamc1 Immt Atp5pb Ndufa13
Exerice-specific
Tuba1a Col1a1 Cs Cox5a Cox6b1 Cox6c Got2 Acat1 Aco2 Slc25a3 Pacsin3
Rapamycin-specific
Krt84 Nefl Rps4x Pfkm Chchd3
Detection
Not detected
Detected in one sample
Detected in ≥ 2 samples
Cellular component
Cytoskeleton
Mitochondrion
Cytosol
Nucleus
ECM
Plasma mem.
ER
Myofibril
Extracellular

**Extended Data Fig. 8 | Cellular composition and treatment-associated loss of protein detection in the aged insoluble proteome. a,** Contributions of cellular components to insoluble protein abundance in old control (Ctl), exercise (Ex) and rapamycin-treated (Rapa) female and male mice. Percentages use all GO-assigned, filtered proteins as the denominator, including omitted components. **b,** Sex-specific relative changes in component representation, calculated as 100 × (mean treated fraction − mean control fraction)/mean control fraction. **c,** Five proteins with the highest pooled old-control mean abundance per component, where available. Color shows $\log_{10}$(mean intensity among positive detections) centered within protein across the six treatment × sex groups; size shows the absolute centered value, and crosses indicate unavailable values. **d,e,** Detection-loss patterns in female (**d**) and male (**e**) mice. All candidates have ≥2 control detections. Shared loss requires ≤1 detection after each intervention; exercise-specific loss requires ≤1 exercise and ≥2 rapamycin detections; rapamycin-specific loss requires ≥2 exercise and ≤1 rapamycin detections. Symbols distinguish zero, one and ≥2 detections; fractions show samples with detection/total samples. These categories describe detection patterns. For all analyses, n = 5 control female, 4 control male, 4 exercise female, 4 exercise male, 3 rapamycin female and 5 rapamycin male mice. Composition analyses retained proteins detected in ≥2 samples in any treatment group, pooling sexes within treatment. In **a**, symbols represent individual mice and values are means ± s.e.m. Arcsine-square-root-transformed component fractions were analyzed using full six-group models with planned two-sided t-tests of exercise-versus-control and rapamycin-versus-control effects, averaged equally across sexes. Benjamini–Hochberg correction was applied across components separately for each treatment effect and treatment-specific sex interaction. Internal brackets identify nominally significant treatment × sex interactions. Error bars in **b** show model-based 95% confidence intervals calculated using the delta method. Exact P and FDR values are shown where indicated.

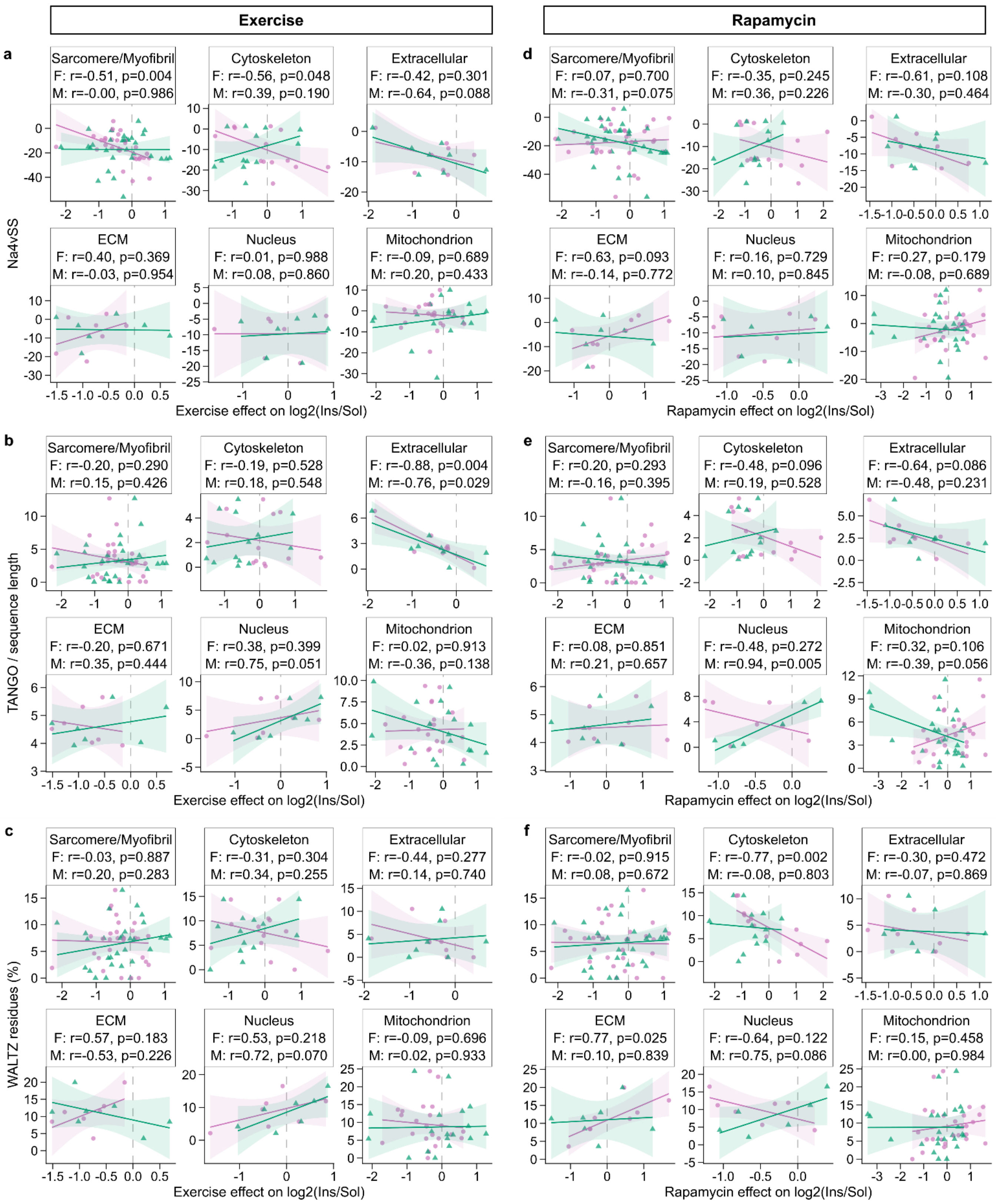

Exercise
Rapamycin
a
Sarcomere/Myofibril
F: r=-0.51, p=0.004
M: r=-0.00, p=0.986
Cytoskeleton
F: r=-0.56, p=0.048
M: r=0.39, p=0.190
Extracellular
F: r=-0.42, p=0.301
M: r=-0.64, p=0.088
ECM
F: r=0.40, p=0.369
M: r=-0.03, p=0.954
Nucleus
F: r=0.01, p=0.988
M: r=0.08, p=0.860
Mitochondrion
F: r=-0.09, p=0.689
M: r=0.20, p=0.433
Na4vSS
Exercise effect on log2(Ins/Sol)
d
Sarcomere/Myofibril
F: r=0.07, p=0.700
M: r=-0.31, p=0.075
Cytoskeleton
F: r=-0.35, p=0.245
M: r=0.36, p=0.226
Extracellular
F: r=-0.61, p=0.108
M: r=-0.30, p=0.464
ECM
F: r=0.63, p=0.093
M: r=-0.14, p=0.772
Nucleus
F: r=0.16, p=0.729
M: r=0.10, p=0.845
Mitochondrion
F: r=0.27, p=0.179
M: r=-0.08, p=0.689
Rapamycin effect on log2(Ins/Sol)
b
Sarcomere/Myofibril
F: r=-0.20, p=0.290
M: r=0.15, p=0.426
Cytoskeleton
F: r=-0.19, p=0.528
M: r=0.18, p=0.548
Extracellular
F: r=-0.88, p=0.004
M: r=-0.76, p=0.029
ECM
F: r=-0.20, p=0.671
M: r=0.35, p=0.444
Nucleus
F: r=0.38, p=0.399
M: r=0.75, p=0.051
Mitochondrion
F: r=0.02, p=0.913
M: r=-0.36, p=0.138
TANGO / sequence length
Exercise effect on log2(Ins/Sol)
e
Sarcomere/Myofibril
F: r=0.20, p=0.293
M: r=-0.16, p=0.395
Cytoskeleton
F: r=-0.48, p=0.096
M: r=0.19, p=0.528
Extracellular
F: r=-0.64, p=0.086
M: r=-0.48, p=0.231
ECM
F: r=0.08, p=0.851
M: r=0.21, p=0.657
Nucleus
F: r=-0.48, p=0.272
M: r=0.94, p=0.005
Mitochondrion
F: r=0.32, p=0.106
M: r=-0.39, p=0.056
Rapamycin effect on log2(Ins/Sol)
c
Sarcomere/Myofibril
F: r=-0.03, p=0.887
M: r=0.20, p=0.283
Cytoskeleton
F: r=-0.31, p=0.304
M: r=0.34, p=0.255
Extracellular
F: r=-0.44, p=0.277
M: r=0.14, p=0.740
ECM
F: r=0.57, p=0.183
M: r=-0.53, p=0.226
Nucleus
F: r=0.53, p=0.218
M: r=0.72, p=0.070
Mitochondrion
F: r=-0.09, p=0.696
M: r=0.02, p=0.933
WALTZ residues (%)
Exercise effect on log2(Ins/Sol)
f
Sarcomere/Myofibril
F: r=-0.02, p=0.915
M: r=0.08, p=0.672
Cytoskeleton
F: r=-0.77, p=0.002
M: r=-0.08, p=0.803
Extracellular
F: r=-0.30, p=0.472
M: r=-0.07, p=0.869
ECM
F: r=0.77, p=0.025
M: r=0.10, p=0.839
Nucleus
F: r=-0.64, p=0.122
M: r=0.75, p=0.086
Mitochondrion
F: r=0.15, p=0.458
M: r=0.00, p=0.984
Rapamycin effect on log2(Ins/Sol)

**Extended Data Fig. 9 | Relationships between intrinsic aggregation propensity and intervention-associated changes in protein partitioning. a–f,** Relationships between treatment effects on the volume-corrected $\log_2$ insoluble-to-soluble protein ratio and AGGRESCAN Na4vSS (**a,d**), sequence-length-normalized TANGO scores (**b,e**) or WALTZ-predicted amyloidogenic residues (%) (**c,f**). Exercise-versus-control effects are shown in **a–c** and rapamycin-versus-control effects in **d–f**. Points represent individual proteins; circles and triangles distinguish females and males, respectively. Solid lines show linear regressions with 95% confidence intervals. The reference pool comprised proteins detected in ≥2 old-control insoluble samples, pooling sexes. Sex-specific effects required ≥2 valid paired soluble/insoluble observations in both control and the corresponding treatment group. The proteomic cohort comprised n = 5 control female, 4 control male, 4 exercise female, 4 exercise male, 3 rapamycin female and 5 rapamycin male mice. Aggregation-propensity scores represent protein-level properties rather than independent biological replicates. Protein numbers depend on the sex-specific paired-detection criteria. Associations were evaluated using two-sided Pearson correlation tests within each treatment, metric, component and sex when ≥5 proteins had nonconstant values. Headers show r and unadjusted P values; F and M denote females and males. Benjamini–Hochberg correction was applied across the component × sex × metric tests within each treatment; adjusted values are retained in the analysis outputs.

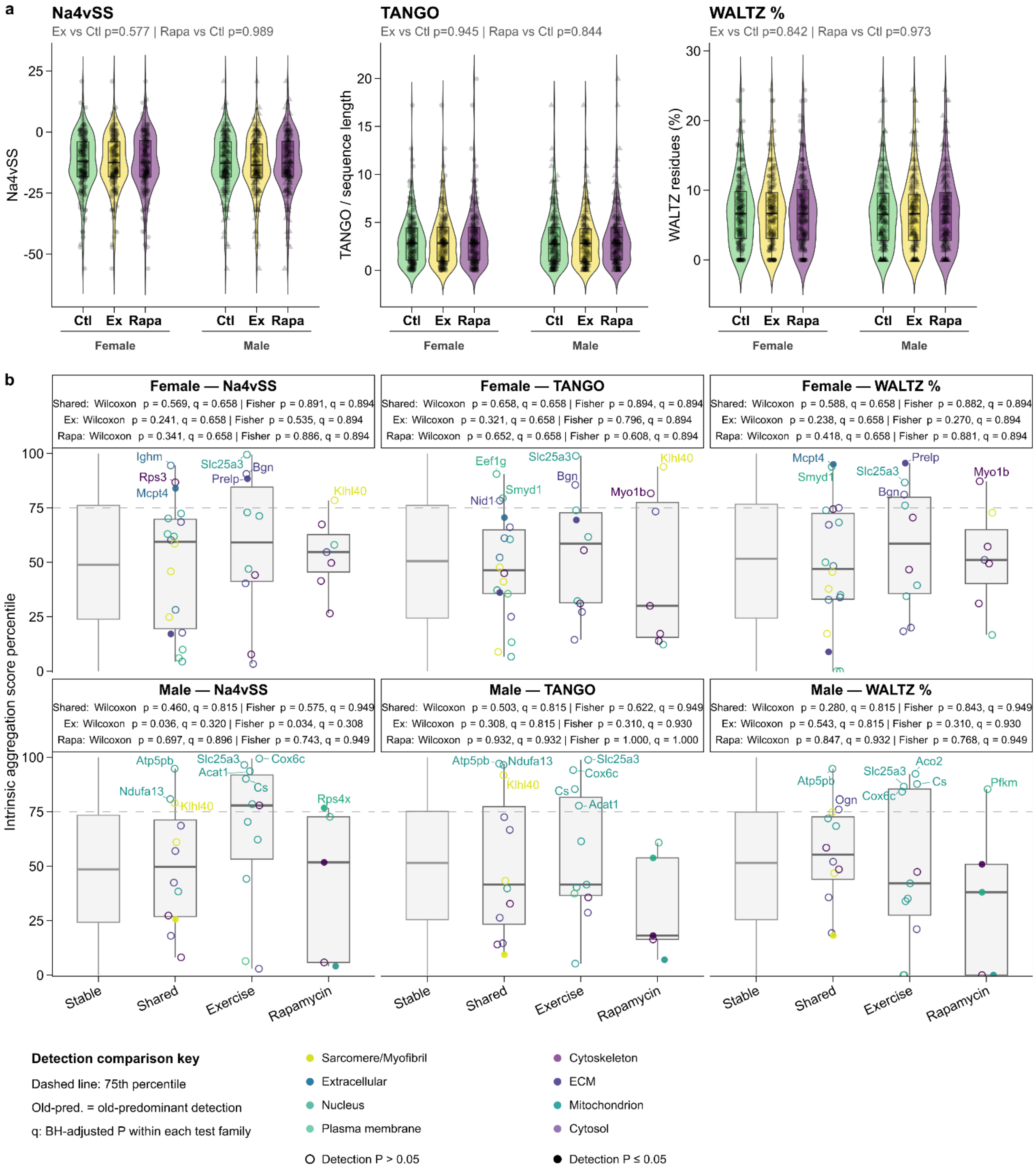


**Extended Data Fig. 10 | Global aggregation propensity and detection-associated protein subsets following exercise or rapamycin treatment. a,** Unweighted distributions of AGGRESCAN Na4vSS, sequence-length-normalized TANGO scores and WALTZ-predicted amyloidogenic residues (%) among insoluble proteins with ≥2 detections within each treatment × sex group. **b,** Score percentiles for stable proteins and proteins showing shared, exercise-specific or rapamycin-specific detection loss, separated by sex. All groups require ≥2 old-control detections. Stable proteins have ≥2 detections after both interventions; shared loss requires ≤1 detection after each; exercise-specific loss requires ≤1 exercise and ≥2 rapamycin detections; rapamycin-specific loss requires ≥2 exercise and ≤1 rapamycin detections. Percentiles reference all scored proteins with ≥2 old-control insoluble detections within sex. Colors indicate cellular components. Filled symbols indicate nominal detection P < 0.05 for the relevant intervention; either intervention meeting this threshold is sufficient for shared loss. Open symbols indicate other candidates. Proteomic measurements were obtained from n = 5 control female, 4 control male, 4 exercise female, 4 exercise male, 3 rapamycin female and 5 rapamycin male mice. Each propensity point represents a protein rather than an independent biological replicate. In **b**, shared/exercise-specific/rapamycin-specific protein counts were 18/10/7 in females and 12/11/5 in males for each metric. Stable-protein counts were 146 females and 144 males for Na4vSS, and 145 females and 143 males for TANGO and WALTZ. Comparisons in **a** used separate treatment × sex factorial models for exercise versus control and rapamycin versus control: Type III HC3-robust linear-model F-tests for Na4vSS and aligned-rank-transform models for TANGO and WALTZ. Displayed treatment P values are unadjusted. In **b**, boxes show medians and interquartile ranges; whiskers extend to the most extreme values within 1.5 times the interquartile range. Each loss category was compared with stable proteins using one-sided Wilcoxon rank-sum tests for higher scores and one-sided Fisher's exact tests for enrichment at or above the 75th percentile. Benjamini–Hochberg correction was applied across nine category × metric comparisons within sex, separately for each test family. Ctl, control; Ex, exercise; Rapa, rapamycin.
