## Supplementary Figures for "Skeletal muscle aging drives sex- and compartment-specific protein insolubility that is modifiable by proteostasis-directed treatments"

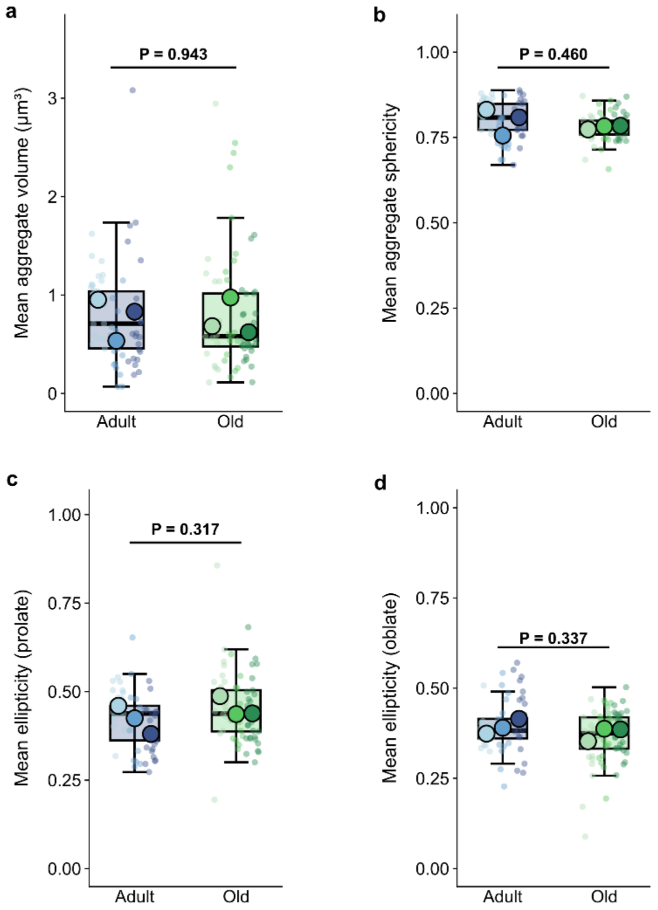


**Supplementary Fig. 1 | Morphological characteristics of amyloid-like protein aggregates are not significantly altered with aging. a–d,** Quantification of amyloid-like aggregate morphology in skeletal muscle from adult and old mice using three-dimensional microscopy. **a,** Mean volume of individual aggregates ($\mu m^3$). **b,** Mean aggregate sphericity. **c,** Mean prolate ellipticity. **d,** Mean oblate ellipticity. Small translucent circles represent individual microscopy images, whereas larger, black-outlined circles represent individual mouse-level estimates (n = 3 mice per age group). Boxplots show the median and interquartile range (25th–75th percentiles) of image-level measurements, with whiskers extending to the most extreme values within 1.5 times the interquartile range. Statistical comparisons were performed using two-sided unpaired Student's t-tests with equal variances, using mice as independent biological replicates. Exact P values are shown above each comparison.

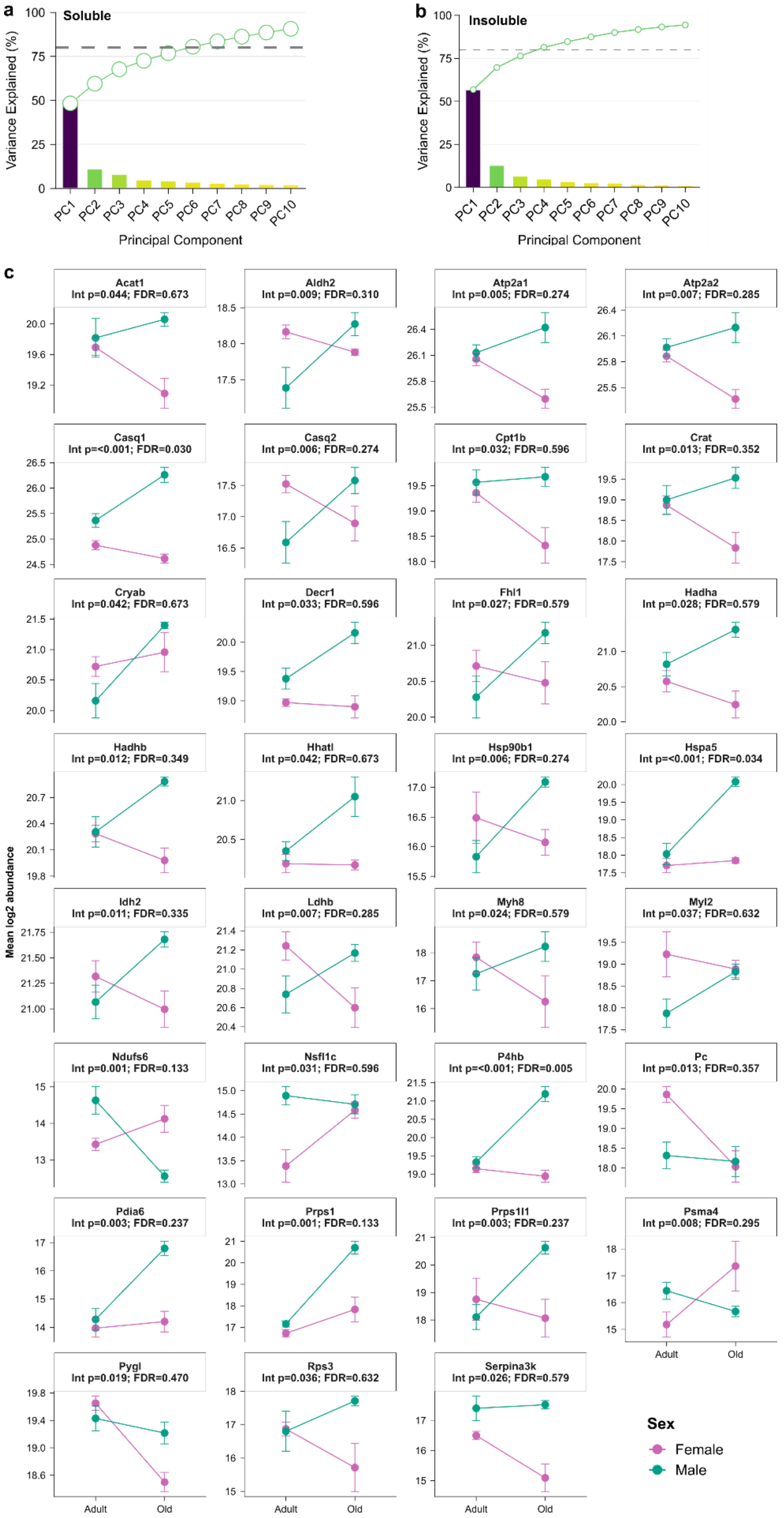
a
Soluble
Variance Explained (%)
Principal Component
PC1 PC2 PC3 PC4 PC5 PC6 PC7 PC8 PC9 PC10
b
Insoluble
Variance Explained (%)
Principal Component
PC1 PC2 PC3 PC4 PC5 PC6 PC7 PC8 PC9 PC10
c
Mean log2 abundance
Acat1
Int p=0.044; FDR=0.673
Aldh2
Int p=0.009; FDR=0.310
Atp2a1
Int p=0.005; FDR=0.274
Atp2a2
Int p=0.007; FDR=0.285
Casq1
Int p=<0.001; FDR=0.030
Casq2
Int p=0.006; FDR=0.274
Cpt1b
Int p=0.032; FDR=0.596
Crat
Int p=0.013; FDR=0.352
Cryab
Int p=0.042; FDR=0.673
Decr1
Int p=0.033; FDR=0.596
Fhl1
Int p=0.027; FDR=0.579
Hadha
Int p=0.028; FDR=0.579
Hadhb
Int p=0.012; FDR=0.349
Hhatl
Int p=0.042; FDR=0.673
Hsp90b1
Int p=0.006; FDR=0.274
Hspa5
Int p=<0.001; FDR=0.034
Idh2
Int p=0.011; FDR=0.335
Ldhb
Int p=0.007; FDR=0.285
Myh8
Int p=0.024; FDR=0.579
Myl2
Int p=0.037; FDR=0.632
Ndufs6
Int p=0.001; FDR=0.133
Nsfl1c
Int p=0.031; FDR=0.596
P4hb
Int p=<0.001; FDR=0.005
Pc
Int p=0.013; FDR=0.357
Pdia6
Int p=0.003; FDR=0.237
Prps1
Int p=0.001; FDR=0.133
Prps1l1
Int p=0.003; FDR=0.237
Psma4
Int p=0.008; FDR=0.295
Pygl
Int p=0.019; FDR=0.470
Rps3
Int p=0.036; FDR=0.632
Serpina3k
Int p=0.026; FDR=0.579
Adult
Old
Sex
Female
Male

**Supplementary Fig. 2 | Principal-component variance and sex-dependent aging responses in the soluble proteome. a,b,** Scree plots for principal-component analysis (PCA) of the soluble (**a**) and insoluble (**b**) proteomes from adult and old gastrocnemius muscle. Bars indicate the variance explained by each of the first ten principal components; lines and open circles indicate cumulative explained variance. The dashed line marks 80% cumulative variance. **c,** Soluble protein abundances for proteins showing nominally significant age × sex interactions ($P \leq 0.05$). Colors distinguish females and males; lines connect group means. For all analyses, n = 4 adult female, 5 adult male, 5 old female and 4 old male mice. PCA used $\log_2$-transformed abundances, with proteins centered and scaled, and the NIPALS algorithm to accommodate missing values without imputation. The filtered datasets comprised 477 soluble and 148 insoluble proteins. Values in **c** are means ± s.e.m., calculated from available measurements for each protein and group. Interaction effects were tested using age × sex linear models with intensity-dependent empirical Bayes moderation (limma) and two-sided moderated t-tests. Benjamini–Hochberg correction was applied across all tested soluble proteins for the interaction contrast. Selection imposed no fold-change cutoff. Headers show raw interaction P values (Int p) and false discovery rate (FDR)-adjusted P values; nominal significance does not imply FDR significance.

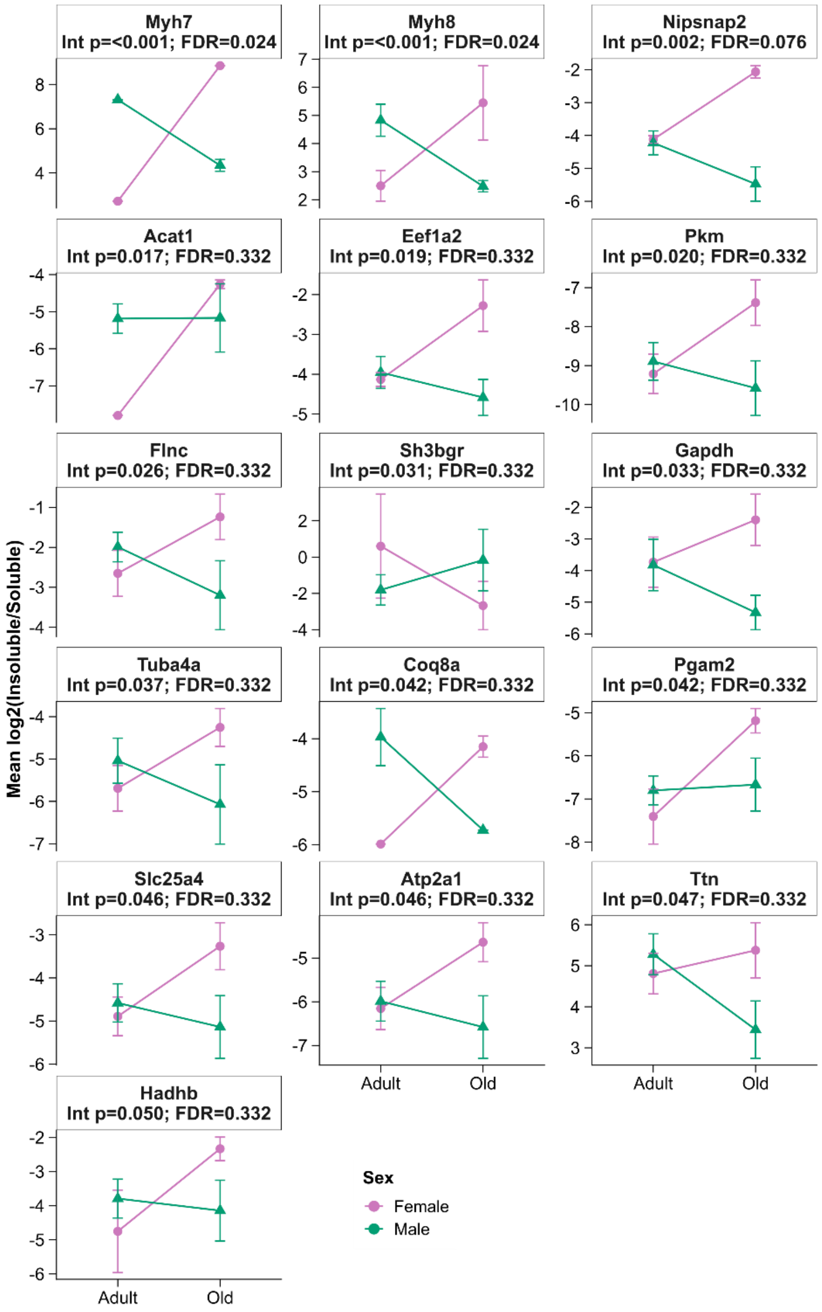


**Supplementary Fig. 3 | Sex-dependent age-associated changes in protein partitioning.** Age-associated changes in the $\log_2$ insoluble-to-soluble protein ratio for proteins showing nominally significant age × sex interactions ($P \le 0.05$). Circles and triangles represent female and male group means, respectively; lines connect the same sex across ages. Soluble and insoluble abundances were matched by protein and animal and corrected for fraction volume and aliquot dilution before calculating ratios. The proteomic cohort comprised n = 4 adult female, 5 adult male, 5 old female and 4 old male mice. Values are means ± s.e.m. of valid paired measurements; available observations vary by protein. Analyses retained 117 proteins with at least two valid paired measurements per age group, pooling sexes for eligibility. Interaction effects were assessed using age × sex linear models with intensity-dependent empirical Bayes moderation (limma) and two-sided moderated t-tests. Benjamini–Hochberg correction was applied across the tested proteins for the interaction contrast. Proteins were selected at nominal interaction $P \le 0.05$ without a fold-change cutoff. Headers show raw interaction P values (Int p) and FDR-adjusted P values.

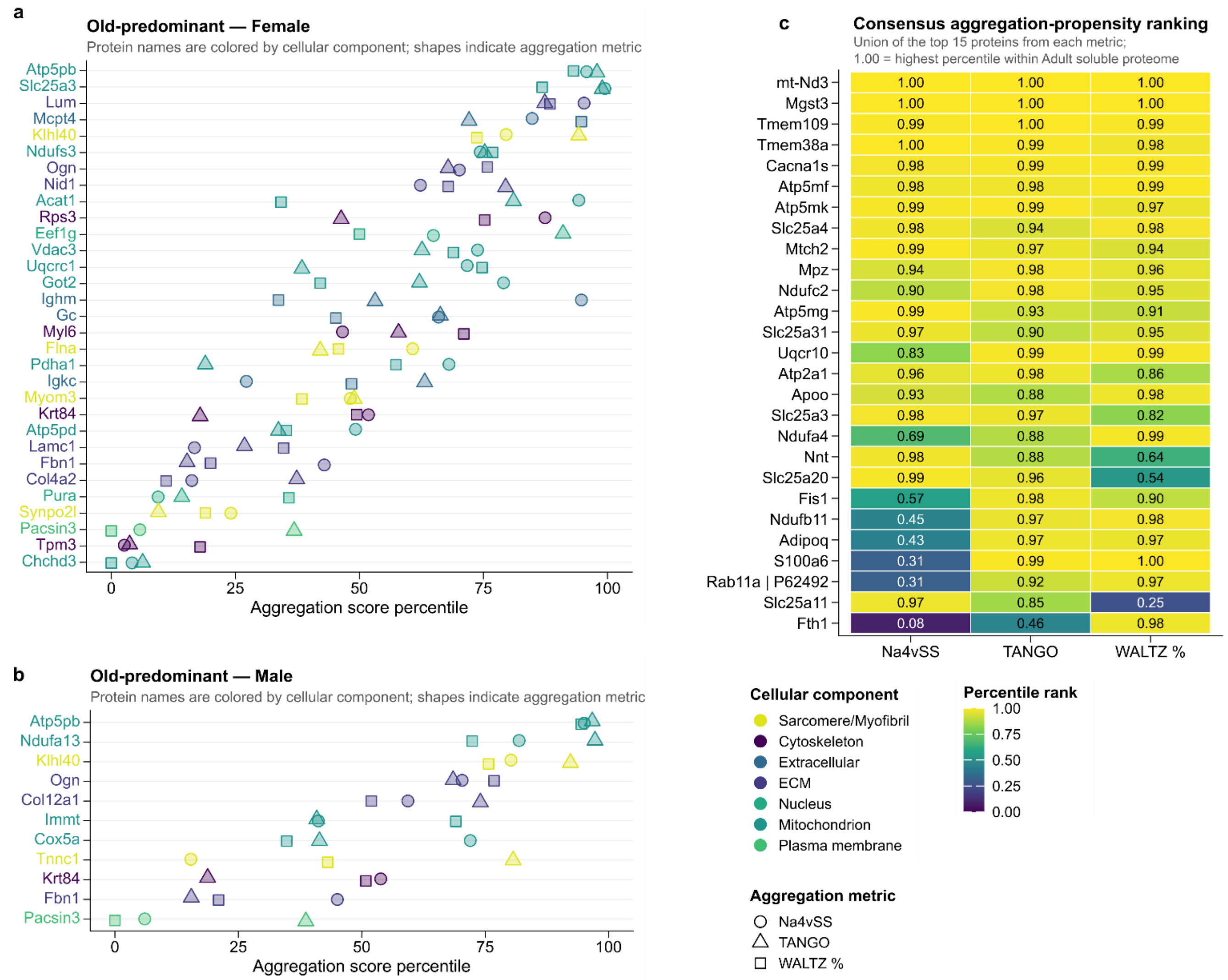


**Supplementary Fig. 4 | Aggregation-propensity profiles of old-predominant proteins and high-scoring adult soluble proteins. a,b,** Intrinsic aggregation-propensity percentiles of old-predominant insoluble proteins in females (**a**) and males (**b**). Old-predominant proteins were detected in ≥2 old and ≤1 adult samples within the same sex. Percentiles are referenced to all scored proteins with ≥2 old insoluble detections within that sex. Circles, triangles and squares indicate AGGRESCAN Na4vSS, sequence-length-normalized TANGO and WALTZ-predicted amyloidogenic residues (%), respectively. Colors of symbols and protein names indicate cellular components. Proteins are ordered by their mean percentile across the three metrics. **c,** Consensus aggregation-propensity ranking of adult soluble proteins, showing the union of the top 15 proteins for each metric. Cell colors and numbers show metric-specific percentile ranks from 0 to 1 within the eligible adult soluble proteome; 1 denotes the highest percentile. Rows are ordered by the mean of the three percentile ranks. The adult-soluble reference pool required detection in ≥2 adult samples, irrespective of sex. Proteomic measurements were obtained from n = 4 adult female, 5 adult male, 5 old female and 4 old male mice. Panels **a** and **b** show 31 and 11 scored proteins, respectively; **c** shows 27 proteins. Scores were derived from protein amino acid sequences and are protein-level properties rather than independent biological replicates. These panels are descriptive; candidates in **a,b** were selected by detection pattern, without requiring a significant detection P value.

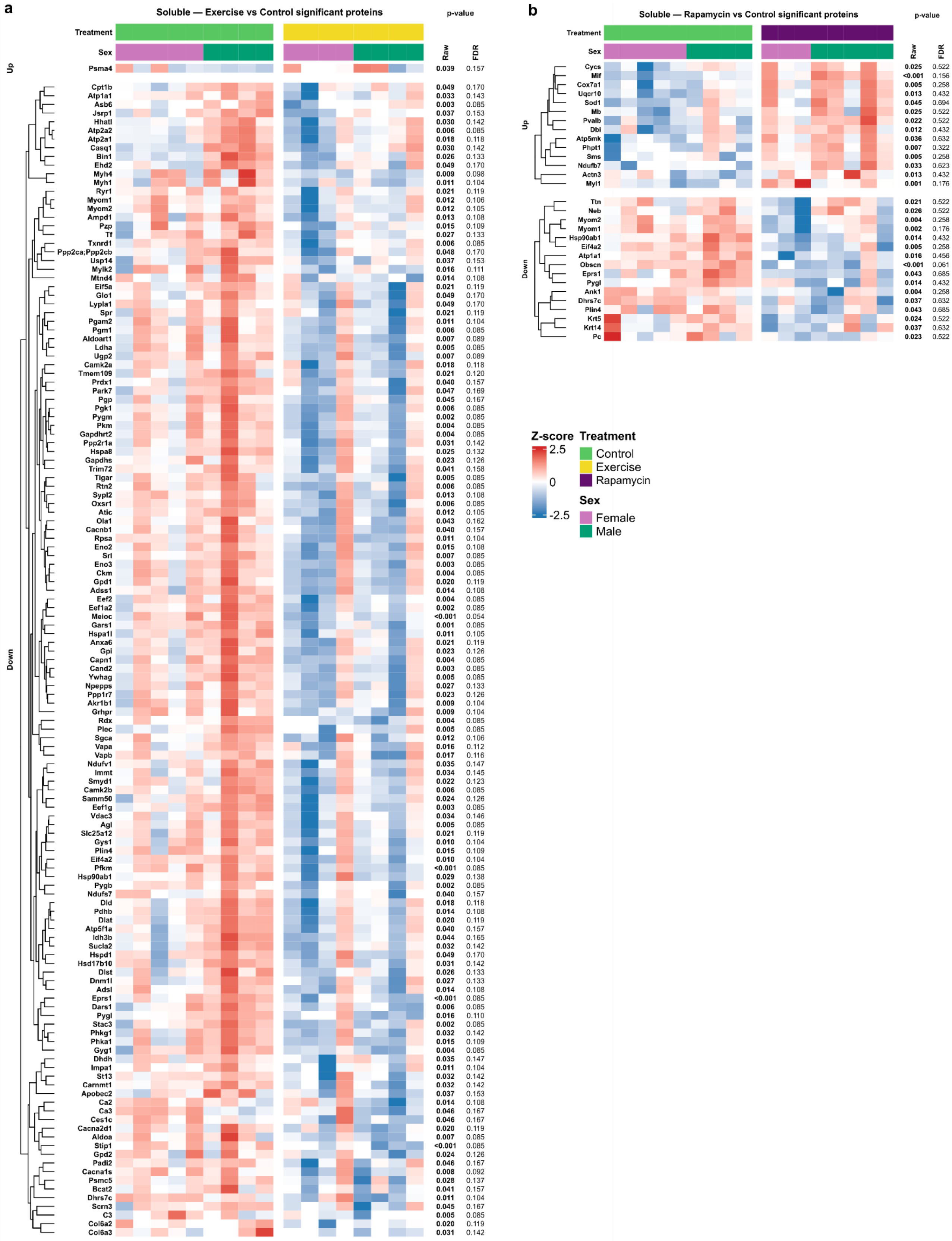
a
Soluble — Exercise vs Control significant proteins
p-value
Treatment
Sex
Raw
FDR
Up
Psma4 0.039 0.157
Down
Cpt1b 0.049 0.170
Atp1a1 0.033 0.143
Asb6 0.003 0.085
Jsrp1 0.037 0.153
Hhatl 0.030 0.142
Atp2a2 0.006 0.085
Atp2a1 0.018 0.118
Casq1 0.030 0.142
Bin1 0.026 0.133
Ehd2 0.049 0.170
Myh4 0.009 0.098
Myh1 0.011 0.104
Ryr1 0.021 0.119
Myom1 0.012 0.106
Myom2 0.012 0.105
Ampd1 0.013 0.108
Pzp 0.015 0.109
Tf 0.027 0.133
Txnrd1 0.006 0.085
Ppp2ca;Ppp2cb 0.048 0.170
Usp14 0.037 0.153
Mylk2 0.016 0.111
Mtnd4 0.014 0.108
Eif5a 0.021 0.119
Glo1 0.049 0.170
Lypla1 0.049 0.170
Spr 0.021 0.119
Pgam2 0.011 0.104
Pgm1 0.006 0.085
Aldoart1 0.007 0.089
Ldha 0.005 0.085
Ugp2 0.007 0.089
Camk2a 0.018 0.118
Tmem109 0.021 0.120
Prdx1 0.040 0.157
Park7 0.047 0.169
Pgp 0.045 0.167
Pgk1 0.006 0.085
Pygm 0.002 0.085
Pkm 0.004 0.085
Gapdhrt2 0.004 0.085
Ppp2r1a 0.031 0.142
Hspa8 0.025 0.132
Gapdhs 0.023 0.126
Trim72 0.041 0.158
Tigar 0.005 0.085
Rtn2 0.006 0.085
Sypl2 0.013 0.108
Oxsr1 0.006 0.085
Atic 0.012 0.105
Ola1 0.043 0.162
Cacnb1 0.040 0.157
Rpsa 0.011 0.104
Eno2 0.015 0.108
Srl 0.007 0.085
Eno3 0.003 0.085
Ckm 0.004 0.085
Gpd1 0.020 0.119
Adss1 0.014 0.108
Eef2 0.004 0.085
Eef1a2 0.002 0.085
Meioc <0.001 0.054
Gars1 0.001 0.085
Hspa1l 0.011 0.105
Anxa6 0.021 0.119
Gpi 0.023 0.126
Capn1 0.004 0.085
Cand2 0.003 0.085
Ywhag 0.005 0.085
Npepps 0.027 0.133
Ppp1r7 0.023 0.126
Akr1b1 0.009 0.104
Grhpr 0.009 0.104
Rdx 0.004 0.085
Plec 0.005 0.085
Sgca 0.012 0.106
Vapa 0.016 0.112
Vapb 0.017 0.116
Ndufv1 0.035 0.147
Immt 0.034 0.145
Smyd1 0.022 0.123
Camk2b 0.006 0.085
Samm50 0.024 0.126
Eef1g 0.003 0.085
Vdac3 0.034 0.146
Agl 0.005 0.085
Slc25a12 0.021 0.119
Gys1 0.010 0.104
Plin4 0.015 0.109
Eif4a2 0.010 0.104
Pfkm <0.001 0.085
Hsp90ab1 0.029 0.138
Pygb 0.002 0.085
Ndufs7 0.040 0.157
Dld 0.018 0.118
Pdhb 0.014 0.108
Dlat 0.020 0.119
Atp5f1a 0.040 0.157
Idh3b 0.044 0.165
Sucla2 0.032 0.142
Hspd1 0.049 0.170
Hsd17b10 0.031 0.142
Dlst 0.026 0.133
Dnm1l 0.027 0.133
Adsl 0.014 0.108
Eprs1 <0.001 0.085
Dars1 0.006 0.085
Pygl 0.016 0.110
Stac3 0.002 0.085
Phkg1 0.032 0.142
Phka1 0.015 0.109
Gyg1 0.004 0.085
Dhdh 0.035 0.147
Impa1 0.011 0.104
St13 0.032 0.142
Carnmt1 0.032 0.142
Apobec2 0.037 0.153
Ca2 0.014 0.108
Ca3 0.046 0.167
Ces1c 0.046 0.167
Cacna2d1 0.020 0.119
Aldoa 0.007 0.085
Stip1 <0.001 0.085
Gpd2 0.024 0.126
Padi2 0.046 0.167
Cacna1s 0.008 0.092
Psmc5 0.028 0.137
Bcat2 0.041 0.157
Dhrs7c 0.011 0.104
Scrn3 0.045 0.167
C3 0.005 0.085
Col6a2 0.020 0.119
Col6a3 0.031 0.142
b
Soluble — Rapamycin vs Control significant proteins
p-value
Treatment
Sex
Raw
FDR
Up
Cycs 0.025 0.522
Mif <0.001 0.156
Cox7a1 0.005 0.258
Uqcr10 0.013 0.432
Sod1 0.045 0.694
Mb 0.025 0.522
Pvalb 0.022 0.522
Dbi 0.012 0.432
Atp5mk 0.036 0.632
Phpt1 0.007 0.322
Sms 0.005 0.258
Ndufb7 0.033 0.623
Actn3 0.013 0.432
Myl1 0.001 0.176
Down
Ttn 0.021 0.522
Neb 0.026 0.522
Myom2 0.004 0.258
Myom1 0.002 0.176
Hsp90ab1 0.014 0.432
Eif4a2 0.005 0.258
Atp1a1 0.016 0.456
Obscn <0.001 0.061
Eprs1 0.043 0.685
Pygl 0.014 0.432
Ank1 0.004 0.258
Dhrs7c 0.037 0.632
Plin4 0.043 0.685
Krt5 0.024 0.522
Krt14 0.037 0.632
Pc 0.023 0.522
Z-score
2.5
0
-2.5
Treatment
Control
Exercise
Rapamycin
Sex
Female
Male

**Supplementary Fig. 5 | Exercise- and rapamycin-associated changes in soluble protein abundance. a,b,** Heatmaps of soluble proteins showing nominally significant abundance changes ($P \leq 0.05$) after exercise (**a**) or rapamycin treatment (**b**) relative to old controls. Columns represent individual mice, annotated by treatment and sex; rows represent proteins, grouped by direction of change and hierarchically clustered within each group. Colors show protein-wise Z-scores of $\log_2$ abundance, calculated using only the samples in the displayed comparison and clipped at ±2.5. Right-hand columns show raw and FDR-adjusted treatment-effect P values. The proteomic cohort comprised n = 5 control female, 4 control male, 4 exercise female, 4 exercise male, 3 rapamycin female and 5 rapamycin male mice. Each panel includes controls and the corresponding intervention group. Analyses tested 461 proteins retained by the 50% detection filter within each treatment group, pooling sexes for eligibility. Treatment effects were estimated using treatment × sex linear models with intensity-dependent empirical Bayes moderation (limma), averaging female and male effects equally. Tests were two-sided, with Benjamini–Hochberg correction across proteins separately for each treatment contrast. Selection imposed no fold-change cutoff. Missing abundances were not imputed for statistical testing; missing heatmap entries were displayed at Z = 0. Nominal selection does not imply FDR significance.

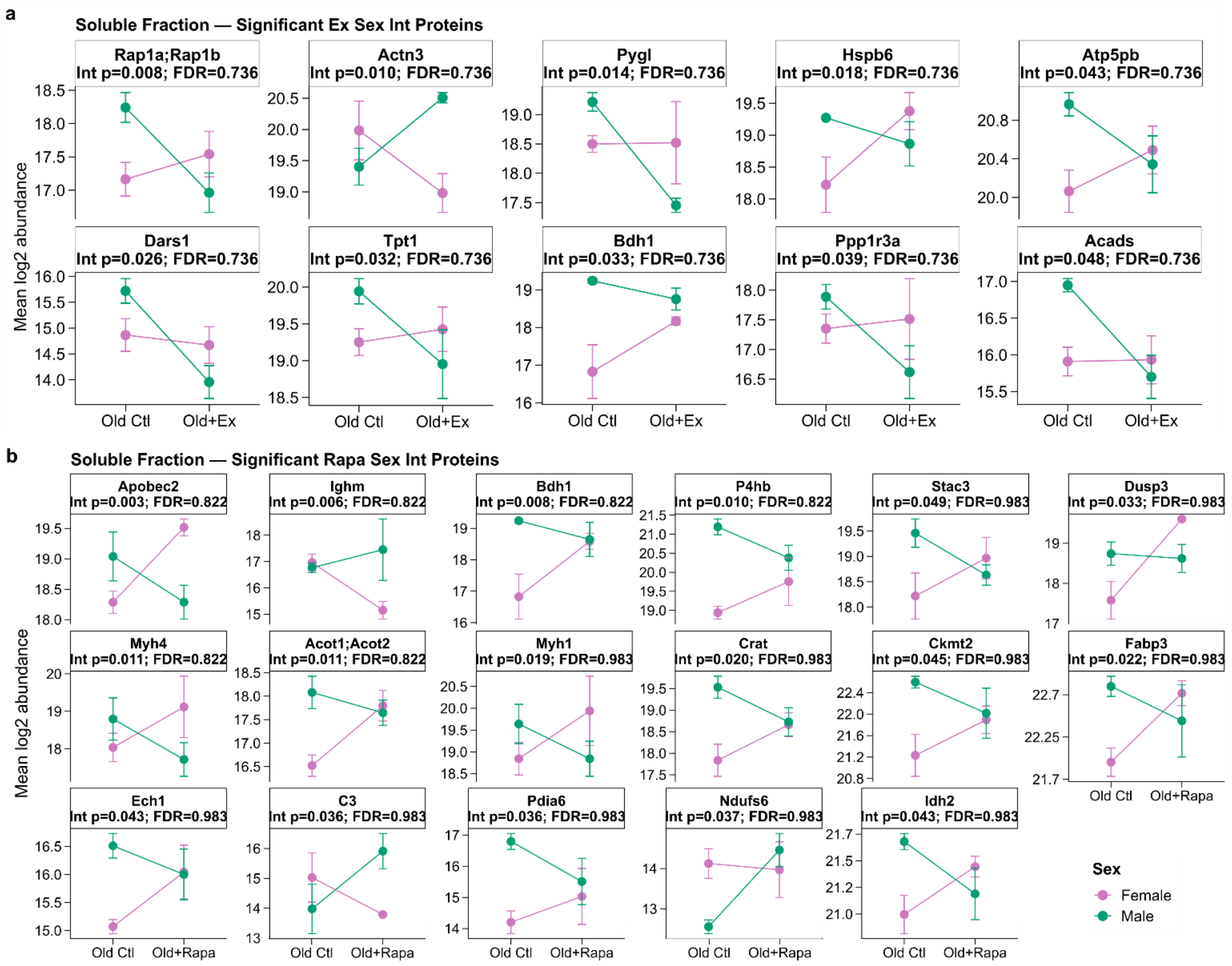


**Supplementary Fig. 6 | Sex-dependent soluble protein responses to exercise and rapamycin. a,b,** Soluble protein abundances for proteins showing nominally significant exercise × sex (**a**) or rapamycin × sex (**b**) interactions ($P \leq 0.05$) in old gastrocnemius muscle. Colors distinguish females and males; lines connect control and intervention group means within sex. The proteomic cohort comprised n = 5 control female, 4 control male, 4 exercise female, 4 exercise male, 3 rapamycin female and 5 rapamycin male mice. Values are means ± s.e.m. calculated from available $\log_2$ abundance measurements; valid observations vary by protein. Interaction effects were estimated using full treatment × sex linear models with intensity-dependent empirical Bayes moderation (limma) and two-sided moderated t-tests. Each interaction tests the difference between female and male treatment responses. Benjamini–Hochberg correction was applied across the 461 tested soluble proteins separately for each interaction contrast. Proteins were selected at nominal interaction $P \leq 0.05$ without a fold-change cutoff. Headers show raw interaction P values (Int p) and FDR-adjusted P values. Ctl, control; Ex, exercise; Rapa, rapamycin.

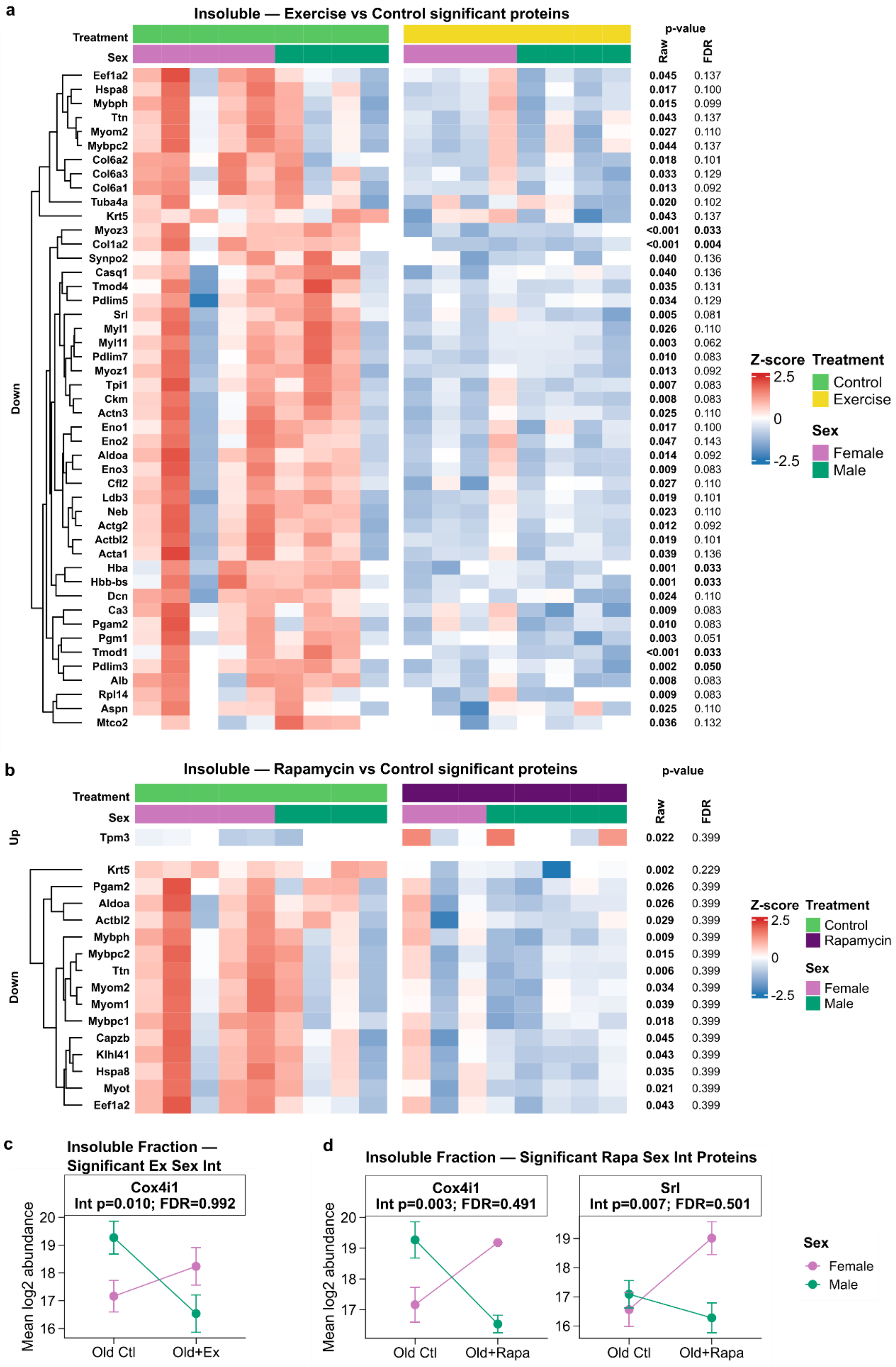

a
Insoluble — Exercise vs Control significant proteins
Treatment
Sex
p-value
Raw
FDR
Down
Eef1a2 0.045 0.137
Hspa8 0.017 0.100
Mybph 0.015 0.099
Ttn 0.043 0.137
Myom2 0.027 0.110
Mybpc2 0.044 0.137
Col6a2 0.018 0.101
Col6a3 0.033 0.129
Col6a1 0.013 0.092
Tuba4a 0.020 0.102
Krt5 0.043 0.137
Myoz3 <0.001 0.033
Col1a2 <0.001 0.004
Synpo2 0.040 0.136
Casq1 0.040 0.136
Tmod4 0.035 0.131
Pdlim5 0.034 0.129
Srl 0.005 0.081
Myl1 0.026 0.110
Myl11 0.003 0.062
Pdlim7 0.010 0.083
Myoz1 0.013 0.092
Tpi1 0.007 0.083
Ckm 0.008 0.083
Actn3 0.025 0.110
Eno1 0.017 0.100
Eno2 0.047 0.143
Aldoa 0.014 0.092
Eno3 0.009 0.083
Cfl2 0.027 0.110
Ldb3 0.019 0.101
Neb 0.023 0.110
Actg2 0.012 0.092
Actbl2 0.019 0.101
Acta1 0.039 0.136
Hba 0.001 0.033
Hbb-bs 0.001 0.033
Dcn 0.024 0.110
Ca3 0.009 0.083
Pgam2 0.010 0.083
Pgm1 0.003 0.051
Tmod1 <0.001 0.033
Pdlim3 0.002 0.050
Alb 0.008 0.083
Rpl14 0.009 0.083
Aspn 0.025 0.110
Mtco2 0.036 0.132
Z-score
2.5
0
-2.5
Treatment
Control
Exercise
Sex
Female
Male
b
Insoluble — Rapamycin vs Control significant proteins
Treatment
Sex
p-value
Raw
FDR
Up
Tpm3 0.022 0.399
Down
Krt5 0.002 0.229
Pgam2 0.026 0.399
Aldoa 0.026 0.399
Actbl2 0.029 0.399
Mybph 0.009 0.399
Mybpc2 0.015 0.399
Ttn 0.006 0.399
Myom2 0.034 0.399
Myom1 0.039 0.399
Mybpc1 0.018 0.399
Capzb 0.045 0.399
Klhl41 0.043 0.399
Hspa8 0.035 0.399
Myot 0.021 0.399
Eef1a2 0.043 0.399
Z-score
2.5
0
-2.5
Treatment
Control
Rapamycin
Sex
Female
Male
c
Insoluble Fraction — Significant Ex Sex Int
Cox4i1
Int p=0.010; FDR=0.992
Mean log2 abundance
20
19
18
17
16
Old Ctl
Old+Ex
d
Insoluble Fraction — Significant Rapa Sex Int Proteins
Cox4i1
Int p=0.003; FDR=0.491
Mean log2 abundance
20
19
18
17
Old Ctl
Old+Rapa
Srl
Int p=0.007; FDR=0.501
19
18
17
16
Old Ctl
Old+Rapa
Sex
Female
Male

**Supplementary Fig. 7 | Treatment-associated changes and sex-dependent responses in the insoluble proteome.** **a,b,** Heatmaps of insoluble proteins showing nominally significant abundance changes ($P \leq 0.05$) after exercise (**a**) or rapamycin treatment (**b**) relative to old controls. Columns represent individual mice, annotated by treatment and sex; proteins are grouped by direction of change and hierarchically clustered within each group. Colors show protein-wise Z-scores of $\log_2$ abundance calculated within each displayed comparison and clipped at ±2.5. Right-hand columns show raw and FDR-adjusted treatment-effect P values. **c,d,** Insoluble protein abundances for proteins showing nominally significant exercise × sex (**c**) or rapamycin × sex (**d**) interactions ($P \leq 0.05$). Colors distinguish sexes; lines connect group means. The proteomic cohort comprised n = 5 control female, 4 control male, 4 exercise female, 4 exercise male, 3 rapamycin female and 5 rapamycin male mice. Each display includes controls and the corresponding intervention group. Values in **c,d** are means ± s.e.m. of available $\log_2$ abundances. Effects were estimated using full treatment × sex linear models with intensity-dependent empirical Bayes moderation (limma). Treatment contrasts in **a,b** average female and male effects equally; interactions in **c,d** test the difference between female and male treatment responses. Tests were two-sided, with Benjamini–Hochberg correction across the 142 tested insoluble proteins separately for each contrast. Selection imposed no fold-change cutoff. Missing abundances were not imputed for testing; missing heatmap entries were displayed at Z = 0. Headers in **c,d** show raw interaction P values (Int p) and FDR-adjusted P values. Ctl, control; Ex, exercise; Rapa, rapamycin.

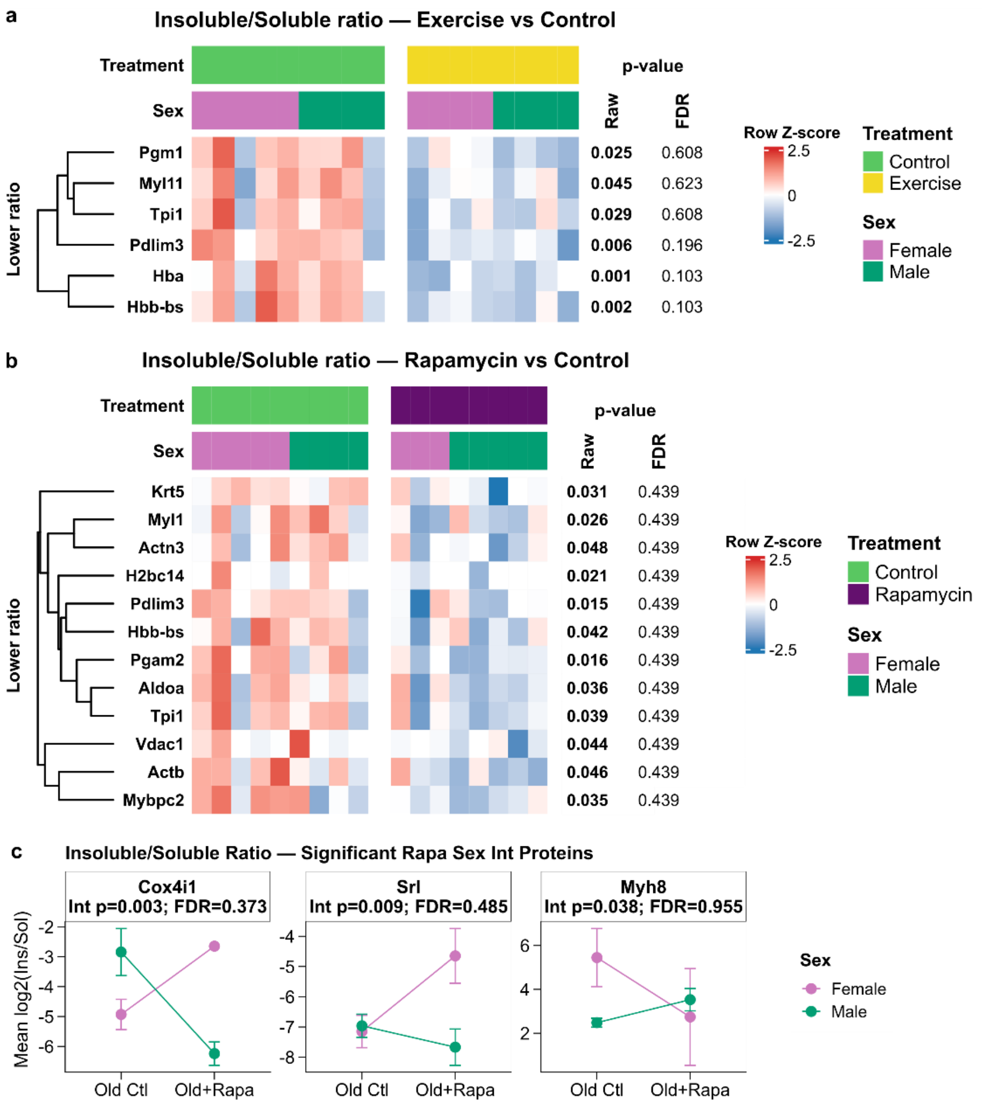


**Supplementary Fig. 8 | Exercise and rapamycin alter the partitioning of selected proteins between soluble and insoluble fractions. a,b,** Heatmaps of proteins showing nominally significant changes ($P \leq 0.05$) in the insoluble-to-soluble protein ratio after exercise (**a**) or rapamycin treatment (**b**) relative to old controls. All displayed treatment effects indicate lower ratios. Columns represent individual mice, annotated by treatment and sex; rows are hierarchically clustered proteins. Colors show protein-wise Z-scores of $\log_2$ ratios, calculated within the displayed comparison and clipped at ±2.5. Right-hand columns show raw and FDR-adjusted treatment-effect P values. **c,** Ratios for proteins showing nominally significant rapamycin × sex interactions ($P \leq 0.05$). Colors distinguish sexes; lines connect group means. Soluble and insoluble abundances were matched by protein and animal and corrected for fraction volume and aliquot dilution before calculating ratios. The proteomic cohort comprised n = 5 control female, 4 control male, 4 exercise female, 4 exercise male, 3 rapamycin female and 5 rapamycin male mice. Each display includes controls and the corresponding intervention group. Values in **c** are means ± s.e.m. of valid paired $\log_2$ ratios. Analyses tested 110 proteins with ≥2 valid paired measurements in each treatment group, pooling sexes for eligibility. Effects were estimated using treatment × sex linear models with intensity-dependent empirical Bayes moderation (limma). Treatment effects average females and males equally; interactions test sex differences in treatment responses. Tests were two-sided, with Benjamini–Hochberg correction across proteins separately for each contrast. Selection imposed no fold-change cutoff. Missing ratios were not imputed for testing; missing heatmap entries were displayed at Z = 0. Headers in **c** show raw interaction P values (Int p) and FDR-adjusted P values. Ctl, control; Ex, exercise; Rapa, rapamycin.

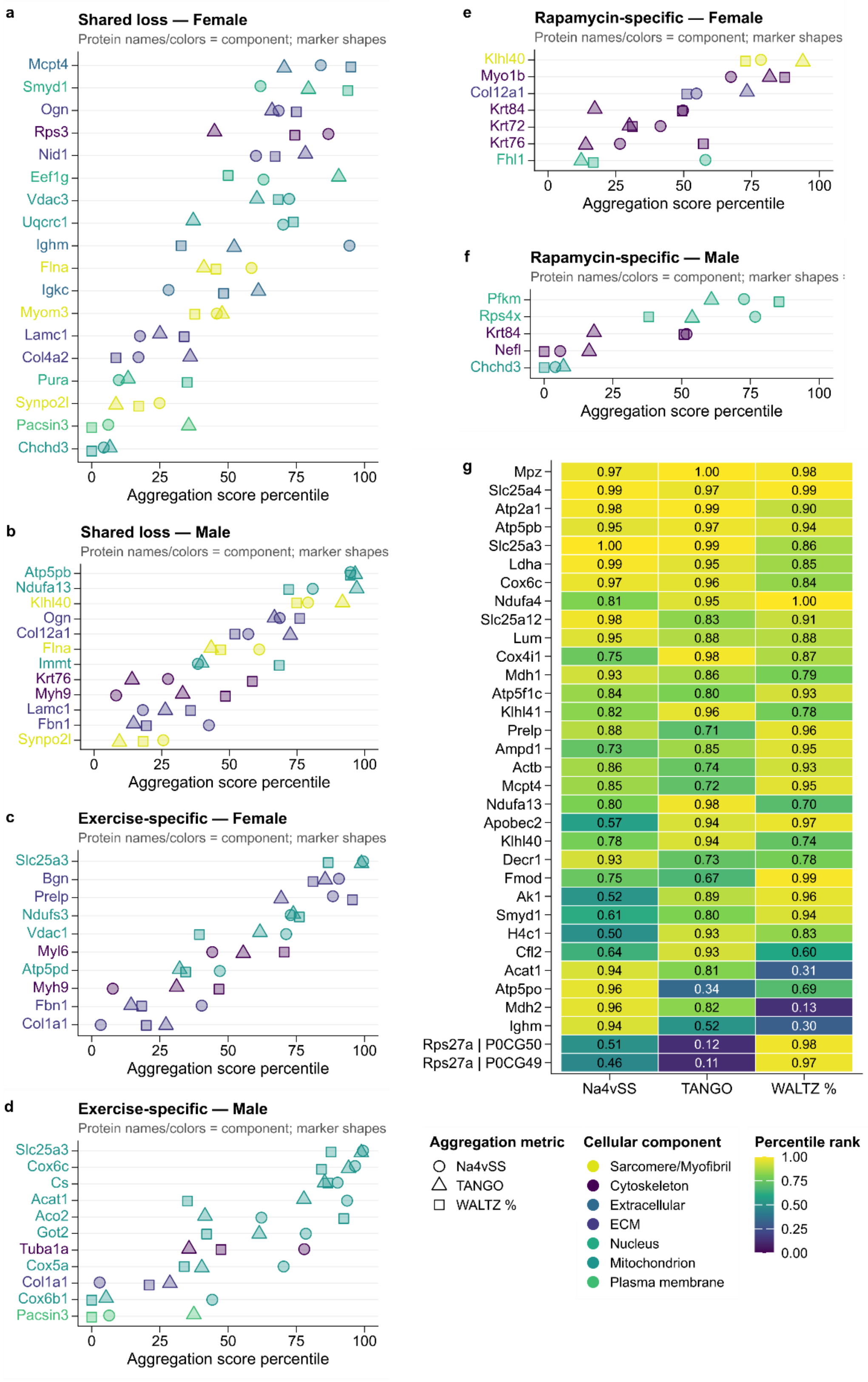
a
Shared loss — Female
Protein names/colors = component; marker shapes
Mcpt4
Smyd1
Ogn
Rps3
Nid1
Eef1g
Vdac3
Uqcrc1
Ighm
Flna
Igkc
Myom3
Lamc1
Col4a2
Pura
Synpo2l
Pacsin3
Chchd3
0 25 50 75 100
Aggregation score percentile
b
Shared loss — Male
Protein names/colors = component; marker shapes
Atp5pb
Ndufa13
Klhl40
Ogn
Col12a1
Flna
Immt
Krt76
Myh9
Lamc1
Fbn1
Synpo2l
0 25 50 75 100
Aggregation score percentile
c
Exercise-specific — Female
Protein names/colors = component; marker shapes
Slc25a3
Bgn
Prelp
Ndufs3
Vdac1
Myl6
Atp5pd
Myh9
Fbn1
Col1a1
0 25 50 75 100
Aggregation score percentile
d
Exercise-specific — Male
Protein names/colors = component; marker shapes
Slc25a3
Cox6c
Cs
Acat1
Aco2
Got2
Tuba1a
Cox5a
Col1a1
Cox6b1
Pacsin3
0 25 50 75 100
Aggregation score percentile
e
Rapamycin-specific — Female
Protein names/colors = component; marker shapes
Klhl40
Myo1b
Col12a1
Krt84
Krt72
Krt76
Fhl1
0 25 50 75 100
Aggregation score percentile
f
Rapamycin-specific — Male
Protein names/colors = component; marker shapes
Pfkm
Rps4x
Krt84
Nefl
Chchd3
0 25 50 75 100
Aggregation score percentile
g
Na4vSS TANGO WALTZ %
Mpz 0.97 1.00 0.98
Slc25a4 0.99 0.97 0.99
Atp2a1 0.98 0.99 0.90
Atp5pb 0.95 0.97 0.94
Slc25a3 1.00 0.99 0.86
Ldha 0.99 0.95 0.85
Cox6c 0.97 0.96 0.84
Ndufa4 0.81 0.95 1.00
Slc25a12 0.98 0.83 0.91
Lum 0.95 0.88 0.88
Cox4i1 0.75 0.98 0.87
Mdh1 0.93 0.86 0.79
Atp5f1c 0.84 0.80 0.93
Klhl41 0.82 0.96 0.78
Prelp 0.88 0.71 0.96
Ampd1 0.73 0.85 0.95
Actb 0.86 0.74 0.93
Mcpt4 0.85 0.72 0.95
Ndufa13 0.80 0.98 0.70
Apobec2 0.57 0.94 0.97
Klhl40 0.78 0.94 0.74
Decr1 0.93 0.73 0.78
Fmod 0.75 0.67 0.99
Ak1 0.52 0.89 0.96
Smyd1 0.61 0.80 0.94
H4c1 0.50 0.93 0.83
Cfl2 0.64 0.93 0.60
Acat1 0.94 0.81 0.31
Atp5po 0.96 0.34 0.69
Mdh2 0.96 0.82 0.13
Ighm 0.94 0.52 0.30
Rps27a | P0CG50 0.51 0.12 0.98
Rps27a | P0CG49 0.46 0.11 0.97
Aggregation metric
Na4vSS
TANGO
WALTZ %
Cellular component
Sarcomere/Myofibril
Cytoskeleton
Extracellular
ECM
Nucleus
Mitochondrion
Plasma membrane
Percentile rank
1.00
0.75
0.50
0.25
0.00

**Supplementary Fig. 9 | Aggregation-propensity profiles of proteins with treatment-associated detection loss and high-scoring old-control insoluble proteins. a–f,** Intrinsic aggregation-propensity percentiles of insoluble proteins showing shared detection loss in females (**a**) and males (**b**), exercise-specific loss in females (**c**) and males (**d**), or rapamycin-specific loss in females (**e**) and males (**f**). All candidates have ≥2 old-control detections within sex. Shared loss requires ≤1 detection after each intervention; exercise-specific loss requires ≤1 exercise and ≥2 rapamycin detections; rapamycin-specific loss requires ≥2 exercise and ≤1 rapamycin detections. Percentiles reference all scored proteins with ≥2 old-control insoluble detections within the same sex. Circles, triangles and squares indicate AGGRESCAN Na4vSS, sequence-length-normalized TANGO and WALTZ-predicted amyloidogenic residues (%), respectively. Protein-name colors indicate cellular components; proteins are ordered by their mean percentile across metrics. **g,** Consensus ranking of high-scoring old-control insoluble proteins, comprising the union of the top 15 proteins for each metric. Cell colors and numbers show metric-specific percentile ranks from 0 to 1 within the eligible old-control insoluble pool; 1 denotes the highest percentile. Rows are ordered by mean percentile across the three metrics. The reference pool for **g** required ≥2 old-control insoluble detections, irrespective of sex. Proteomic measurements were obtained from n = 5 control female, 4 control male, 4 exercise female, 4 exercise male, 3 rapamycin female and 5 rapamycin male mice. Panels **a–f** show 18, 12, 10, 11, 7 and 5 scored proteins, respectively; **g** shows 33 proteins. Scores are sequence-derived protein properties rather than independent biological replicates. These panels are descriptive; loss categories are based on detection counts and do not require significant detection P values.

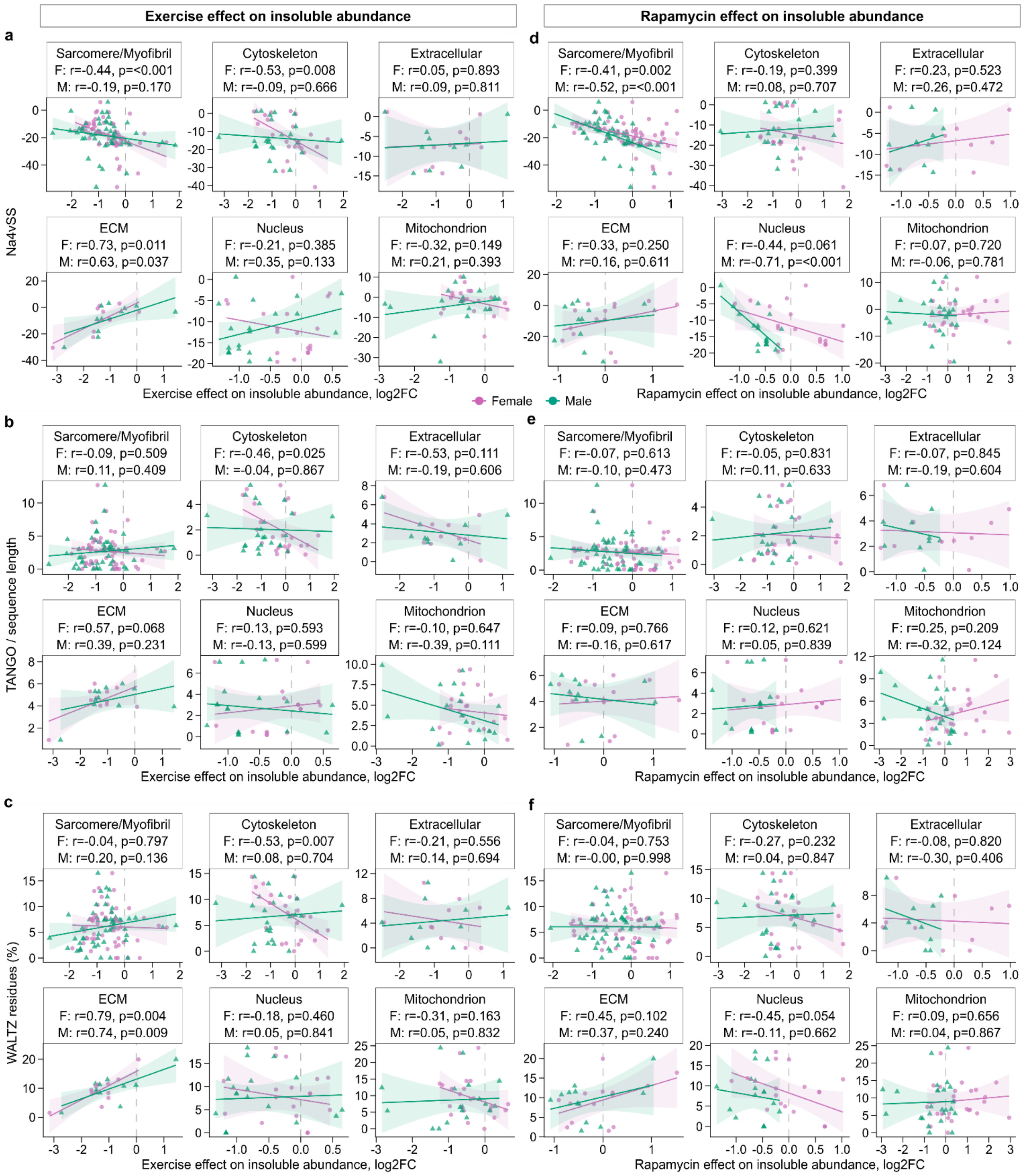
Exercise effect on insoluble abundance
Rapamycin effect on insoluble abundance
a
Sarcomere/Myofibril
F: r=-0.44, p=<0.001
M: r=-0.19, p=0.170
Cytoskeleton
F: r=-0.53, p=0.008
M: r=-0.09, p=0.666
Extracellular
F: r=0.05, p=0.893
M: r=0.09, p=0.811
ECM
F: r=0.73, p=0.011
M: r=0.63, p=0.037
Nucleus
F: r=-0.21, p=0.385
M: r=0.35, p=0.133
Mitochondrion
F: r=-0.32, p=0.149
M: r=0.21, p=0.393
Na4vSS
Exercise effect on insoluble abundance, log2FC
d
Sarcomere/Myofibril
F: r=-0.41, p=0.002
M: r=-0.52, p=<0.001
Cytoskeleton
F: r=-0.19, p=0.399
M: r=0.08, p=0.707
Extracellular
F: r=0.23, p=0.523
M: r=0.26, p=0.472
ECM
F: r=0.33, p=0.250
M: r=0.16, p=0.611
Nucleus
F: r=-0.44, p=0.061
M: r=-0.71, p=<0.001
Mitochondrion
F: r=0.07, p=0.720
M: r=-0.06, p=0.781
Rapamycin effect on insoluble abundance, log2FC
Female
Male
b
Sarcomere/Myofibril
F: r=-0.09, p=0.509
M: r=0.11, p=0.409
Cytoskeleton
F: r=-0.46, p=0.025
M: =-0.04, p=0.867
Extracellular
F: r=-0.53, p=0.111
M: r=-0.19, p=0.606
ECM
F: r=0.57, p=0.068
M: r=0.39, p=0.231
Nucleus
F: r=0.13, p=0.593
M: r=-0.13, p=0.599
Mitochondrion
F: r=-0.10, p=0.647
M: r=-0.39, p=0.111
TANGO / sequence length
Exercise effect on insoluble abundance, log2FC
e
Sarcomere/Myofibril
F: r=-0.07, p=0.613
M: r=-0.10, p=0.473
Cytoskeleton
F: r=-0.05, p=0.831
M: r=0.11, p=0.633
Extracellular
F: r=-0.07, p=0.845
M: r=-0.19, p=0.604
ECM
F: r=0.09, p=0.766
M: r=-0.16, p=0.617
Nucleus
F: r=0.12, p=0.621
M: r=0.05, p=0.839
Mitochondrion
F: r=0.25, p=0.209
M: r=-0.32, p=0.124
Rapamycin effect on insoluble abundance, log2FC
c
Sarcomere/Myofibril
F: r=-0.04, p=0.797
M: r=0.20, p=0.136
Cytoskeleton
F: r=-0.53, p=0.007
M: r=0.08, p=0.704
Extracellular
F: r=-0.21, p=0.556
M: r=0.14, p=0.694
ECM
F: r=0.79, p=0.004
M: r=0.74, p=0.009
Nucleus
F: r=-0.18, p=0.460
M: r=0.05, p=0.841
Mitochondrion
F: r=-0.31, p=0.163
M: r=0.05, p=0.832
WALTZ residues (%)
Exercise effect on insoluble abundance, log2FC
f
Sarcomere/Myofibril
F: r=-0.04, p=0.753
M: r=-0.00, p=0.998
Cytoskeleton
F: r=-0.27, p=0.232
M: r=0.04, p=0.847
Extracellular
F: r=-0.08, p=0.820
M: r=-0.30, p=0.406
ECM
F: r=0.45, p=0.102
M: r=0.37, p=0.240
Nucleus
F: r=-0.45, p=0.054
M: r=-0.11, p=0.662
Mitochondrion
F: r=0.09, p=0.656
M: r=0.04, p=0.867
Rapamycin effect on insoluble abundance, log2FC

**Supplementary Fig. 10 | Component-specific associations between intrinsic aggregation propensity and intervention-associated changes in insoluble protein abundance. a–f,** Relationships between exercise-associated (**a–c**) or rapamycin-associated (**d–f**) changes in insoluble protein abundance and AGGRESCAN Na4vSS (**a,d**), TANGO scores normalized to sequence length (**b,e**) or WALTZ-predicted amyloidogenic residues (%) (**c,f**), analyzed within cellular components. Each point represents a protein; circles and triangles indicate females and males, respectively. Solid lines show linear regressions with shaded 95% confidence intervals. Dashed vertical lines indicate no treatment-associated change in abundance. The reference pool comprised proteins detected in ≥2 old-control insoluble samples, irrespective of sex. Sex-specific treatment effects were calculated as $\log_2$(mean treated/mean control) using positive abundance measurements, requiring ≥2 valid measurements in each comparison group. Proteomic measurements were obtained from n = 5 control female, 4 control male, 4 exercise female, 4 exercise male, 3 rapamycin female and 5 rapamycin male mice. Aggregation-propensity scores represent protein-level properties rather than independent biological replicates. Associations were evaluated using two-sided Pearson correlation tests when ≥5 proteins with nonconstant values were available. Benjamini–Hochberg correction was applied across component-, sex- and metric-specific tests separately within each intervention. Headers show correlation coefficients (r) and unadjusted P values; F and M denote females and males. Protein numbers vary with component, sex and score availability.
